# Cohesin distribution alone predicts chromatin organization in yeast via conserved-current loop extrusion

**DOI:** 10.1101/2023.10.05.560890

**Authors:** Tianyu Yuan, Hao Yan, Kevin C. Li, Ivan Surovtsev, Megan C. King, Simon G. J. Mochrie

**Affiliations:** Integrated Graduate Program in Physical and Engineering Biology, Yale University, New Haven, Connecticut 06520, USA; Department of Physics, Yale University, New Haven, Connecticut 06520, USA; Department of Cell Biology, Yale School of Medicine, New Haven, Connecticut 06520, USA; Department of Molecular, Cell and Developmental Biology, Yale University, New Haven, Connecticut 06511, USA; Department of Applied Physics, Yale University, New Haven, Connecticut 06520, USA

## Abstract

Inhomogeneous patterns of enhanced chromatin-chromatin contacts within 10-100 kb-sized regions of the genome are a generic feature of chromatin spatial organization. These features, termed topologically associating domains (TADs), have led to the loop extrusion factor (LEF) model, where TADs arise from loop extrusion by cohesin complexes. Currently, our ability to model TADs relies on the observation that in vertebrates TAD boundaries are correlated with DNA sequences that bind CTCF, which therefore is inferred to block loop extrusion. However, although TADs feature prominently in their Hi-C maps, non-vertebrate eukaryotes either do not express CTCF or show few TAD boundaries that correlate with CTCF sites. In all of these organisms, the counterparts of CTCF remain unknown, frustrating comparisons between Hi-C data and simulations. To extend the LEF model across the tree of life, here, we propose the *conserved-current loop extrusion (CCLE) model* that interprets loop-extruding cohesin as a nearly-conserved probability current. From cohesin ChIP-seq data alone, we thus derive a position-dependent loop extrusion rate, allowing for a modified paradigm for loop extrusion, that goes beyond solely discrete, localized barriers to also include loop extrusion rates that vary more continuously across the genome. To demonstrate its utility in organisms lacking CTCF, we applied the CCLE model to the Hi-C maps of interphase *Schizosaccharomyces pombe*, as well as to those of meiotic and mitotic *Saccharomyces cerevisiae*. In all cases, even though their Hi-C maps appear quite different, the model accurately predicts the TAD-scale Hi-C maps. It follows that loop extrusion by cohesin is indeed the primary mechanism underlying TADs in these systems. CCLE allows us to obtain loop extrusion parameters such as the LEF density and processivity, which compare well to independent estimates. The model also provides new insights into *in vivo* LEF composition and function.

## Introduction

Our knowledge of chromatin architecture has been transformed by sequencing-based chromatin-capture (Hi-C) techniques, which provide quantitative metrics of relative population-averaged contact probability between all pairs of genomic loci [1–10]. Hi-C data, typically presented as a contact map, together with theory and modeling have led to a new understanding of chromatin organization, based on three coexisting mechanisms that operate on largely different length scales: (1) On several-megabase scales, “checkerboard patterns” in Hi-C maps, encompassing distant contacts on the same chromosome and contacts on different chromosomes, have led to a block co-polymer-inspired picture of chromatin compartments, comprised of several types of epigenetically-distinguished heterochromatin and euchromatin, which each exhibits preferential affinity for like-regions [11–24]. (2) At smaller length scales – tens of kilobases – overlapping squares of high contact probability within definite regions of the same chromosome, termed topologically associating domains (TADs), have led to the loop extrusion factor (LEF) model [25–33], in which LEFs first bind to the chromatin polymer, and then initiate ATP-dependent loop extrusion by moving their two anchor points away from each other. Loop extrusion at an anchor stalls when the anchor encounters either another anchor or a so-called boundary element (BE). LEFs can also dissociate from chromatin. These processes collectively give rise to a dynamic steady-state of chromatin loops, in turn leading to TADs in Hi-C maps [26, 27, 30, 34]. (3) At few kilobase scales, high resolution Hi-C experiments reveal patterns of contacts [35, 36] that can be explained based on nucleosomal structures [37–39].

Depletion of cohesin, a member of the structural maintenance of chromosomes (SMC) complex family, leads to the disappearance both of TADs and of their accompanying enhanced contact probabilities across a variety of species, including human [40, 41], fission yeast [6], and budding yeast [42], thus identifying cohesin as the predominant LEF. Bolstering cohesin’s LEF identity, single-molecule experiments show that cohesin possesses ATP-dependent loop extrusion activity on DNA *in vitro* [43–46]. In vertebrates, the locations of TAD boundaries show a strong correlation with binding sites of the DNA-binding protein, CTCF [47,48], while depletion of CTCF, causes loss of most TAD boundaries [41,47]. These observations suggest that CTCF is the most important BE in vertebrates. Indeed, loop extrusion simulations using boundary elements, whose locations are defined by peaks in the CTCF chromatin immunoprecipitation sequencing (ChIP-seq) signal, are able to recapitulate many aspects of experimental vertebrate Hi-C maps [26, 27, 30, 34].

However, although TADs feature prominently in Hi-C maps across the tree of life, many non-vertebrate organisms either do not express CTCF orthologs, including yeasts (*Schizosaccharomyces pombe* and *Saccharomyces cerevisiae* [49]), plants (*Arabidopsis thaliana* [50] and *Oryza sativa* [51]), and *Caenorhabditis elegans* [52], or show only a limited number of TAD boundaries that correlate with CTCF binding sites, as in the case of *Drosophila melanogaster* [53]. In all of these organisms, even if the LEF model is applicable, which remains uncertain, the identities of the boundary elements are unknown, frustrating quantitative comparisons between Hi-C data and simulations.

With the goal of modeling TADs across the tree of life beyond vertebrates, here, we introduce a novel, physics-based version of the LEF model, named the conserved-current loop extrusion (CCLE) model, that should be applicable in any organism, in which loop extrusion is a major driver of TAD formation. Specifically, by interpreting loop-extruding LEFs as a probability current, that is approximately conserved at steady-state, we derive a position-dependent loop extrusion rate, using cohesin ChIP-seq data as input, which we then incorporate into loop extrusion simulations without explicit boundary elements. This model has intuitive appeal in that loop extrusion rates are small at genomic locations with high cohesin ChIP-seq signal, as if the LEFs are blocked there, while the rates are high at positions with low cohesin ChIP-seq signal, because LEFs spend little time in locations where they are not blocked. By design, CCLE is agnostic concerning the identities of BEs and other proteins that interact with cohesin. Indeed, CCLE allows for the boundary element concept to be extended, beyond localized barriers to loop extrusion, to include more widely distributed variations in the loop extrusion rate, that may occur in response to chromatin composition, for example.

A key inspiration for the development of CCLE was Reference [54], which successfully simulates chromatin organization in meiotic *S. cerevisiae*, using a version of the previous vertebrate-focused LEF models, except with cohesin binding sites replacing CTCF binding sites. CCLE improves on this approach in principle, by eliminating both (1) the need to specify how to pick out cohesin binding sites from ChIP-seq data and (2) the need to specify how such binding sites affect loop extrusion, both of which are accomplished in an *ad hoc* fashion in Ref. [54] (and its CTCF-based precursors [26, 27, 30, 34] regarding CTCF binding sites). Limitation (1) ignores the low-contrast cohesin peaks that could represent a weak yet essential blocking effect by barriers to loop extrusion due to more elaborate chromatin composition, while limitation (2) hinders the model’s ability to self-consistently reproduce the cohesin ChIP-seq data, thus rendering the model a weak physical basis. The CCLE model eliminates both limitations in principle.

To focus on the role of loop extrusion and avoid possible ambiguities associated with chromatin compartments, in this paper, we apply our model to meiotic budding yeast, following Ref. [54], mitotic budding yeast, and interphase fission yeast. None of the corresponding Hi-C maps shows a checkerboard pattern, characteristic of chromatin compartments. Using cohesin ChIP-seq data as input, CCLE seeks to describe the measured Hi-C maps of *S. pombe* quantitatively, with just four fitting parameters, namely the LEF processivity in the absence of obstructions, the chromatin persistence length, the linear density of loop-extruding cohesins (*ρ*), and the linear density of cohesive cohesins (*ρ*_*c*_). For meiotic and mitotic *S. cerevisiae*, we use a fifth parameter to empirically account for the increased polymer volume exclusion in the more compact meiotic and mitotic chromosomes. Using this approach, we demonstrate that CCLE achieves excellent experiment-simulation agreement in all three cases on the 10–100 kb scales, despite major differences in their Hi-C features. Thus, CCLE transforms cohesin ChIP-seq data into an ensemble of fluctuating loop configurations that define three-dimensional chromosomal organization. Since CCLE does not incorporate genomic data on nucleosome positioning, it does not describe the kilobase-scale features in high-resolution Hi-C maps that originate from nucleosomes [37–39]. CCLE also provides corresponding values for loop extrusion parameters, such as the LEF density and processivity, in each case.

## Results

### Conserved-current loop extrusion (CCLE) model enables calculation of chromatin loop configurations from genomic distribution of LEF

In order to develop our approach, we envision loop extrusion as giving rise to probability currents of LEF anchors, flowing through chromatin lattice sites. Assuming two-sided loop extrusion, we are led to the following coarse-grained master equations for the probabilities, *R*_*n*_ and *L*_*n*_, that chromatin site *n* is occupied by the right-moving or left-moving anchor of a LEF:

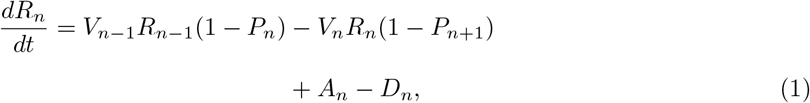

and

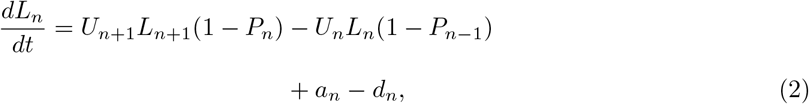

where *V*_*n*_ is the rate at which right-moving LEF anchors step from site *n* to site *n* + 1, *U*_*n*_ is the rate at which left-moving LEF anchors step from site *n* to site *n* − 1, *P*_*n*_ = *R*_*n*_ + *L*_*n*_ is the probability that site *n* is occupied by either a left- or right-moving LEF anchor. The first and second terms on the right

hand sides of each of Eqs. 1 and 2 correspond to the current – *i*.*e*. the number per second – of LEF anchors to and from, respectively, site *n* by loop extrusion along the chromatin, while *A*_*n*_ and *D*_*n*_ (*a*_*n*_ and *d*_*n*_) are the association and dissociation currents of right-moving (left-moving) LEF anchors at site *n*, respectively.

At steady state, assuming the difference between LEF binding and unbinding terms is small compared to the loop-extruding terms, and that the mean probabilities of right- and left-moving LEF anchors being at site *n* are equal (i.e., 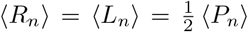), and neglecting correlations among the anchor probabilities, Eqs. 1 and 2 lead to (Supplementary Methods)

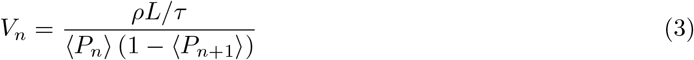

and

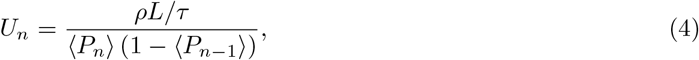

where *ρ* is the mean LEF density in units of kb^−1^, *L* is the mean LEF processivity in units of kb, and *τ* is the mean LEF lifetime, estimated to be of order 10^3^ s in Ref. [31]. The ratio, *L/τ* , can be interpreted as the loop extrusion rate of an isolated LEF along the chromatin polymer, *i*.*e*. twice the extrusion rate of an isolated LEF anchor.

The utility of Eqs. 3 and 4 becomes apparent, when we realize that cohesin ChIP-seq data specifies the *n*-dependence of ⟨*P*_*n*_⟩, assuming that cohesin is the LEF in question. Thus, we can use cohesin ChIP-seq to calculate the position-dependent loop extrusion rates. These rates are subsequently used as input for Gillespie-type loop extrusion simulations (Methods), that implement loop extrusion, LEF association and dissociation, and mutual LEF blocking, as for the existing LEF models, but for which there are no explicit boundary elements. These simulations generate time-dependent loop configurations, that we average over time and over multiple independent simulations to calculate contact probability maps (Methods). Our approach neatly sidesteps needing to know the identities and positions of BEs by exploiting the fact that the effect of BEs is encoded in ⟨*P*_*n*_⟩ and, therefore, is incorporated into the loop extrusion rates via Eqs. 3 and 4.

Functionally, there are two populations of cohesin complexes, namely trans-acting, cohesive cohesins, which give rise to cohesion between sister chromatids, and cis-acting, loop-extruding cohesins [32]. For *S. pombe* both types are present in interphase, and contribute to cohesin ChIP-seq. To determine *P*_*n*_, which is the probability that chromatin site *n* is occupied by loop-extruding cohesin, we assume that cohesive cohesin is randomly loaded along chromatin, giving rise to an *n*-independent contribution to the cohesin ChIP-seq, which we describe with a fitting parameter, *ρ*_*c*_, that represents the uniform density of cohesive cohesin.

Finally, to compare experimental and simulated Hi-C maps, we incorporate the polymer physics of self-contacts, within the confined volume of the nucleus, into simulated loop configurations, using a simple, albeit approximate, analytic approach, described in Methods.

### CCLE Quantitatively Describes TADs and Loop Configurations in Interphase *S. pombe*

#### CCLE simulations accurately reproduce experimental interphase *S. pombe* Hi-C maps

As a first application of CCLE, we sought to describe the interphase fission yeast Hi-C map from Ref. [6], on the basis of ChIP-seq data of the protein, Psc3, which is a component of the cohesin core complex, also from Ref. [6]. The right-hand side of Fig. 1*A* depicts a 1 Mb portion of the Hi-C map of *S. pombe*’s Chr 2 starting 0.3 Mb from the end of the left arm and extending to 1.3 Mb from the end of the left arm, using a logarithmic false-color intensity scale. The maximum interaction distance shown is 120 kb. Each pixel in the experimental Hi-C maps corresponds to 10 kb. The original resolution of the simulation is 1 kb, which is binned to 10 kb to match the experimental resolution. In comparison, the left-hand side of Fig. 1*A* presents the corresponding conserved-current simulated Hi-C map using the best-fit parameters (Table 2). Figure 1*B* magnifies the experiment-simulation comparison for three representative subregions, each 150 kb in size. In both Figs. 1*A* and *B*, visual inspection immediately reveals a high degree of left-right symmetry, corresponding to excellent agreement between the experimental and simulated Hi-C maps. Clearly, our simulations well reproduce the experimentally observed pattern of overlapping squares. By contrast, although the left- and right-hand sides of Figs. 1*C* and *D* look generally similar, clearly there is no left-right symmetry in these figures, which compare the experimental Hi-C maps of two non-overlapping regions of Chr 2 with each other, and which therefore are expected to be dissimilar. In comparison to other published comparisons between Hi-C experiments and simulations at the TAD scale [15,21,23,26,27,40,54], by eye, we judge the agreement displayed in Figs. 1*A* and *B* to be comparable or superior. To quantitatively compare the simulated and experimental contact maps, we examined the ratio of each pair of compared Hi-C maps in logarithmic scale, plotted as Figs. 1*E* and *F*, which further illustrate a good agreement between experiment and simulation of the same region and a mismatch between non-overlapping regions. Each pixel in the ratio maps shown in Figs. 1*E* and *F* is the ratio between the corresponding pixels, *E*_*n*_ and *S*_*n*_, whichever is larger, from the two compared Hi-C maps, respectively, *i*.*e*. 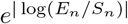. Therefore, all pixels in the ratio maps are always greater or equal to 1. Overall, Figure 1*E* is lightly shaded, indicating relatively small discrepancies overall between the simulation and experiment of Figs. 1*A* and *B*. By contrast, Figure 1*F* contains many more darker pixels, indicating relatively large differences between the experimental Hi-C maps of different genomic regions. Overall, we have simulated Hi-C maps for a total of five 1.2-Mb-sized genomic regions (only the middle 1 Mb regions are displayed to avoid any possible distortions from the simulation boundaries), one from each chromosome arm that exceeds 1.2 Mb in length. (The left arm of chromosome 3 was omitted, since it is 1.1 Mb long.) These five regions together span more than one-third of the entire fission yeast genome. In every case, we achieve good agreement between the experimental and simulated Hi-C maps (Fig. 1 and Supplementary Figs. 1–4).

**Figure 1:**
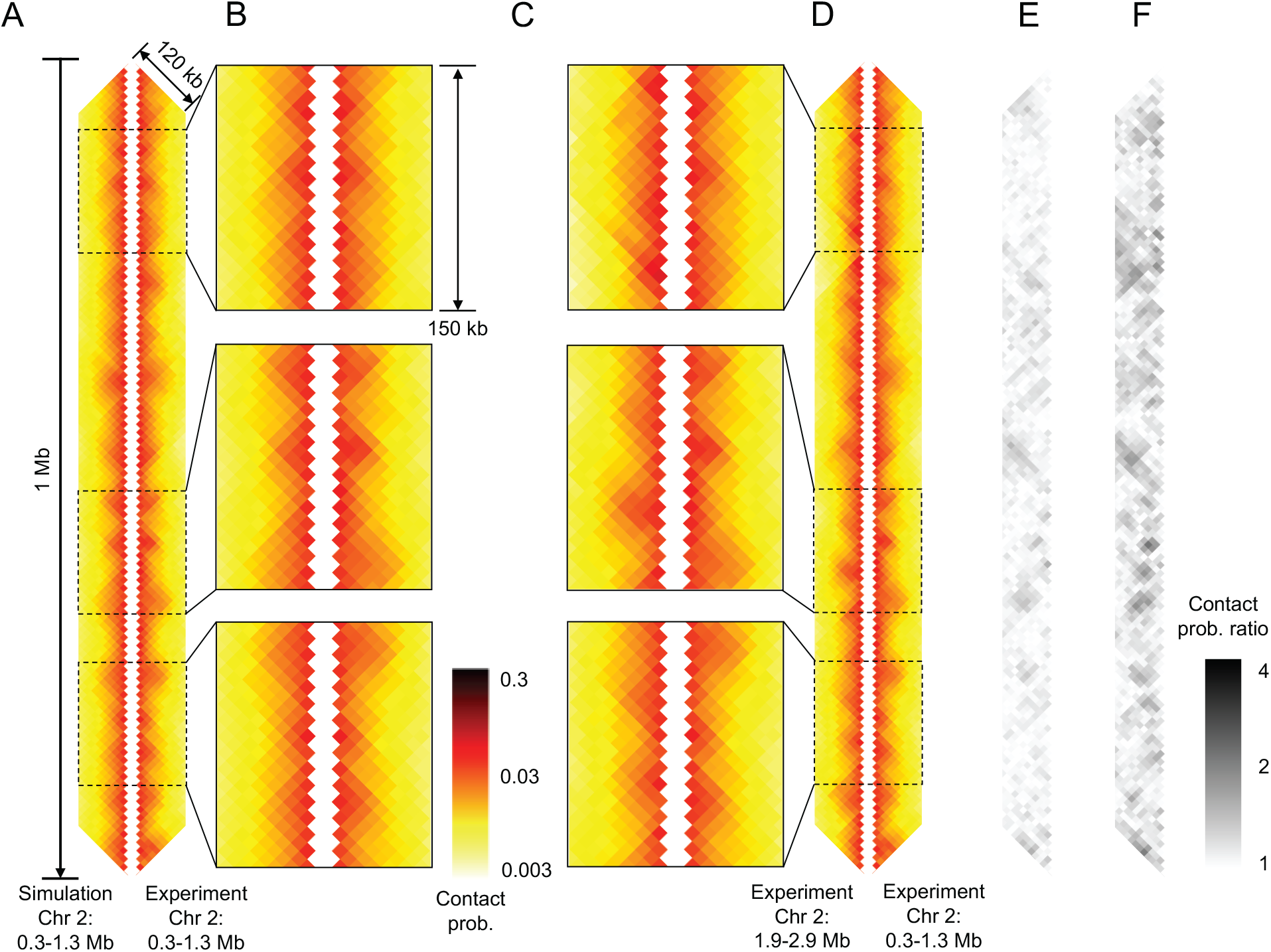
Conserved-current loop extrusion (CCLE) model recapitulates TAD-scale chromatin organization in interphase *S. pombe* using cohesin ChIP-seq data. (A) Comparison between Hi-C map of 1 Mb region, generated by the CCLE model (using cohesin ChIP-seq data from Ref. [6]), and the experimental Hi-C map [6] of the same region. (B), (C) Magnified comparisons of experimental and simulated Hi-C from panel (A), and of two experimental Hi-C from panel (D), respectively, for three representative sub-regions, each 150 kb in size, located at 0.39–0.54 Mb, 0.83–0.98 Mb, and 1.04–1.19 Mb, from top to bottom. (D) Comparison between two experimental Hi-C maps [6] from two different regions. All Hi-C maps show interactions up to a genomic separation of 120 kb. (E) Contact probability ratio maps between the simulated and experimental Hi-C shown in panel (A). Each pixel represents the ratio of contact probabilities of the corresponding pixels of the simulated and experimental Hi-C shown in panel (A). (F) Contact probability ratio maps between the two experimental Hi-C maps shown in panel (D). Pixels with darker shades indicate a poorer agreement than pixels with lighter shades. All maps are displayed in log-scale. The simulated Hi-C map is generated by the CCLE model using the best-fit parameters given in Table. 2 The optimization process is discussed in Supplementary Methods.

Supplementary Figures 5–9 also present the comparisons between the CCLE-predicted Hi-C maps and the newer Micro-C data for *S. pombe* from Hsieh *et al*. [36], both binned at a higher resolution of 2 kb. Since there are a reduced number of counts in each genomic pixel at 2 kb-resolution, the experimental contact map tends to be relatively noisy, which explains the relatively high MPR and low PCC values (given in the figure captions) compared to the Hi-C comparisons displayed at 10 kb-resolution. Nevertheless, the comparisons at 2 kb-resolution reveal clear left-right symmetry as well, indicating that CCLE also accurately describes the experimental contact maps at this higher resolution. Also included in Supplementary Figures 5–9 are comparisons between the original Hi-C maps of Mizuguchi *et al*. and the newer maps of Hsieh *et al*., binned to 10 kb-resolution. Inspection of these comparisons does not reveal any major new features in the interphase fission yeast Hi-C map on the 10-100 kb scale that were not already apparent from the Hi-C maps of Mizuguchi *et al*.

An additional, commonly-applied way to compare Hi-C experiments and simulations is to examine the mean contact probability, *P*(*s*), for loci with genomic separation, *s*. Figure 2 displays five *P*(*s*)-versus-*s* curves, each corresponding to one of the five regions simulated. Experimental and simulated mean contact probabilities are shown as the open circles and solid lines, respectively. Evidently, the experimental mean contact probabilities are very similar for different genomic regions, except for large separation contacts in the 1.2–2.4 Mb region of Chr 3, whose probability exceeds that of the other regions by about 15%. For every region, the simulated mean contact probability agrees well with its experimental counterpart, throughout the range of genomic separations studied (20 kb-500 kb), bolstering *a posteriori* the simple, analytic approach to polymer self-contacts, described in Methods.

**Figure 2:**
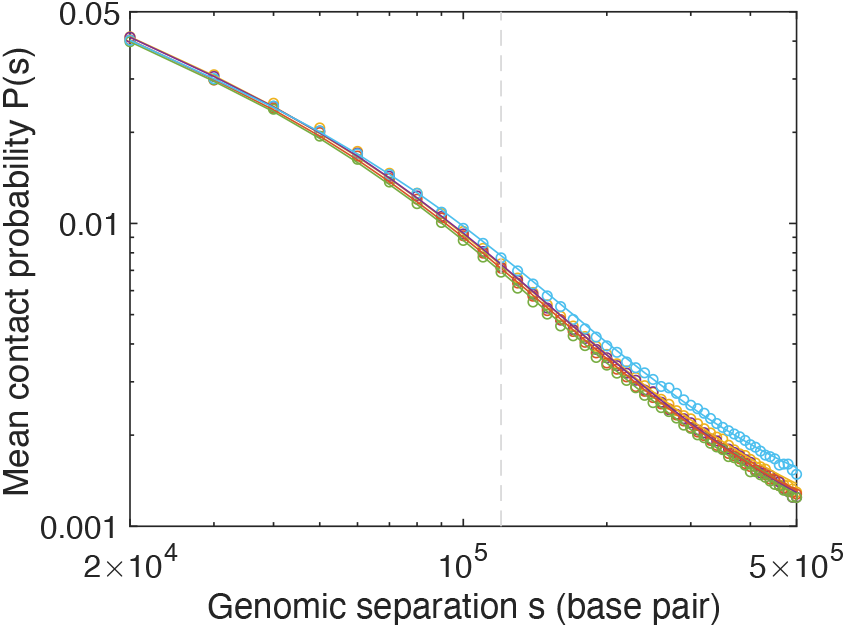
CCLE reproduces *P*(*s*) curves of experimental Hi-C contact maps across the entire *S. pombe* genome. Chromatin mean contact probabilities, *P*(*s*), as a function of genomic separation, *s*, are plotted in circles for five 1.2 Mb genomic regions, using experimental Hi-C contact map data from Ref. [6]. The corresponding *P*(*s*) curves of CCLE-simulated Hi-C contact maps are plotted as lines. Different colors represent different regions: yellow, 0.5-1.7 Mb of Chr 1; purple, 4.2-5.4 Mb of Chr 1; red, 0.2-1.4 Mb of Chr 2; green, 1.8-3.0 Mb of Chr 2; cyan, 1.2-2.4 Mb of Chr 3. The vertical gray dashed line indicates the maximum genomic separation displayed in the Hi-C comparison maps and ratio maps shown in Fig. 1.

To achieve the good agreement between simulation and experiment, evident in Figs. 1*A*, 1*B* and 2, we chose to quantify and minimize discrepancies between experimental and simulated Hi-C maps, similarly to Refs. [27, 54], using an objective function, that can be interpreted as the *mean pairwise ratio* (MPR) of experimental and simulated pixels, whichever is larger, averaged over all pixels:

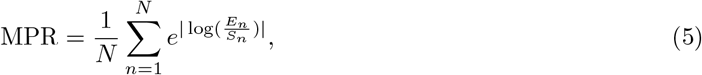

where *E*_*n*_ and *S*_*n*_ are the experimental and simulated contact probabilities, respectively, for pixel *n*, and *N* is the total number of pixels in the map of Fig. 1*A*. Note that the MPR value of the two compared maps is exactly the mean of all pixel values of their ratio map. A value of MPR closer to unity indicates better agreement. The MPR between the simulated and experimental Hi-C maps of Fig. 1*A* is 1.0911 (Table 1), while for the experimental Hi-C maps from two different regions (Fig. 1*D*), it is 1.1965 (Supplementary Table 1), showing a deviation from unity more than twice as large.

**Table 1:** Mean pairwise ratio (MPR) score and *P*(*s*)-scaled Pearson correlation coefficient (PCC) between experimental and model-generated Hi-C after model parameter optimization.

| Regions | MPR | PCC |
| --- | --- | --- |
| *S. pombe* Chr1: 0.6–1.6 Mb | 1.0984 | 0.5955 |
| *S. pombe* Chr1: 4.3–5.3 Mb | 1.1102 | 0.5249 |
| *S. pombe* Chr2: 0.3–1.3 Mb | 1.0911 | 0.6728 |
| *S. pombe* Chr2: 1.9–2.9 Mb | 1.0877 | 0.6493 |
| *S. pombe* Chr3: 1.3–2.3 Mb | 1.0902 | 0.6188 |
| *S. cerevisiae* Chr13: 290–790 kb (meiotic) | 1.2516 | 0.7409 |
| *S. cerevisiae* Chr10: 250–350 kb (mitotic) | 1.9536 | 0.4488 |
| *S. pombe* Chr2: 0.3–1.3 Mb (condensin) | 1.1793 | -0.0837 |

**Table 2:** Optimized parameters of CCLE simulations for different genomic regions in interphase *S. pombe*, meiotic *S. cerevisiae*, and mitotic *S. cerevisiae*.

| *S. pombe* | LEF density, $\rho$<br>(1/kb) | Mean processivity, $L$<br>(kb) | Persistence<br>length (nm) | Cohesive cohesin<br>density, $\rho_c$ (1/kb) | |
| --- | --- | --- | --- | --- | --- |
| Chr1:<br>0.5–1.7 Mb | 0.033 (± 0.001) | 30.0 (± 0.7) | 85 (± 19) | 0.037 (± 0.003) | |
| Chr1:<br>4.2–5.4 Mb | 0.033 (± 0.001) | 31.2 (± 0.8) | 75 (± 23) | 0.014 (± 0.003) | |
| Chr2:<br>0.2–1.4 Mb | 0.033 (± 0.001) | 30.0 (± 0.8) | 80 (± 21) | 0.050 (± 0.004) | |
| Chr2:<br>1.8–3.0 Mb | 0.033 (± 0.001) | 30.0 (± 0.6) | 80 (± 13) | 0.028 (± 0.005) | |
| Chr3:<br>1.2–2.4 Mb | 0.030 (± 0.001) | 33.3 (± 1.0) | 95 (± 17) | 0.043 (± 0.004) | |

| *S. cerevisiae* | LEF density,<br>$\rho$ (1/kb) | Mean processivity,<br>$L$ (kb) | Persistence<br>length (nm) | Cohesive cohesin<br>density, $\rho_c$ (1/kb) | Gaussian<br>$\sigma$ (kb) |
| --- | --- | --- | --- | --- | --- |
| Chr13:<br>240–840 kb<br>(meiotic) | 0.058 (± 0.003) | 38.4 (± 3.3) | 160 (± 19) | 0.018 (± 0.009) | 90 (± 25) |
| Chr10:<br>100–700 kb<br>(mitotic) | 0.033 (± 0.003) | 7.2 (± 0.2) | 60 (± 2) | 0 | 98 (± 3) |

LEF density is given in terms of number of LEFs per kilobase-pair; processivity is defined as the averaged LEF processivity in the corresponding region, in the absence of obstacles; cohesive cohesin density is also given in terms of number of cohesive cohesins per kilobase-pair. Parameters are optimized to minimize the mean pairwise ratio (MPR). See Supplementary Methods for details of statistical error calculation and the optimization process.

In addition to the MPR, we also calculated the *P*(*s*)-scaled Pearson correlation coefficient (PCC) for pairs of Hi-C maps (Tables 1 and Supplementary Table 1). The PCC for the simulated and experimental Hi-C maps of Fig. 1*A* is 0.6728, indicating a high correlation between the two maps. In contrast, the PCC for the experimental Hi-C maps shown in Fig. 1*D*, corresponding to non-overlapping genomic regions, is -0.0443 (Supplementary Table 1), indicating no correlation, as expected for the comparison between two different regions. For all five regions studied, the MPR scores and PCCs correspond to distinctly smaller differences overall and significantly greater correlation, respectively, between experimental and simulated Hi-C maps (Table 1) than between different experimental Hi-C maps (Supplementary Table 1), demonstrating that the CCLE model is able to predict and reproduce TAD patterns in *S. pombe*, based solely on cohesin ChIP-seq data.

To obtain the best agreement between simulation and experiment, we optimized the four model parameters, namely, LEF density (*ρ*), LEF processivity (*L*), cohesive cohesin density (*ρ*_*c*_), and chromatin persistence length, by minimizing Eq. 5, as described in more detail in Supplementary Methods. For each simulated region, the model parameters were optimized independently to their best-fit values, which are presented in Table 2. The optimized values of the LEF density and the LEF processivity are similar and have small errors, suggesting that the best-fit values of these parameters are robust and that cohesin-driven loop extrusion processes are essentially uniform across the *S. pombe* genome, at least at a resolution of 10 kb. The best-fit LEF density of 0.033 kb^−1^ corresponds to a cellular copy number of about 400 loop-extruding cohesin complexes per cell. If we also take the best-fit “cohesive cohesin density” parameter at face value, then there are approximately an additional 400 cohesive cohesins per cell, that is, 800 cohesins in total, which may be compared to the copy numbers from Ref. [55] of the constituents of the cohesion core complex, namely Psc3, Psm1, Psm3, and Rad21, of 723, 664, 1280, 173, respectively (mean 710). The best-fit values of the chromatin persistence length are also similar for different genomic regions, reflecting the similar behavior of the mean contact probability, *P*(*s*), versus genomic separation, *s*, for different genomic regions (Fig. 2). The best-fit values of the chromatin persistence length, which lie in the 75-95 nm range, may be compared to the value of 70 nm, reported in Ref. [56]. The relatively larger errors in the best-fit values of persistence length reflect the fact that our polymer model is only sensitive to the persistence length for large genomic separations (Eq. S36 and Supplementary Fig. 10). The bestfit values of the cohesive cohesin density show a greater variation for different genomic regions than the other parameters, hinting that the level of sister chromatid cohesion may vary across the genome. Alternatively, however, the apparent variation in this parameter could reflect different Psc3 ChIP-seq background levels or shifts for different regions (Methods).

In addition to cohesin, we also tested whether condensin, which is also a member of the SMC complex family shown to extrude DNA loops *in vitro* [57], could be another LEF that determines chromatin spatial configurations in *S. pombe*. However, when we carry out CCLE simulations using the ChIP-seq signal of interphase *S. pombe* condensin, the resultant best-fit simulated Hi-C map shows poor agreement with experiment (Supplementary Fig. 11 and Table 1), reinforcing that cohesin predominantly determines TAD-scale chromatin organization in interphase *S. pombe*.

#### CCLE self-consistently reproduces cohesin ChIP-seq data

In addition to predicting chromatin organization as measured by Hi-C maps and mean contact probability curves, loop extrusion simulations simultaneously yield position-dependent LEF occupancy probabilities. For a typical 200 kb-sized region of *S. pombe*’s Chr 2, Figure 3*A* compares the simulated time- and population-averaged probability that a chromatin lattice site is occupied by a LEF (red curve) to the corresponding experimental ChIP-seq data for Psc3 (blue curve), converted to occupancy probability, as described in Methods. Evidently, the simulated LEF occupancy probability matches the experimental Psc3 occupancy probability well. Indeed, as shown in Fig. 3*B*, the cross-correlation of experimental and simulated occupancy probability is nearly 0.7. In both cases, a number of peaks are apparent, extending above background to about twice, or less than twice, background. In the context of CCLE, this relatively weak contrast in occupancy probability gives rise to a corresponding relative lack of contrast in *S. pombe*’s interphase Hi-C pattern.

**Figure 3:**
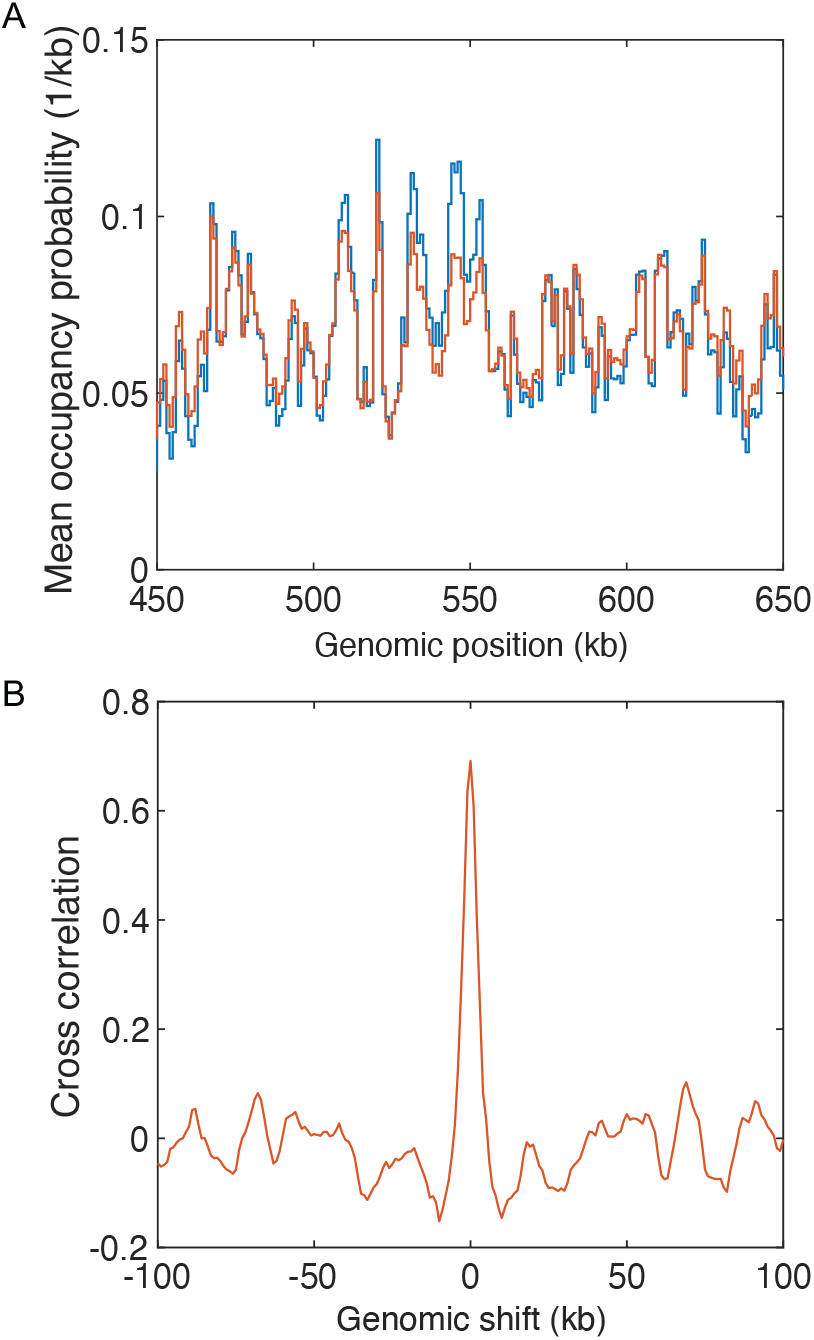
CCLE reproduces experimental cohesin occupancy landscape in *S. pombe*. (A) Comparison between the experimental cohesin occupancy probability landscape (blue) and the simulated LEF occupancy probability landscape by CCLE (red) in the 450–650 kb region of Chr 2 of interphase *S. pombe*. The occupancy probability curves are normalized by the corresponding optimized LEF density of 0.033 kb^−1^. (B) Cross-correlation between the experimental cohesin occupancy probability landscape and the simulated LEF occupancy probability landscape by CCLE, as a function of relative genomic shift.

While unsurprising, given that we set the position-dependent loop extrusion rates using the Psc3 ChIP-seq data, the good agreement between experimental and simulated occupancy probabilities implies that our simulations are self-consistent, and that the assumptions, leading to Eqs. 3 and 4, are valid for the parameters that are determined to best-describe the Hi-C contact map. The extent to which our best-fit simulations satisfy the assumptions, underlying CCLE, can be further assessed by examining the simulated distributions of the following dimensionless quantities, all of which should be small when CCLE is applicable:

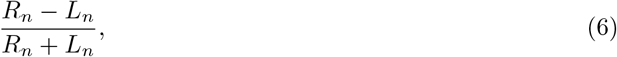

which specifies the fractional imbalance in the numbers of left- and right-moving LEF anchors at each lattice site;

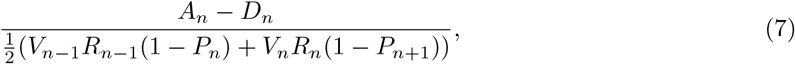

which is the ratio of the net current of right moving LEF anchors at each site, that violates current conservation, as a result of binding or unbinding, to the mean current of right-moving anchors, that satisfies current conservation; and

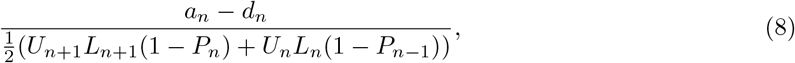

which is the corresponding conserved-current violation ratio for left-moving LEF anchors. As shown in Supplementary Fig. 12, all three of these quantities are distributed around zero with only small excursions (standard deviations ∼ 0.1), consistent with the assumptions, leading to Eqs. 3 and 4. While loop extrusion in interphase *S. pombe* seems to well satisfy the assumptions underlying CCLE, this may not always be the case.

#### Loop configurations in interphase *S. pombe*

The good agreement between our simulated Hi-C maps and experimental Hi-C maps suggests that the corresponding simulated loop configurations are realistic of loop configurations in live *S. pombe*. Figure 4*A* shows three representative simulated loop configurations for a 1.2 Mb region of Chr 2, corresponding to the best-fit parameters. In this figure, as in Ref. [58], each loop is represented as a semicircle connecting the genomic locations of the two LEF anchor points. Because the model does not permit LEF anchors to pass each other, correspondingly semicircles never cross, although they frequently contact each other and nest, as is apparent for the configurations in Fig. 4*A*. The distributions of loop sizes for all five regions simulated in *S. pombe* are presented in Fig. 4*B*. Evidently, the loop size distributions are similar for all five regions with an overall mean and standard deviation of 22.1 kb and 19.5 kb, respectively. The mean loop size may be compared to the number of base pairs within the chromatin persistence length, estimated to comprise 3.5 kb by taking the chromatin linear density to be 50 bp/nm [56]. Thus, typical loops contain several (∼ 6) persistence lengths of the chromatin polymer. Figure 4*C* shows the distributions of chromatin backbone segment lengths, *i*.*e*. the distributions of lengths of chromatin segments, that lie outside of loops. Again, these distributions are similar for all regions simulated with an overall mean and standard deviation of 28.9 kb and 26.8 kb, respectively, again corresponding to several (∼ 8) persistence lengths between loops. Since a LEF’s anchors bring the chromosomal loci, bound by the LEF anchors, into spatial proximity, loops lead to a significant linear compaction of the chromatin polymer. Figure 4*D* shows the distributions of chromatin compaction across an ensemble of loop configurations, defined as the fraction of the chromatin contour length contained within the backbone. These distributions too are similar for all five regions simulated, with overall mean and standard deviation of 0.4161 and 0.0663, implying that the chromatin polymer’s contour length in fission yeast is effectively 2.5-times shorter with loops than without.

**Figure 4:**
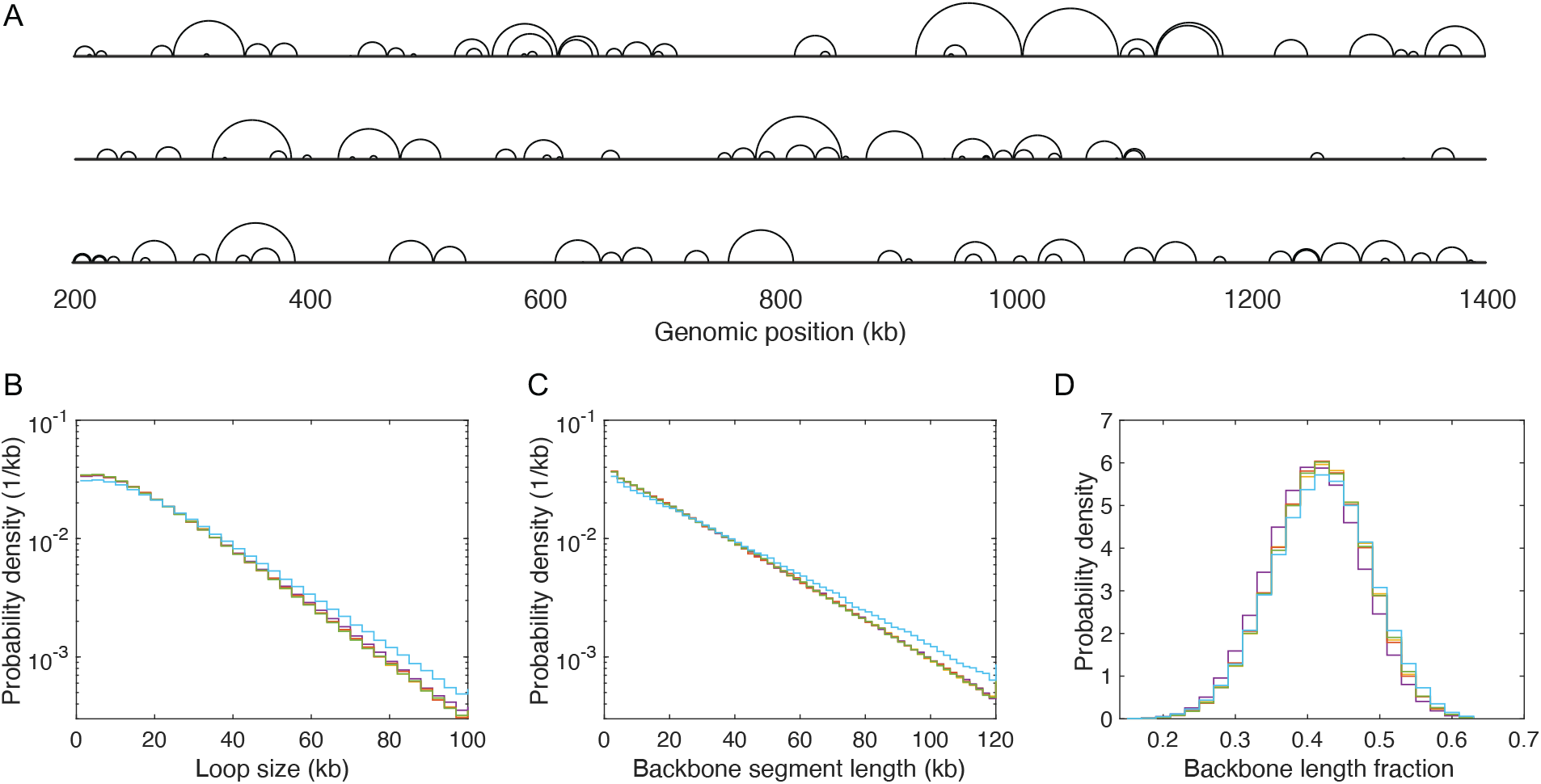
Loop configurations and properties. (A) Snapshots of three representative simulated loop configurations of interphase *S. pombe*’s 200–1400 kb region of Chr 2. In each case, the chromatin backbone is represented as a straight line, while loops are represented as semicircles connecting loop anchors, following Ref. [58]. Because LEF anchors block each other, loops can nest but they cannot cross. (B), (C), and (D) Distributions of loop size, backbone segment length, and chromatin compaction ratio (as measured by the fraction of the chromatin contour length within the backbone), respectively, for the five genomic regions of interphase *S. pombe*, listed in Table 2: Yellow, Chr 1: 0.5–1.7 Mb; purple, Chr 1: 4.2–5.4 Mb; red, Chr 2: 0.2–1.4 Mb; green, Chr 2: 1.8–3.0 Mb; cyan, Chr 3: 1.2–2.4 Mb.

#### Diffusion capture model does not reproduce experimental interphase *S. pombe* Hi-C maps

Because Hi-C and ChIP-seq both characterize chromatin configuration at a single instant of time, and do not provide any direct time-scale information, an alternative mechanism that somehow generates the same instantaneous loop distributions and loop correlations as loop extrusion would lead to the same Hi-C map as does loop extrusion. This observation has motivated consideration of diffusion capture models in which loops occur and persist as a result of loop capture via binding events [11,14,15,17,20,22–24,59,60]. Such a scenario is well recognized for a number of isolated enhancer-gene loci pairs in, for example, *Drosophila melanogaster* [61]. A major obstacle to the application of such models across the genome is that there is no physical basis for diffusion capture models to give rise to the approximately-exponential loop size distributions, which emerge naturally from the loop extrusion model and well describe Hi-C maps (Fig. 4). Instead, the loops in diffusion capture models can be expected to realize an equilibrium, power-law distribution of loop sizes (Supplementary Fig. 15*D* ), corresponding to the return probability of a random walk. To investigate quantitatively how well a physically-sensible diffusion capture model can describe Hi-C maps, while remaining consistent with cohesin ChIP-seq, we considered a diffusion capture model in which the probability that a loop connects sites *i* and *j* is proportional to

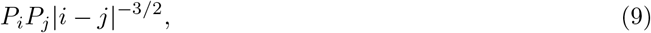

where *P*_*i*_ is the probability that site *i* is occupied by cohesin, which we determine from cohesin ChIP-seq data without distinguishing between cohesive and loop-extruding cohesins, and the factor 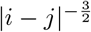 corresponds to the probability that a unconstrained random walk returns to its starting point after *i* − *j* steps. (Self-avoidance would change the exponent, but not the power-law behavior.)

By using the experimental interphase *S. pombe* cohesin (Psc3) ChIP-seq data to determine *P*_*i*_, we have carried out Monte Carlo simulations of this model with only two possible parameters: LEF density and chromatin persistence length, both derived from the CCLE-best-fit values for *S. pombe*. These simulations yield an ensemble of loop configurations, in turn allowing us to calculate the corresponding Hi-C maps in the same way that we calculate Hi-C maps from the ensemble of loop configurations generated by CCLE simulations. Supplementary Figure 15*A* compares a portion of the Hi-C map, simulated on the basis of the diffusion capture model, to the corresponding experimental Hi-C. Also illustrated in Supplementary Figure 15*B* is the ratio map between the diffusion-capture-simulated Hi-C and experiment, showing more darker pixels than the ratio map between the CCLE-simulated Hi-C and the experiment (Fig. 1*E*). Evidently, the diffusion capture model gives rise to a Hi-C map that provides a much poorer description of the experimental Hi-C map (with an MPR of 1.1754 and a PCC of 0.2675), than does CCLE, largely failing to reproduce the inhomogeneous pattern of squares corresponding to TADs and to match the measured *P*(*s*) (Supplementary Fig. 15*C* ). This comparison suggests that loop extrusion-based models for chromatin organization should be much preferred over diffusion capture models.

### CCLE Describes TADs and Loop Configurations in Meiotic *S. cerevisiae*

#### CCLE simulations accurately reproduce experimental Hi-C maps of meiotic *S. cerevisiae*

To further examine the ability of CCLE to describe TAD-scale chromatin organization, we next sought to describe the Hi-C map of meiotic *S. cerevisiae* from Ref. [54], using the ChIP-seq data of the meiotic cohesin core subunit, Rec8, from Ref. [62]. In contrast to the semi-dilute polymer solution envisioned to describe chromatin in interphase, in meiosis, the chromatin polymer is significantly compacted and is conspicuously organized about the chromosomal axis. Therefore, meiotic chromatin represents a very different polymer state than interphase chromatin, in which to test CCLE. To empirically account for increased polymer volume exclusion as a result of this more compacted polymer state, we scale the *P*(*s*) of the simulated Hi-C by a Gaussian scaling factor with standard deviation, *σ*, given in Table 2.

The right-hand side of Fig. 5*A* depicts a 500 kb portion of the experimental Hi-C map of meiotic *S. cerevisiae*’s Chr 13 from 290 kb to 790 kb, using a logarithmic false-color intensity scale. In comparison, the left-hand side of Fig. 5*A* presents the corresponding best-fit simulated Hi-C map. Both maps are shown at 2 kb resolution (the original simulated Hi-C map was at 500 bp resolution and binned to 2 kb to match the experimental resolution). Figure 5*B* magnifies the experiment-simulation comparison for three representative 80 kb sub-regions. Both Figures 5*A* and *B* reveal similar patterns of high-probability contacts for the experimental and simulated Hi-C maps, manifested in a high-degree of left-right symmetry. However, in contrast to the patch-like TAD patterns featured in the experimental and simulated Hi-C maps of interphase *S. pombe*, which consist of overlapping squares with more-or-less evenly-distributed enhanced contact probability, the TADs of meiotic *S. cerevisiae* are dominated by discrete lines of high contact probability and their intersection points. Nevertheless, Figures 5*A* and *B* make it clear that in spite of their strikingly different appearances to the TAD patterns of interphase *S. pombe*, the grid-like patterns of TADs in meiotic *S. cerevisiae* are also well reproduced by CCLE model. In addition, the three quantities, given by Eqs. 6, 7, and 8, are distributed around zero with relatively small fluctuations (Supplementary Fig. 13), indicating that CCLE model is self-consistent in this case also.

**Figure 5:**
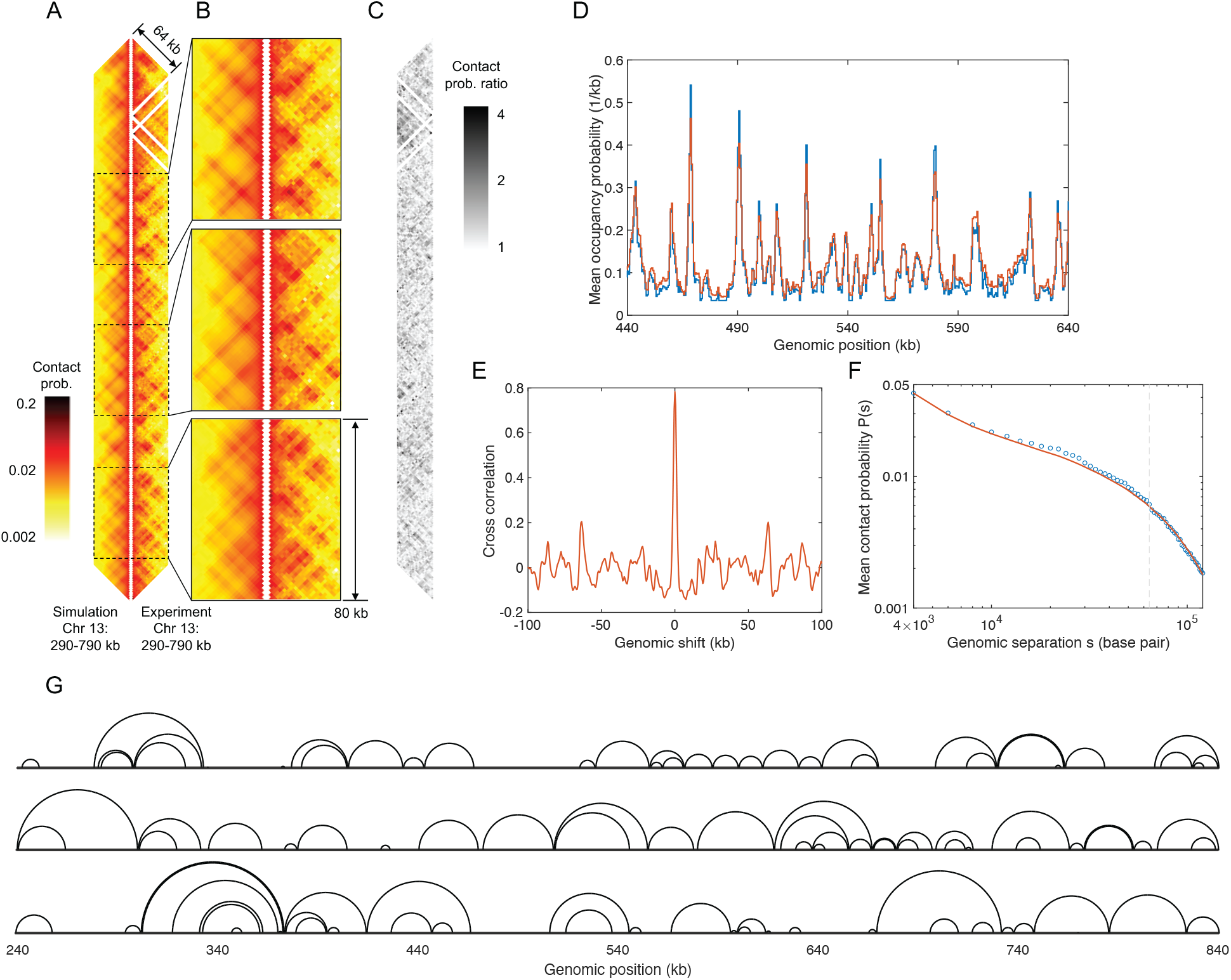
Conserved-current loop extrusion (CCLE) model recapitulates TAD-scale chromatin organization in meiotic *S. cerevisiae* using cohesin ChIP-seq data. (A) Comparison between the simulated Hi-C map of 290–790 kb region of Chr 13, generated by the CCLE model (using meiotic Rec8 ChIP-seq data from Ref. [62]), and the experimental Hi-C map [54] of the same region. Both Hi-C maps show interactions up to a genomic separation of 64 kb. (B) Magnified experiment-simulation Hi-C comparisons for three representative sub-regions of 80 kb in size: 412–492 kb, 546–626 kb, and 672–752 kb, from top to bottom. (C) Contact probability ratio map between the experimental and simulated Hi-C maps in panel (A). (D) Normalized experimental meiotic cohesin (Rec8) occupancy probability (blue) and simulated LEF occupancy probability landscape (red). The occupancy probability curves are plotted for 440–640 kb of Chr 13 and are normalized by the corresponding optimized LEF density of 0.058 kb^−1^. (E) Cross-correlation between the experimental meiotic cohesin (Rec8) occupancy landscape and the simulated LEF occupancy probability landscape, as a function of relative genomic shift. (F) Chromatin mean contact probability, *P*(*s*), plotted as a function of genomic separation, *s*, for the experimental (blue circles) and simulated (red line) Hi-C, scaled by the Gaussian scaling factor as described in Methods. The vertical gray dashed line indicates the maximum genomic separation displayed in the Hi-C comparison map in panels (A) and (B). (G) Snapshots of three representative simulated meiotic loop configurations in the 240–840 kb region of Chr 13. In each case, the chromatin backbone is represented as a straight line, while loops are represented as semicircles connecting loop anchors, following Ref. [58]. Because LEF anchors block each other, loops can nest but they cannot cross, although they frequently come into contact. Simulation results shown in this figure are generated by CCLE using the best-fit parameters given in Table 2. The optimization process is discussed in Supplementary Methods.

The very different appearances of the Hi-C maps of interphase *S. pombe* and meiotic *S. cerevisiae* directly follow from the much greater contrast of meiotic *S. cerevisiae*’s cohesin ChIP-seq profile, which consists of a dense pattern of strong, narrow peaks, which extend above the background to reach 4 or 5 times background (Fig. 5*D*). Also different is the best-fit value of the LEF density, which is about twice as large in meiotic *S. cerevisiae* as in interphase *S. pombe* (Table 2). However, the best-fit value for the LEF processivity of meiotic *S. cerevisiae* is similar to that of interphase *S. pombe*, suggesting loop-extruding cohesins possess similar properties in both species. Values for the LEF density (0.058 kb^−1^) and processivity (38.4 kb) may be compared to the best-fit values given in Ref. [54], of 0.03-04 kb^−1^ and 64-76 kb, respectively. The best-fit value of the cohesive cohesin density is noticeably smaller for meiotic *S. cerevisiae* than for interphase *S. pombe*. It is also significantly smaller than the best-fit density of loop-extruding cohesins, suggesting that the preponderance of cellular cohesins are involved in loop extrusion in meiotic *S. cerevisiae*. The best-fit value of persistence length of meiotic chromatin in *S. cerevisiae* is about twice that of interphase chromatin in *S. pombe*, potentially indicating stiffer chromatin due to a more compact chromatin state.

In spite of the agreement between simulation and experiment, evident in Fig. 5*A* and *B*, the experiment-simulation comparison for meiotic *S. cerevisiae* shows a higher MPR and lower PCC than for interphase *S. pombe* (Table 1), indicating a poorer fit. However, we ascribe this poorer fit, at least in part, to the larger experimental errors of the former data set. These larger experimental errors are apparent when we calculate the MPR and the PCC for the two duplicate Hi-C data sets available in Ref. [54], which take values of 1.5714 and 0.8180, respectively. The MPR score reveals greater discrepancies between the two nominally identical meiotic *S. cerevisiae* data sets than between the experiment and the best-fit simulation. Furthermore, the PCC score of the comparison between two nominally identical data sets is not significantly higher value than that for the experiment-simulation comparison.

#### CCLE self-consistently reproduces Rec8 ChIP-seq data in meiotic *S. cerevisiae*

For a typical 200-kb-sized region of Chr 13 of *S. cerevisiae*, Figure 5*D* compares the simulated time- and population-averaged probability that a chromatin lattice site is occupied by a LEF (red curve) to the corresponding experimental ChIP-seq data for Rec8 (blue curve), converted to occupancy probability, as described in Methods. Clearly, the simulated LEF occupancy probability matches the experimental Rec8 occupancy probability well, with a cross-correlation value of almost 0.8 (Fig. 5*E*).

#### Loop configurations in meiotic *S. cerevisiae*

Figure 5*G* shows three representative simulated loop configurations for the 240–840 kb region of Chr 13, corresponding to the best-fit parameters. In comparison to the loop configurations in interphase *S. pombe*, the loops in meiotic *S. cerevisiae* appear more regularly spaced, corresponding to the more regularly-distributed peaks of meiotic *S. cerevisiae*’s cohesin ChIP-seq data. Supplementary Figures 16*A* and *B* present the distributions of loop sizes and backbone segment lengths, respectively, for the same region in *S. cerevisiae*. The mean and standard deviation of these quantities are: 19.82 and 16.06 kb (mean), and 15.97 and 14.67 kb (SD), respectively. Supplementary Figure 16*C* shows the distributions of chromatin relative compaction, whose mean and standard deviation are 0.2129 and 0.0702, *i*.*e*. the chromatin polymer’s contour length in meiotic *S. cerevisiae* is effectively 5-times shorter with loops than without, twice as compact as the interphase *S. pombe*’s chromatin. Motivated by the experimental observation that the inactivation of “cohesin release factor”, Wpl1, gives rise to larger loops [41, 42, 63, 64], we performed CCLE simulations of wild-type and Wpl1-depleted meiotic *S. cerevisiae*. Indeed, we observe enhanced loop sizes in Wpl1-depleted cells with a mean loop size that is two and a half times larger than in wild-type cells (Supplementary Fig. 17).

### CCLE Describes TADs and Loop Configurations in Mitotic *S. cerevisiae*

Next, we applied CCLE to the Hi-C map of mitotic *S. cerevisiae* using the mitotic ChIP-seq data for the cohesin core protein, Mcd1, from Ref. [42]. The right hand side of Fig. 6*A* displays the Hi-C map of the 250–350 kb region of mitotic *S. cerevisiae*’s chromosome 10 at 500 bp resolution, reproducing the Hi-C data shown in Fig. 2*A* of Ref. [42], up to a genomic separation of 45 kb. All else being equal, as a result of the 500 bp-resolution, there are a reduced number of counts in each genomic pixel, compared to Hi-C maps displayed at lower resolutions. It follows that this experimental contact map is relatively noisy compared to both interphase *S. pombe* and meiotic *S. cerevisiae*. Nevertheless, standing above a field of relatively weak contacts, it is apparent that mitotic *S. cerevisiae*’s Hi-C map is primarily characterized by the presence of a number of prominent, isolated points of high-probability contacts – often called “puncta”. Unsurprisingly, mitotic *S. cerevisiae*’s Hi-C map is very different from that of interphase *S. pombe* (Fig. 1). However, it also appears distinct from the Hi-C map of meiotic *S. cerevisiae* (Fig. 5), in spite of the fact that the cohesin ChIP-seq of mitotic *S. cerevisiae* shows the same peak locations as the cohesin ChIP-seq of meiotic *S. cerevisiae*, as illustrated in Supplementary Figure 18*A*. Importantly, however, the cohesin ChIP-seq peaks are higher and narrower in the mitotic case than in the meiotic case, while the ChIP-seq signal between the peaks is suppressed in the mitotic case relative to the meiotic case. The lack of cohesin background signal in the mitotic case likely suggests that the mitotic cohesive cohesin density is very low.

**Figure 6:**
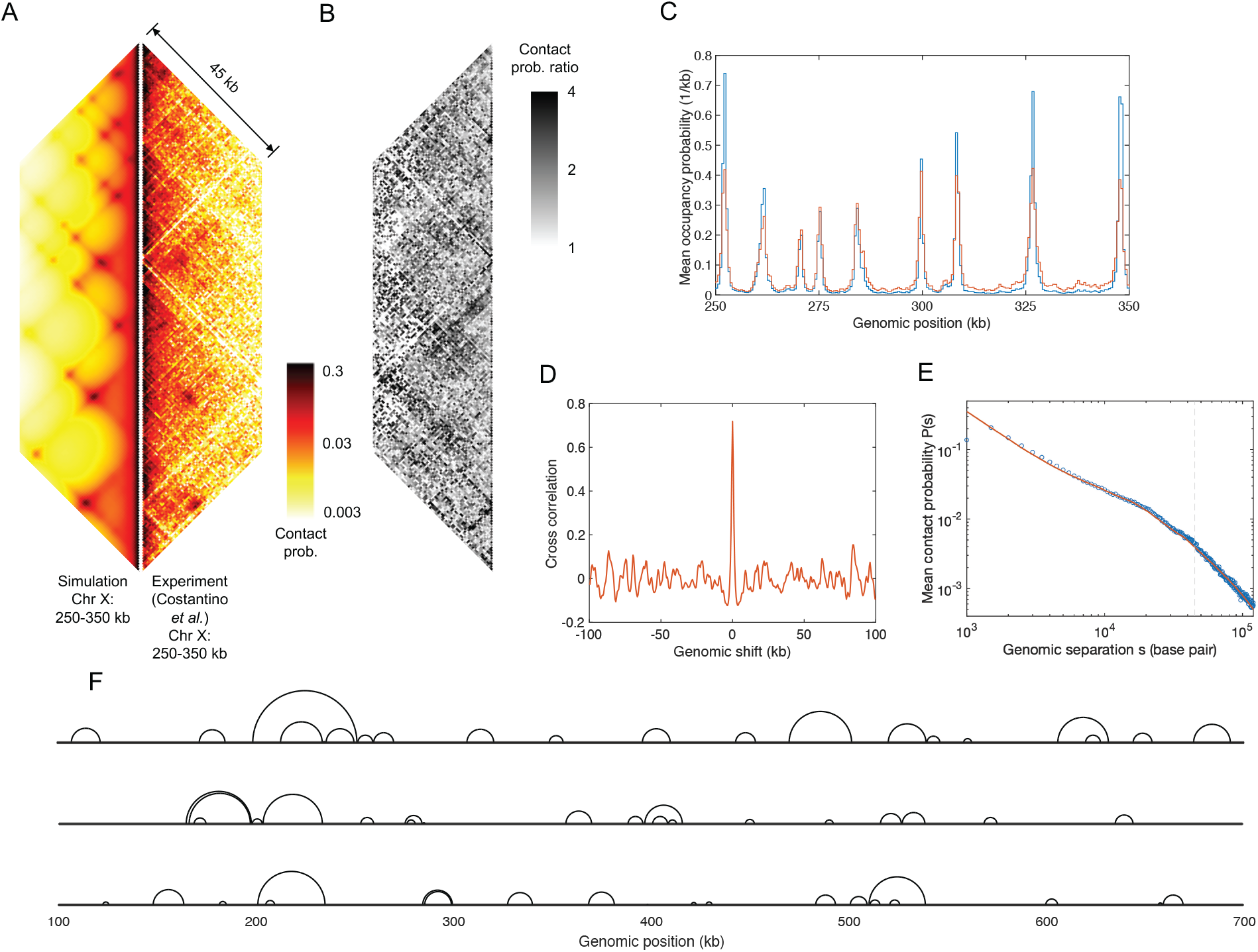
Conserved-current loop extrusion (CCLE) model recapitulates TAD-scale chromatin organization in mitotic *S. cerevisiae* using cohesin ChIP-seq data. (A) Comparison between the Hi-C map of 250–350 kb of Chr 10, generated by the CCLE model (using mitotic Mcd1 ChIP-seq data from Ref. [42]), and the experimental Hi-C map [42] of the same region. Both Hi-C maps show interactions up to a genomic separation of 45 kb. (B) Contact probability ratio map between the experimental and simulated Hi-C in panel (A). (C) Normalized experimental mitotic cohesin (Mcd1) occupancy probability (blue) and simulated LEF occupancy probability landscape (red). The occupancy probability curves are normalized by the corresponding optimized LEF density of 0.033 kb^−1^. (D) Cross-correlation between the experimental mitotic cohesin (Mcd1) occupancy landscape and the simulated LEF occupancy probability landscape, as a function of relative genomic shift. (E) Chromatin mean contact probability, *P*(*s*), plotted as a function of genomic separation, *s*, for the experimental (blue circles) and simulated (red line) Hi-C , scaled by the Gaussian correction factor as described in Methods. The vertical gray dashed line indicates the maximum genomic separation displayed in the Hi-C comparison map in panel (A). (F) Snapshots of three representative simulated mitotic loop configurations in the 100–700 kb region of Chr 10. In each case, the chromatin backbone is represented as a straight line, while loops are represented as semicircles connecting loop anchors, following Ref. [58]. Because LEF anchors block each other, loops can nest but they cannot cross, although they frequently come into contact. Simulation results shown in this figure are generated by CCLE using the best-fit parameters given in Table 2. The optimization process is discussed in Supplementary Methods.

In comparison with the experimental Hi-C map on the right-hand side Fig. 6*A*, the left-hand side of Fig. 6*A* shows the corresponding best-fit simulated Hi-C map generated by the CCLE model. The CCLE-simulated Hi-C map shows puncta, that well match the experimental puncta, demonstrating that CCLE is also able to well-describe chromatin looping in mitotic *S. cerevisiae*. In spite of the agreement between simulation and experiment, the experiment-simulation comparison for mitotic *S. cerevisiae* reveals a higher MPR and lower PCC compared to both interphase *S. pombe* and meiotic *S. cerevisiae*, which may indicate a poorer fit or noisier experimental data (Table 1). Given the noisier nature of the Hi-C data at this higher resolution, we ascribe the lower MPR and PCC scores to the latter case. However, we ascribe this poorer fit to the noisy nature of Hi-C data at higher resolutions. Figure 6*B* shows the contact probability ratio map between the experimental and simulated Hi-C maps from which the noise present in these data is further apparent.

Interestingly, except the Gaussian scaling factors, the best-fit values of all other model parameters for mitotic *S. cerevisiae* (Table 2) are quite different from those for meiotic *S. cerevisiae*. Specifically, the LEF density is about half of the value observed in meiotic *S. cerevisiae*, the LEF processivity for mitotic *S. cerevisiae* is only about 20% of the value observed in meiotic *S. cerevisiae*, and the persistence length is only about 40%. It is important to realize that, since the persistence length and the Gaussian scaling factor primarily determine the long-range genomic contact, altering one of these parameters is likely to affect the best-fit value of the other. To reflect the lack of background signal in the experimental mitotic cohesin ChIP-seq data, we set the value of cohesive cohesin density to zero (Table 2). The circumstances of fewer and less extended cohesin loops in the mitotic case, compared to the meiotic case, are reflected in the sparser and smaller loops in Fig. 6*F*, compared to Fig. 5*G*. Thus, according to the CCLE model, the different appearances of the Hi-C maps of meiotic and mitotic *S. cerevisiae* originate from the differences in their ChIP-seq profiles, as well as variations in cohesin density and processivity between the two cases.

Figure 6*C* displays the normalized experimental mitotic cohesin occupancy probability derived from Mcd1 ChIP-seq (blue line), versus LEF occupancy probability from the CCLE simulation (red line). Both landscapes appear very similar, with strong peaks at the same genomic locations (although the experimental peaks are higher), and they exhibit a very strong cross-correlation (Fig. 6*D*). As shown in Fig. 6*E*, the experimental chromatin mean contact probability, *P*(*s*), as a function of genomic separation, *s*, is also well reproduced by the CCLE model.

However, in comparison to the previous two examples, the three quantities, given by Eqs. 6, 7, and 8, have broader distributions and larger standard deviations around the mean values, which are nevertheless close to zero (Supplementary Fig. 14). Interestingly, lattices with a greater imbalance of left- and right-moving LEFs are found on both sides of cohesin peaks. Indeed, Supplementary Figure 14*C* presents the cross-correlation between the cohesin ChIP-seq and the imbalance landscape, where the trough-peak pattern around zero suggests that there are more right-moving LEF anchors approaching the left sides of cohesin peaks and more left-moving LEF anchors approaching the right sides of cohesin peaks. Furthermore, Supplementary Figures 14*F* and *I*, which present the cross-correlations between cohesion ChIP-seq and current violation landscapes, show significant troughs at zero, suggesting that LEF unbinding events outnumber the binding events at cohesin peaks. We ascribe these greater violations of CCLE’s assumptions at the locations of cohesin peaks in part to the low processivity of mitotic cohesin in *S. cerevisiae*, compared to that of meiotic *S. cerevisiae* and interphase *S. pombe*, and in part to the low CCLE loop extrusion rate at the cohesin peaks. While CCLE assumptions are violated at these sites, the model (1) does allow for these approximations to be modeled, (2) provides internal metrics to check for violations, and (3) still recovers chromatin contact distribution correctly (based on Hi-C comparison). In the future, we plan to develop an improved version of CCLE that will self-consistently account for binding and unbinding, as well as imbalance of left- and right-moving LEFs, as follows: (1) from the previous best-fit simulation results, evaluate the empirical binding/unbinding rates and left/right-moving LEF imbalance for each lattice position, (2) update and solve the exact master equations, Eq. 1 and 2, for the position-dependent loop extrusion rates using the empirical binding/unbinding rates and left/right-moving LEF proportions, (3) use the new loop extrusion rates to run CCLE simulations and obtain a new set of best-fit simulation results, (4) iterate until the simulation parameters converge.

## Discussion

By examining an approximate steady-state solution of master equations describing the motions of chromatin loop extruding complexes, we have been led to a new version of the loop extrusion factor model – the conserved-current loop extrusion (CCLE) model – that does not require the input of genomic positions of boundary elements and uses cohesin ChIP-seq data as the sole input. To demonstrate its utility, we applied the CCLE model to accurately reproduce the TAD-scale (∼ 10 − 100 kb) Hi-C maps for each of interphase *S. pombe* and mitotic *S. cerevisiae*, for the first time, and of meiotic *S. cerevisiae*, in effect recapitulating the results of Ref. [54] but using a model with an improved physical basis. Importantly, the fact that the CCLE model can convert cohesin ChIP-seq data into an accurate Hi-C map, highlights that essential aspects of the three-dimensional chromatin configuration are encoded in the one-dimensional cohesin distribution, and strongly suggests that the loop configurations generated, as well as the best-fit values of the density and processivity of loop-extruding cohesins, are realistic. Not limited to cohesin, other SMC complexes, such as condensin [57, 65–67] and Smc5/6 [68], could play similar roles in other organisms or different stages of the cell cycle. The CCLE model is agnostic to how SMC distributions are established, providing greater flexibility to account for different LEF-chromatin interactions, such as loop extrusion barriers blocking LEFs [27, 69, 70] and arrays of factors slowing down LEF translocation [69–71]. Overall, our results provide compelling evidence that chromatin organization at the TAD scale in interphase *S. pombe*, as well as in meiotic and mitotic *S. cerevisiae*, is primarily the result of loop extrusion, and that the cohesin complex is the dominant loop extrusion factor, marking the base of all, or the overwhelming majority, of TAD-scale loops in these systems.

In vertebrates, CTCF defines the locations of most TAD boundaries. It is interesting to ask what might play that role in interphase *S. pombe* as well as in meiotic and mitotic *S. cerevisiae*. A number of papers have suggested that convergent gene pairs are correlated with cohesin ChIP-seq in both *S. pombe* [72, 73] and *S. cerevisiae* [64, 73–77]. Because CCLE ties TADs to cohesin ChIP-seq, a strong correlation between cohesin ChIP-seq and convergent gene pairs would be an important clue to the mechanism of TAD formation in yeasts. To investigate this correlation, we introduce a convergent-gene variable that has a non-zero value between convergent genes and an integrated weight of unity for each convergent gene pair. Supplementary Figure 18*A* shows the convergent gene variable, so-defined, alongside the corresponding cohesin ChIP-seq for meiotic and mitotic *S. cerevisiae*. It is apparent from this figure that a peak in the ChIP-seq data is accompanied by a non-zero value of the convergent-gene variable in about 80% of cases, suggesting that chromatin looping in meiotic and mitotic *S. cerevisiae* may indeed be tied to convergent genes. Conversely, about 50% of convergent genes match peaks in cohesion ChIP-seq. The cross-correlation between the convergent-gene variable and the ChIP-seq of meiotic and mitotic *S. cerevisiae* is quantified in Supplementary Figures 18*B* and *C*. By contrast, in interphase *S. pombe*, cross-correlation between convergent genes and cohesin ChIP-seq in each of five considered regions is unobservably small (Supplementary Figure 19*A*), suggesting that convergent genes per se do not have a role in defining TAD boundaries in interphase *S. pombe*.

Although “bottom-up” models which incorporate explicit boundary elements do not exist for non-vertebrate eukaryotes, one may wonder how well such LEF models, if properly modified and applied, would perform in describing Hi-C maps with diverse features. To this end, we examined the performance of the model described in Ref. [54] in describing the Hi-C map of interphase *S. cerevisiae*. Reference [54] uses cohesin ChIP-seq peaks in meiotic *S. cerevisiae* to define the positions of loop extrusion barriers which either completely stall an encountering LEF anchor with a certain probability or let it pass. We apply this “explicit barrier” model to interphase *S. pombe*, using its cohesin ChIP-seq peaks to define the positions of loop extrusion barriers, and using Ref. [54]’s best-fit value of 0.05 for the pass-through probability. Although the applicability of a pass-through probability of 0.05, derived from meiotic *S. cerevisiae*, to interphase *S. pombe* is uncertain, in fact, simulations reveal that varying this quantity across the range from 0.005 to 0.5 causes only modest changes in the corresponding simulated Hi-C maps, as illustrated in Supplementary Figs. 20*E* and *F*. Supplementary Figure 20*A* presents the simulated Hi-C map of the 0.3–1.3 kb region of Chr 2 of interphase *S. pombe* in comparison with the corresponding Hi-C data. It is evident that the explicit barrier model provides a poorer description of the Hi-C data of interphase *S. pombe* compared to the CCLE model, as indicated by the MPR and Pearson correlation scores of 1.6489 and 0.2267, respectively. While the explicit barrier model appears capable of accurately reproducing Hi-C data with punctate patterns, typically accompanied by strong peaks in the corresponding cohesin ChIP-seq, it seems less effective in cases such as in interphase *S. pombe*, where the Hi-C data lacks punctate patterns and sharp TAD boundaries, and the corresponding cohesin ChIP-seq shows low-contrast peaks. The success of the CCLE model in describing the Hi-C data of both *S. pombe* and *S. cerevisiae*, which exhibit very different features, suggests that the current paradigm of localized, well-defined boundary elements may not be the only approach to understanding loop extrusion. By contrast, CCLE allows for a concept of continuous distribution of position-dependent loop extrusion rates, arising from the aggregate effect of multiple interactions between loop extrusion complexes and chromatin. This paradigm offers greater flexibility in recapitulating diverse features in Hi-C data than strictly localized loop extrusion barriers.

In our current CCLE implementation, cohesin binds to chromatin at random locations. However, the fission yeast protein, Mis4, has been previously identified, as a component of the cohesin loading complex [78–80], prompting us to envision a modification of the model to incorporate position-dependent cohesin binding with a binding rate proportional to the Mis4 ChIP-seq signal. However, for *S. pombe*, the effect of such a modification on the resultant loop configurations must necessarily be small, because cohesin distribution essentially defines the steady-state distribution of loop anchors but there is no correlation between the Mis4 and Psc3 ChIP-seq signals (Supplementary Fig. 21*A*). Therefore, the overall distribution of cohesin along the genome (given by the Psc3 ChIP-seq) is independent of where the cohesin was putatively loaded (specified by the Mis4 ChIP-seq). This observation suggests that, in *S. pombe* at least, the collective spreading of cohesins following association to chromatin is sufficiently large, so as to obscure their initial positions.

As noted above, the input for our CCLE simulations of chromatin organization in *S. pombe* was the ChIP-seq of Psc3, which is a component of the cohesin core complex [81]. Accordingly, Psc3 ChIP-seq represents how the cohesin core complex is distributed along the genome. In *S. pombe*, the other components of the cohesin core complex are Psm1, Psm3, and Rad21. Because these proteins are components of the cohesin core complex, we expect that the ChIP-seq of any of these proteins would closely match the ChIP-seq of Psc3, and would equally well serve as input for CCLE simulations of *S. pombe* genome organization. Supplementary Figure 21*C* confirms significant correlations between Psc3 and Rad21. In light of this observation, we then reason that the CCLE approach offers the opportunity to investigate whether other proteins beyond the cohesin core are constitutive components of the loop extrusion complex during the extrusion process (as opposed to cohesin loading or unloading). To elaborate, if the ChIP-seq of a non-cohesin-core protein is highly correlated with the ChIP-seq of a cohesin core protein, we can infer that the protein in question is associated with the cohesin core and therefore is a likely participant in loop-extruding cohesin, alongside the cohesin core. Conversely, if the ChIP-seq of a putative component of the loop-extruding cohesin complex is uncorrelated with the ChIP-seq of a cohesin core protein, then we can infer that the protein in question is unlikely to be a component of loop-extruding cohesin, or at most is transiently associated with it.

For example, in *S. pombe*, the ChIP-seq of the cohesin regulatory protein, Pds5 [80], is correlated with the ChIP-seq of Psc3 (Supplementary Fig. 21*B* ) and with that of Rad21 (Supplementary Fig. 21*D* ), suggesting that Pds5 can be involved in loop-extruding cohesin in *S. pombe*, alongside the cohesin core proteins. Similar correlation between the ChIP-seq of Mcd1 and Pds5 is also found in *S. cerevisiae* (Supplementary Fig. 22*B* ). Interestingly, this inference concerning the fission yeast and budding yeast cohesin subunit, Pds5, stands in contrast to the conclusion from a recent single-molecule study [43] concerning cohesin in vertebrates. Specifically, Reference [43] found that cohesin complexes containing Pds5, instead of Nipbl, are unable to extrude loops. Other studies also found that Pds5 restricts DNA loop extrusion by cohesin in both budding yeast [42, 64] and vertebrates [41]. In light of these findings, an attractive explanation for the observed correlations between the ChIP-seq of Mcd1 and Pds5 in *S. pombe* and *S. cerevisiae* is that Pds5 could act like a boundary element that stalls cohesin complexes upon contact, thus achieving colocalization with cohesin.

Additionally, as noted above, in interphase *S. pombe* the ChIP-seq signal of the cohesin loader, Mis4, is uncorrelated with the Psc3 ChIP-seq signal in non-centromeric regions (Supplementary Fig. 21*A*), suggesting that Mis4 is, at most, a very transient component of cohesin in *S. pombe*. Such a correlation between the ChIP-seq of Scc2 (counterpart of *S. pombe*’s Mis4) and the cohesin core (Mcd1) is also lacking in mitotic *S. cerevisiae* (Supplementary Fig. 23). However, Reference [82] found that, in addition to its role as a cohesin loader, Scc2 drives expansion of DNA loops *in vivo* in mitotic *S. cerevisiae*. The absence of correlation between the ChIP-seq of Scc2 and cohesin core in mitotic *S. cerevisiae* suggests that the activity of Scc2 in driving DNA loop expansion involves a mechanism other than colocalization or co-translocation with the cohesin core. In contrast to the lack of correlation between the ChIP-seq of Mis4/Scc2 and cohesin in interphase *S. pombe* and mitotic *S. cerevisiae*, there are significant correlations between the ChIP-seq of Nipbl (counterpart of Mis4/Scc2) and the cohesin core protein, Smc1, in humans (Supplementary Fig. 21*G* ). Unsurprisingly, both References [43] and [44] found that Nipbl is an obligate component of the loop-extruding human cohesin complex, in addition to its role as a cohesin loader. Although CCLE has not yet been applied to vertebrates, from a CCLE perspective, the possibility that Nipbl may be required for the loop extrusion process in humans is bolstered by the significant correlations between the ChIP-seq of human Nipbl and the cohesin core (Supplementary Fig. 21G), consistent with Ref. [32]’s hypothesis that Nipbl is involved in loop-extruding cohesin in vertebrates. A recent theoretical model of the molecular mechanism of loop extrusion by cohesin hypothesizes that transient binding by Mis4/Nipbl is essential for permitting directional reversals and therefore for two-sided loop extrusion [46]. Surprisingly, there are significant correlations between Mis4 and Pds5 in *S. pombe* (Supplementary Fig. 21*E* ), indicating Pds5-Mis4 association, outside of the cohesin core complex; however, similar correlations are lacking between Scc2 and Pds5 in *S. cerevisiae* (Supplementary Fig. 22).

Beyond yeast, because the CCLE model is agnostic about the identity of any particular boundary element, in principle it extends the LEF model to organisms across the tree of life, including organisms that do not express CTCF. By contrast, prior LEF models have been overwhelmingly limited to vertebrates, which express CTCF and where CTCF is the principal boundary element. Two exceptions, in which the LEF model was applied to non-vertebrates, are Ref. [54], discussed above, and Ref. [83], which models the Hi-C map of the prokaryote, *Bacillus subtilis*, on the basis of condensin loop extrusion with gene-dependent barriers. In future work, it will be interesting to explore the applicability of CCLE model to other model organisms, that exhibit TADs, including, for example, *Drosophila melanogaster, Arabidopsis thaliana, Oryza sativa, Caenorhabditis elegans*, and *Caulobacter crescentus*, as well as to vertebrates.

## Methods

### Gillespie-type loop extrusion simulations

For interphase *S. pombe* (meiotic and mitotic *S. cerevisiae*), we represent 1.2 Mb (600 kb) regions of the chromatin polymer as an array of 1200 discrete lattice sites, each comprising 1000 bp (500 bp), and simulate chromatin loop configurations using a Gillespie-type algorithm applied to the LEFs, populating these lattice sites, similarly to Refs. [27, 29]. Because the simulations are of Gillespie-type, the time separations between successive events are exponentially-distributed, while which event is realized occurs with a probability that is proportional to the rate of that event. LEFs are modeled as objects with two anchors, which initially bind to empty, adjacent chromatin lattice sites with a binding probability, that is uniformly distributed across the genome. Upon binding, a LEF’s anchors translocate stochastically and independently in opposite directions at rates specified by position-dependent loop extrusion rates determined from experimental cohesin ChIP-seq data. Only outward steps, that grow loops, are permitted. Translocation of a LEF anchor into a lattice site, that is already occupied by another LEF’s anchor, is forbidden. LEFs dissociate at a constant rate, dissipating the corresponding loop. In our simulations, after unbinding, a LEF immediately rebinds to an empty pair of neighboring sites, maintaining a constant number of bound LEFs. Neither simulated contact probabilities nor simulated mean occupancy probabilities depend on the time scale of loop extrusion. Therefore, the results of our simulations for chromatin-chromatin contacts and LEF occupancies depend on the mean LEF processivity, which is the ratio of the mean LEF extrusion rate and the dissociation rate, and which therefore is the appropriate fitting parameter. In other words, the LEF dissociation rate (inverse of lifetime) can be arbitrary so long as the processivity remains unchanged by adjusting the extrusion rate accordingly. In practice, however, we set the LEF dissociation rate to 5 × 10^−4^ time-unit^−1^ (equivalent to a lifetime of 2000 time-units), and the nominal LEF extrusion rate (aka *ρL/τ* , see Supplementary Methods) can be determined from the given processivity.

### ChIP-seq to occupancy probability

The *S. pombe* (*S. cerevisiae*) experimental cohesin ChIP-seq data is averaged over each 1000-bp-sized (500-bp-sized) lattice site to yield the ChIP-seq signal, *C*_*n*_, for each chromatin lattice site, *n*. To convert Psc3 ChIP-seq data at lattice site *n, C*_*n*_, to the occupancy probability, *P*_*n*_, we write

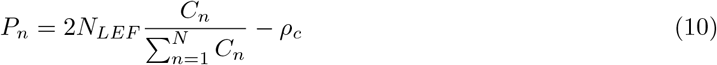

where *ρ*_*c*_ is the cohesive cohesin density (number per chromatin lattice site), *N* is the total number of lattice sites, and *N*_*LEF*_ is the number of LEF simulated in the region represented by *N* lattice sites. The factor of 2 is because one LEF has two anchors and occupies two lattice sites. Although we call *ρ*_*c*_ “cohesive cohesin density”, it is important to note that any experimental backgrounds or shifts in the published ChIP-seq data are also subsumed into this parameter. The excellent experiment-simulation agreement justifies the assumption of a uniform distribution of cohesive cohesins *a posteriori*.

### Modeling the chromatin polymer inside the nucleus

To model chromatin-chromatin contacts, we treat chromatin inside the cell nucleus as a Gaussian polymer in spherical confinement. For a Gaussian polymer in the continuous limit, the probability density, *p*(*r, θ, ϕ, n*), that a genomic locus *n* along the polymer is positioned at coordinates (*r, θ, ϕ*) follows the diffusion equation, *∂p/∂n* = (*l*^2^*/*6)∇2*p*, where *l* is the Kuhn length, which is twice the persistence length. Spherical confinement within a nuclear radius *a* can be enforced by finding solutions to the diffusion equation either with reflecting or absorbing boundary conditions at *r* = *a*. As discussed in Refs. [84–86], reflecting boundary conditions correspond to an attractive surface-polymer interaction given by *ϵ* = −*k*_*B*_*T* log (6*/*5), while absorbing boundary conditions correspond to zero polymer-surface interaction. Comparison of chromatin mean contact probability, *P*(*s*), using reflecting and absorbing boundary conditions is presented in Supplementary Fig. 10. To crudely account for “Rabl configurations”, in which chromatin is known to be attached to the nuclear envelope [87, 88], we chose to use the solution for reflecting boundary conditions to calculate the self-contact probability between any two points along the chromatin polymer. Calculation of the corresponding self-contact probability for two locations with genomic separation, *n*, is presented in detail in Supplementary Methods. A similar method is also adopted in Ref. [89]. To then incorporate loops we replace the actual genomic separation (*n*) between the two loci of interest by their *effective* genomic separation (*n*_*eff*_ ), which is also discussed in detail in Supplementary Methods. In brief, *n*_*eff*_ is the backbone length between the two locations of interest modified by the contribution of any loops that contain the two locations. Our approximate, analytic approach to polymer self-contacts may be justified *a posteriori* on the basis of the excellent data-simulation agreement apparent in Figs. 1 and 2.

Because of the increased compaction of meiotic and mitotic chromosomes, compared to interphase chromosomes, we can expect volume exclusion to play a more prominent role in meiotic and mitotic nuclei than in interphase nuclei. In order to account for volume exclusion in meiotic and mitotic *S. cerevisiae*, we introduce an additional empirical factor, 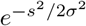, which reduces the probability of contacts with large genomic separations. This factor is characterized by an additional fitting parameter, namely *σ*, which represents a genomic distance scale, beyond which chromatin-chromatin contacts are reduced because of volume exclusion. Inclusion of this factor leads to improved agreement between experimental and simulated *P*(*s*) curves at longer length scales (≳ 50 kb) (Figs. 5*F* and 6*E*), but has little effect on simulated Hi-C maps at shorter length scales (≲ 50 kb).

### Simulated Hi-C contact map generation

To generate simulated Hi-C contact maps, for each genomic region and each set of parameter values, we performed 200 independent loop extrusion simulations, each comprised of 100000 LEF events. In our simulations, loops achieve a dynamic steady state well within 15000 LEF events, as gauged by convergence of the radius of gyration of chromatin to its steady-state value [90] (Supplementary Fig. 24*A*). We also examined two loop size distributions from two different simulation periods: each distribution consists of 1000 data points, equally separated in time, one between LEF event 15000 and 35000, and the other between LEF event 80000 and 100000. The two distributions are within-errors identical (Supplementary Fig. 24*B* ), suggesting that the loop extrusion steady state is well achieved within 15000 LEF events. To ensure that only steady-state loop configurations are included in our simulated Hi-C maps, we discard the first 15000 LEF events. First, to generate a simulated Hi-C contact map for a given loop configuration, we calculate the contact probability for each pair of lattice sites in the given genomic region, using their effective genomic distance, *n*_*eff*_ , and the functional form of self-contact probability for a polymer inside a sphere (Supplementary Methods, Eq. S36). Then, for each of the 200 independent loop extrusion simulation, we create a time-averaged Hi-C map by averaging together 100 simulated Hi-C contact maps, corresponding to 100 simulated loop configurations that are evenly-distributed in time-unit across the simulation time period after the initial 15000 events of non-steady-state period. Finally, we average together the time-averaged Hi-C maps from all 200 simulations to produce an ensemble-averaged Hi-C map. Because the CCLE simulations are performed using a finer resolution than the resolution of experimental Hi-C maps, our ensemble-averaged Hi-C contact maps are binned (block-averaged) to match the resolution of the corresponding experimental Hi-C. The simulated Hi-C maps of interphase *S. pombe* and meiotic *S. cerevisiae* are scaled so that the mean contact probability of each simulated Hi-C map along its second diagonal is equal to the mean contact probability along the second diagonal of the corresponding experimental Hi-C map, while the simulated Hi-C map of mitotic *S. cerevisiae* is scaled so that the mean contact probabilities of the simulated and the corresponding experimental Hi-C maps along the third diagonal are equal. In other words, we require *P*_sim_(20 *kb*) = *P*_exp_(20 *kb*) for interphase *S. pombe, P*_sim_(4 *kb*) = *P*_exp_(4 *kb*) for meiotic *S. cerevisiae*, and *P*_sim_(1.5 *kb*) = *P*_exp_(1.5 *kb*) for mitotic *S. cerevisiae*. The simulated Hi-C maps so-obtained are compared to experimental maps in Fig. 1, 5, 6 and Supplementary Figs. 1–4,

To assess the noise within our simulated Hi-C maps, we also calculated the MPR and the PCC between the averages of two sets of 200 independent, time-averaged Hi-C simulations, for each case of *S. pombe*, meiotic *S. cerevisiae* and mitotic *S. cerevisiae*, giving MPR values of 1.0149, 1.0179, and 1.0121, and PCC values of 0.9834, 0.9924, and 0.9988, respectively. Evidently, these values are close to unity, indicating that our final simulated maps in all presented cases accurately represent the CCLE model with little noise.

## Code availability

Codes used for CCLE simulation and data analysis of ChIP-seq and Hi-C maps are freely and publicly available at https://github.com/bigpaul97/CCLE. All studied experimental data are included and cited in the article and/or Supplementary Information.

## Acknowledgements

This research was supported by NSF EFRI CEE Award EFMA-1830904, NSF PoLS 2412859, and Yale’s Integrated Program Graduate Program in Physical and Engineering Biology (PEB).

## References

[1] Erez Lieberman-Aiden, Nynke L. van Berkum, Louise Williams, Maxim Imakaev, Tobias Ragoczy, Agnes Telling, Ido Amit, Bryan R. Lajoie, Peter J. Sabo, Michael O. Dorschner, Richard Sandstrom, Bradley Bernstein, M. A. Bender, Mark Groudine, Andreas Gnirke, John Stamatoyannopoulos, Leonid A. Mirny, Eric S. Lander, and Job Dekker. Comprehensive mapping of long-range interactions reveals folding principles of the human genome. Science, 326(5950):289–293, 2009.

[2] Nynke L. van Berkum, Erez Lieberman-Aiden, Louise Williams, Maxim Imakaev, Andreas Gnirke, Leonid A. Mirny, Job Dekker, and Eric S. Lander. Hi-C: A method to study the three-dimensional architecture of genomes. Journal of Visualized Experiments : JoVE, 2010.

[3] Jesse R Dixon, Siddarth Selvaraj, Feng Yue, Audrey Kim, Yan Li, Yin Shen, Ming Hu, Jun S Liu, and Bing Ren. Topological domains in mammalian genomes identified by analysis of chromatin interactions. Nature, 485(7398):376–380, 2012.

[4] Jesse R Dixon, David U Gorkin, and Bing Ren. Chromatin domains: the unit of chromosome organization. Molecular Cell, 62(5):668–680, 2016.

[5] Tom Sexton, Eitan Yaffe, Ephraim Kenigsberg, Frédéric Bantignies, Benjamin Leblanc, Michael Hoichman, Hugues Parrinello, Amos Tanay, and Giacomo Cavalli. Three-dimensional folding and functional organization principles of the Drosophila genome. Cell, 148(3):458–472, 2012.

[6] Takeshi Mizuguchi, Geoffrey Fudenberg, Sameet Mehta, Jon-Matthew Belton, Nitika Taneja, Hernan Diego Folco, Peter FitzGerald, Job Dekker, Leonid Mirny, Jemima Barrowman, and Shiv I. S. Grewal. Cohesin-dependent globules and heterochromatin shape 3D genome architecture in S. pombe. Nature, 516(7531):432–435, 2014.

[7] Job Dekker. Two ways to fold the genome during the cell cycle: insights obtained with chromosome conformation capture. Epigenetics & Chromatin, 7(1):1–12, 2014.

[8] Job Dekker and Edith Heard. Structural and functional diversity of topologically associating domains. FEBS Letters, 589(20):2877–2884, 2015.

[9] Thomas D Pollard, William C Earnshaw, Jennifer Lippincott-Schwartz, and Graham Johnson. Cell biology E-book. Elsevier Health Sciences, 2016.

[10] Ivana Jerkovic and Giacomo Cavalli. Understanding 3D genome organization by multidisciplinary methods. Nature Reviews Molecular Cell Biology, 22(8):511–528, 2021.

[11] Mariano Barbieri, Mita Chotalia, James Fraser, Liron-Mark Lavitas, Josée Dostie, Ana Pombo, and Mario Nicodemi. Complexity of chromatin folding is captured by the strings and binders switch model. Proceedings of the National Academy of Sciences, 109(40):16173–16178, 2012.

[12] Luca Giorgetti, Rafael Galupa, Elphège P Nora, Tristan Piolot, France Lam, Job Dekker, Guido Tiana, and Edith Heard. Predictive polymer modeling reveals coupled fluctuations in chromosome conformation and transcription. Cell, 157(4):950–963, 2014.

[13] Mariano Barbieri, Sheila Q Xie, Elena Torlai Triglia, Andrea M Chiariello, Simona Bianco, Ines de Santiago, Miguel R Branco, David Rueda, Mario Nicodemi, and Ana Pombo. Active and poised promoter states drive folding of the extended HoxB locus in mouse embryonic stem cells. Nature Structural & Molecular Biology, 24(6):515–524, 2017.

[14] Chris A Brackley, Jill M Brown, Dominic Waithe, Christian Babbs, James Davies, Jim R Hughes, Veronica J Buckle, and Davide Marenduzzo. Predicting the three-dimensional folding of cis-regulatory regions in mammalian genomes using bioinformatic data and polymer models. Genome Biology, 17(1):59, 2016.

[15] Chris A Brackley, James Johnson, Steven Kelly, Peter R Cook, and Davide Marenduzzo. Simulated binding of transcription factors to active and inactive regions folds human chromosomes into loops, rosettes and topological domains. Nucleic Acids Research, 44(8):3503–3512, 2016.

[16] Michele Di Pierro, Bin Zhang, Erez Lieberman Aiden, Peter G Wolynes, and José N Onuchic. Transferable model for chromosome architecture. Proceedings of the National Academy of Sciences, 113(43):12168–12173, 2016.

[17] Andrea M Chiariello, Carlo Annunziatella, Simona Bianco, Andrea Esposito, and Mario Nicodemi. Polymer physics of chromosome large-scale 3D organisation. Scientific Reports, 6(1):29775, 2016.

[18] Michele Di Pierro, Ryan R Cheng, Erez Lieberman Aiden, Peter G Wolynes, and José N Onuchic. De novo prediction of human chromosome structures: Epigenetic marking patterns encode genome architecture. Proceedings of the National Academy of Sciences, 114(46):12126–12131, 2017.

[19] CA Brackley, J Johnson, D Michieletto, AN Morozov, M Nicodemi, PR Cook, and D Marenduzzo. Extrusion without a motor: a new take on the loop extrusion model of genome organization. Nucleus, 9(1):95–103, 2018.

[20] Simona Bianco, Darío G Lupiáñez, Andrea M Chiariello, Carlo Annunziatella, Katerina Kraft, Robert Schöpflin, Lars Wittler, Guillaume Andrey, Martin Vingron, Ana Pombo, Stefan Mundlos, and Mario Nicodemi. Polymer physics predicts the effects of structural variants on chromatin architecture. Nature Genetics, 50(5):662–667, 2018.

[21] Adam Buckle, Chris A Brackley, Shelagh Boyle, Davide Marenduzzo, and Nick Gilbert. Polymer simulations of heteromorphic chromatin predict the 3D folding of complex genomic loci. Molecular Cell, 72(4):786–797, 2018.

[22] Mattia Conte, Luca Fiorillo, Simona Bianco, Andrea M Chiariello, Andrea Esposito, and Mario Nicodemi. Polymer physics indicates chromatin folding variability across single-cells results from state degeneracy in phase separation. Nature Communications, 11(1):3289, 2020.

[23] Tereza Gerguri, Xiao Fu, Yasutaka Kakui, Bhavin S Khatri, Christopher Barrington, Paul A Bates, and Frank Uhlmann. Comparison of loop extrusion and diffusion capture as mitotic chromosome formation pathways in fission yeast. Nucleic Acids Research, 49(3):1294–1312, 2021.

[24] Mattia Conte, Ehsan Irani, Andrea M Chiariello, Alex Abraham, Simona Bianco, Andrea Esposito, and Mario Nicodemi. Loop-extrusion and polymer phase-separation can co-exist at the single-molecule level to shape chromatin folding. Nature Communications, 13(1):4070, 2022.

[25] Elnaz Alipour and John F Marko. Self-organization of domain structures by DNA-loop-extruding enzymes. Nucleic Acids Research, 40(22):11202–11212, 2012.

[26] Adrian L. Sanborn, Suhas S. P. Rao, Su-Chen Huang, Neva C. Durand, Miriam H. Huntley, Andrew I. Jewett, Ivan D. Bochkov, Dharmaraj Chinnappan, Ashok Cutkosky, Jian Li, Kristopher P. Geeting, Andreas Gnirke, Alexandre Melnikov, Doug McKenna, Elena K. Stamenova, Eric S. Lander, and Erez Lieberman Aiden. Chromatin extrusion explains key features of loop and domain formation in wild-type and engineered genomes. Proceedings of the National Academy of Sciences, 112(47):E6456–E6465, 2015.

[27] Geoffrey Fudenberg, Maxim Imakaev, Carolyn Lu, Anton Goloborodko, Nezar Abdennur, and Leonid A Mirny. Formation of chromosomal domains by loop extrusion. Cell Reports, 15(9):2038– 2049, 2016.

[28] Anton Goloborodko, Maxim V Imakaev, John F Marko, and Leonid Mirny. Compaction and segregation of sister chromatids via active loop extrusion. Elife, 5:e14864, 2016.

[29] Anton Goloborodko, John F Marko, and Leonid A Mirny. Chromosome compaction by active loop extrusion. Biophysical Journal, 110(10):2162–2168, 2016.

[30] Johannes Nuebler, Geoffrey Fudenberg, Maxim Imakaev, Nezar Abdennur, and Leonid A Mirny. Chromatin organization by an interplay of loop extrusion and compartmental segregation. Proceedings of the National Academy of Sciences, 115(29):E6697–E6706, 2018.

[31] Edward J Banigan and Leonid A Mirny. Loop extrusion: theory meets single-molecule experiments. Current Opinion in Cell Biology, 64:124–138, 2020.

[32] Iain F Davidson and Jan-Michael Peters. Genome folding through loop extrusion by SMC complexes. Nature Reviews Molecular Cell Biology, 22(7):445–464, 2021.

[33] A Marieke Oudelaar and Douglas R Higgs. The relationship between genome structure and function. Nature Reviews Genetics, 22(3):154–168, 2021.

[34] Geoffrey Fudenberg, Nezar Abdennur, Maxim Imakaev, Anton Goloborodko, and Leonid A Mirny. Emerging evidence of chromosome folding by loop extrusion. In Cold Spring Harbor symposia on quantitative biology, volume 82, pages 45–55. Cold Spring Harbor Laboratory Press, 2017.

[35] Tsung-Han S Hsieh, Assaf Weiner, Bryan Lajoie, Job Dekker, Nir Friedman, and Oliver J Rando. Mapping nucleosome resolution chromosome folding in yeast by micro-c. Cell, 162(1):108–119, 2015.

[36] Tsung-Han S Hsieh, Geoffrey Fudenberg, Anton Goloborodko, and Oliver J Rando. Micro-c xl: assaying chromosome conformation from the nucleosome to the entire genome. Nature methods, 13(12):1009–1011, 2016.

[37] Masae Ohno, Tadashi Ando, David G Priest, Vipin Kumar, Yamato Yoshida, and Yuichi Taniguchi. Sub-nucleosomal genome structure reveals distinct nucleosome folding motifs. Cell, 176(3):520–534, 2019.

[38] Oliver Wiese, Davide Marenduzzo, and Chris A Brackley. Nucleosome positions alone can be used to predict domains in yeast chromosomes. Proceedings of the National Academy of Sciences, 116(35):17307–17315, 2019.

[39] Elisa Oberbeckmann, Kimberly Quililan, Patrick Cramer, and A Marieke Oudelaar. In vitro reconstitution of chromatin domains shows a role for nucleosome positioning in 3d genome organization. Nature Genetics, 56(3):483–492, 2024.

[40] Suhas S.P. Rao, Su-Chen Huang, Brian Glenn St Hilaire, Jesse M. Engreitz, Elizabeth M. Perez, Kyong-Rim Kieffer-Kwon, Adrian L. Sanborn, Sarah E. Johnstone, Gavin D. Bascom, Ivan D. Bochkov, Xingfan Huang, Muhammad S. Shamim, Jaeweon Shin, Douglass Turner, Ziyi Ye, Arina D. Omer, James T. Robinson, Tamar Schlick, Bradley E. Bernstein, Rafael Casellas, Eric S. Lander, and Erez Lieberman Aiden. Cohesin loss eliminates all loop domains. Cell, 171(2):305–320, 2017.

[41] Gordana Wutz, Csilla Várnai, Kota Nagasaka, David A Cisneros, Roman R Stocsits, Wen Tang, Stefan Schoenfelder, Gregor Jessberger, Matthias Muhar, M Julius Hossain, Nike Walther, Birgit Koch, Moritz Kueblbeck, Jan Ellenberg, Johannes Zuber, Peter Fraser, and Jan-Michael Peters. Topologically associating domains and chromatin loops depend on cohesin and are regulated by CTCF, WAPL, and PDS5 proteins. The EMBO Journal, 36(24):3573–3599, 2017.

[42] Lorenzo Costantino, Tsung-Han S Hsieh, Rebecca Lamothe, Xavier Darzacq, and Douglas Koshland. Cohesin residency determines chromatin loop patterns. Elife, 9:e59889, 2020.

[43] Iain F Davidson, Benedikt Bauer, Daniela Goetz, Wen Tang, Gordana Wutz, and Jan-Michael Peters. DNA loop extrusion by human cohesin. Science, 366(6471):1338–1345, 2019.

[44] Yoori Kim, Zhubing Shi, Hongshan Zhang, Ilya J Finkelstein, and Hongtao Yu. Human cohesin compacts DNA by loop extrusion. Science, 366(6471):1345–1349, 2019.

[45] Stefan Golfier, Thomas Quail, Hiroshi Kimura, and Jan Brugués. Cohesin and condensin extrude DNA loops in a cell cycle-dependent manner. Elife, 9:e53885, 2020.

[46] Torahiko L Higashi, Georgii Pobegalov, Minzhe Tang, Maxim I Molodtsov, and Frank Uhlmann. A brownian ratchet model for DNA loop extrusion by the cohesin complex. Elife, 10:e67530, 2021.

[47] Elphège P Nora, Anton Goloborodko, Anne-Laure Valton, Johan H Gibcus, Alec Uebersohn, Nezar Abdennur, Job Dekker, Leonid A Mirny, and Benoit G Bruneau. Targeted degradation of CTCF decouples local insulation of chromosome domains from genomic compartmentalization. Cell, 169(5):930–944, 2017.

[48] Elissavet Kentepozidou, Sarah J Aitken, Christine Feig, Klara Stefflova, Ximena Ibarra-Soria, Duncan T Odom, Maša Roller, and Paul Flicek. Clustered CTCF binding is an evolutionary mechanism to maintain topologically associating domains. Genome Biology, 21:1–19, 2020.

[49] Chin-Tong Ong and Victor G Corces. CTCF: an architectural protein bridging genome topology and function. Nature Reviews Genetics, 15(4):234–246, 2014.

[50] Congmao Wang, Chang Liu, Damian Roqueiro, Dominik Grimm, Rebecca Schwab, Claude Becker, Christa Lanz, and Detlef Weigel. Genome-wide analysis of local chromatin packing in Arabidopsis thaliana. Genome Research, 25(2):246–256, 2015.

[51] Qianli Dong, Ning Li, Xiaochong Li, Zan Yuan, Dejian Xie, Xiaofei Wang, Jianing Li, Yanan Yu, Jinbin Wang, Baoxu Ding, Zhibin Zhang, Changping Li, Yao Bian, Ai Zhang, Ying Wu, Bao Liu, and Lei Gong. Genome-wide Hi-C analysis reveals extensive hierarchical chromatin interactions in rice. The Plant Journal, 94(6):1141–1156, 2018.

[52] Peter Heger, Birger Marin, and Einhard Schierenberg. Loss of the insulator protein CTCF during nematode evolution. BMC Molecular Biology, 10:1–14, 2009.

[53] Anjali Kaushal, Giriram Mohana, Julien Dorier, Isa Özdemir, Arina Omer, Pascal Cousin, Anastasiia Semenova, Michael Taschner, Oleksandr Dergai, Flavia Marzetta, Christian Iseli, Yossi Eliaz, David Weisz, Muhammad Saad Shamim, Nicolas Guex, Erez Lieberman Aiden, and Maria Cristina Gambetta. CTCF loss has limited effects on global genome architecture in Drosophila despite critical regulatory functions. Nature Communications, 12(1):1011, 2021.

[54] Stephanie A Schalbetter, Geoffrey Fudenberg, Jonathan Baxter, Katherine S Pollard, and Matthew J Neale. Principles of meiotic chromosome assembly revealed in S. cerevisiae. Nature Communications, 10(1):4795, 2019.

[55] Samuel Marguerat, Alexander Schmidt, Sandra Codlin, Wei Chen, Ruedi Aebersold, and Jürg Bähler. Quantitative analysis of fission yeast transcriptomes and proteomes in proliferating and quiescent cells. Cell, 151(3):671–683, 2012.

[56] Jean-Michel Arbona, Sébastien Herbert, Emmanuelle Fabre, and Christophe Zimmer. Inferring the physical properties of yeast chromatin through Bayesian analysis of whole nucleus simulations. Genome Biology, 18:1–15, 2017.

[57] Mahipal Ganji, Indra A Shaltiel, Shveta Bisht, Eugene Kim, Ana Kalichava, Christian H Haering, and Cees Dekker. Real-time imaging of DNA loop extrusion by condensin. Science, 360(6384):102– 105, 2018.

[58] Ralf Bundschuh and Terence Hwa. Statistical mechanics of secondary structures formed by random RNA sequences. Physical Review E, 65(3):031903, 2002.

[59] Tammy MK Cheng, Sebastian Heeger, Raphaël AG Chaleil, Nik Matthews, Aengus Stewart, Jon Wright, Carmay Lim, Paul A Bates, and Frank Uhlmann. A simple biophysical model emulates budding yeast chromosome condensation. Elife, 4:e05565, 2015.

[60] Minzhe Tang, Georgii Pobegalov, Hideki Tanizawa, Zhuo A Chen, Juri Rappsilber, Maxim Molodtsov, Ken-ichi Noma, and Frank Uhlmann. Establishment of dsdna-dsdna interactions by the condensin complex. Molecular cell, 83(21):3787–3800, 2023.

[61] M. Levo, J. Raimundo, X. Y. Bing, Z. Sisco, P. J. Batut, S. Ryabichko, T. Gregor, and M. S. Levine. Transcriptional coupling of distant regulatory genes in living embryos. Nature, 605:754–760, 2022.

[62] Masaru Ito, Kazuto Kugou, Jeffrey A Fawcett, Sachiko Mura, Sho Ikeda, Hideki Innan, and Kunihiro Ohta. Meiotic recombination cold spots in chromosomal cohesion sites. Genes to Cells, 19(5):359– 373, 2014.

[63] Judith H.I. Haarhuis, Robin H. van der Weide, Vincent A. Blomen, J. Omar Yáñez-Cuna, Mario Amendola, Marjon S. van Ruiten, Peter H.L. Krijger, Hans Teunissen, René H. Medema, Bas van Steensel, Thijn R. Brummelkamp, Elzo de Wit, and Benjamin D. Rowland. The cohesin release factor WAPL restricts chromatin loop extension. Cell, 169(4):693–707, 2017.

[64] Lise Dauban, Rémi Montagne, Agnès Thierry, Luciana Lazar-Stefanita, Nathalie Bastié, Olivier Gadal, Axel Cournac, Romain Koszul, and Frédéric Beckoüet. Regulation of cohesin-mediated chromosome folding by Eco1 and other partners. Molecular Cell, 77(6):1279–1293, 2020.

[65] Yasutaka Kakui, Adam Rabinowitz, David J Barry, and Frank Uhlmann. Condensin-mediated remodeling of the mitotic chromatin landscape in fission yeast. Nature Genetics, 49(10):1553–1557, 2017.

[66] Matthew Robert Paul, Tovah Elise Markowitz, Andreas Hochwagen, and Sevinç Ercan. Condensin depletion causes genome decompaction without altering the level of global gene expression in saccharomyces cerevisiae. Genetics, 210(1):331–344, 2018.

[67] Yasutaka Kakui, Christopher Barrington, David J Barry, Tereza Gerguri, Xiao Fu, Paul A Bates, Bhavin S Khatri, and Frank Uhlmann. Fission yeast condensin contributes to interphase chromatin organization and prevents transcription-coupled DNA damage. Genome Biology, 21(1):1–25, 2020.

[68] Biswajit Pradhan, Takaharu Kanno, Miki Umeda Igarashi, Mun Siong Loke, Martin Dieter Baaske, Jan Siu Kei Wong, Kristian Jeppsson, Camilla Björkegren, and Eugene Kim. The smc5/6 complex is a dna loop-extruding motor. Nature, 616(7958):843–848, 2023.

[69] Georg A Busslinger, Roman R Stocsits, Petra Van Der Lelij, Elin Axelsson, Antonio Tedeschi, Niels Galjart, and Jan-Michael Peters. Cohesin is positioned in mammalian genomes by transcription, ctcf and wapl. Nature, 544(7651):503–507, 2017.

[70] Edward J Banigan, Wen Tang, Aafke A van den Berg, Roman R Stocsits, Gordana Wutz, Hugo B Brandão, Georg A Busslinger, Jan-Michael Peters, and Leonid A Mirny. Transcription shapes 3d chromatin organization by interacting with loop extrusion. Proceedings of the National Academy of Sciences, 120(11):e2210480120, 2023.

[71] Iain F Davidson, Daniela Goetz, Maciej P Zaczek, Maxim I Molodtsov, Pim J Huis in’t Veld, Florian Weissmann, Gabriele Litos, David A Cisneros, Maria Ocampo-Hafalla, Rene Ladurner, et al. Rapid movement and transcriptional re-localization of human cohesin on DNA. The EMBO journal, 35(24):2671–2685, 2016.

[72] Monika Gullerova and Nick J Proudfoot. Cohesin complex promotes transcriptional termination between convergent genes in s. pombe. Cell, 132(6):983–995, 2008.

[73] Christine K Schmidt, Neil Brookes, and Frank Uhlmann. Conserved features of cohesin binding along fission yeast chromosomes. Genome Biology, 10(5):1–16, 2009.

[74] Armelle Lengronne, Yuki Katou, Saori Mori, Shihori Yokobayashi, Gavin P Kelly, Takehiko Itoh, Yoshinori Watanabe, Katsuhiko Shirahige, and Frank Uhlmann. Cohesin relocation from sites of chromosomal loading to places of convergent transcription. Nature, 430(6999):573–578, 2004.

[75] Earl F Glynn, Paul C Megee, Hong-Guo Yu, Cathy Mistrot, Elcin Unal, Douglas E Koshland, Joseph L DeRisi, and Jennifer L Gerton. Genome-wide mapping of the cohesin complex in the yeast saccharomyces cerevisiae. PLoS Biology, 2(9):e259, 2004.

[76] Maria Ocampo-Hafalla, Sofía Muñoz, Catarina P Samora, and Frank Uhlmann. Evidence for cohesin sliding along budding yeast chromosomes. Open biology, 6(6):150178, 2016.

[77] Kristian Jeppsson, Toyonori Sakata, Ryuichiro Nakato, Stefina Milanova, Katsuhiko Shirahige, and Camilla Björkegren. Cohesin-dependent chromosome loop extrusion is limited by transcription and stalled replication forks. Science Advances, 8(23):eabn7063, 2022.

[78] Rafal Ciosk, Masaki Shirayama, Anna Shevchenko, Tomoyuki Tanaka, Attila Toth, Andrej Shevchenko, and Kim Nasmyth. Cohesin’s binding to chromosomes depends on a separate complex consisting of Scc2 and Scc4 proteins. Molecular Cell, 5(2):243–254, 2000.

[79] Tatsuro S Takahashi, Pannyun Yiu, Michael F Chou, Steven Gygi, and Johannes C Walter. Recruitment of Xenopus Scc2 and cohesin to chromatin requires the pre-replication complex. Nature Cell Biology, 6(10):991–996, 2004.

[80] V. Makrantoni and A. Marston. Cohesin and chromosome segregation. Current Biology, 28:R688– R693, 2018.

[81] Jan-Michael Peters, Antonio Tedeschi, and Julia Schmitz. The cohesin complex and its roles in chromosome biology. Genes & Development, 22(22):3089–3114, 2008.

[82] Nathalie Bastié, Christophe Chapard, Lise Dauban, Olivier Gadal, Frederic Beckouet, and Romain Koszul. Smc3 acetylation, pds5 and scc2 control the translocase activity that establishes cohesin-dependent chromatin loops. Nature structural & molecular biology, 29(6):575–585, 2022.

[83] Hugo B. Brandão, Payel Paul, Aafke A. van den Berg, and Leonid A. Mirny. RNA polymerases as moving barriers to condensin loop extrusion. Proc. Nat. Acad. Sci., 116:20489–20499, 2019.

[84] Edmund A DiMarzio. Proper accounting of conformations of a polymer near a surface. The Journal of Chemical Physics, 42(6):2101–2106, 1965.

[85] Edward F Casassa. Equilibrium distribution of flexible polymer chains between a macroscopic solution phase and small voids. Journal of Polymer Science Part B: Polymer Letters, 5(9):773–778, 1967.

[86] Giuseppe Allegra and Emanuele Colombo. Polymer chain between attractive walls: A second-order transition. The Journal of Chemical Physics, 98(9):7398–7404, 1993.

[87] Takeshi Mizuguchi, Jemima Barrowman, and Shiv IS Grewal. Chromosome domain architecture and dynamic organization of the fission yeast genome. FEBS Letters, 589(20):2975–2986, 2015.

[88] Sarah M Schreiner, Peter K Koo, Yao Zhao, Simon GJ Mochrie, and Megan C King. The tethering of chromatin to the nuclear envelope supports nuclear mechanics. Nature Communications, 6(1):7159, 2015.

[89] Edward J Banigan, Aafke A van den Berg, Hugo B Brandão, John F Marko, and Leonid A Mirny. Chromosome organization by one-sided and two-sided loop extrusion. Elife, 9:e53558, 2020.

[90] Tianyu Yuan, Hao Yan, Mary Lou P Bailey, Jessica F Williams, Ivan Surovtsev, Megan C King, and Simon GJ Mochrie. Effect of loops on the mean-square displacement of rouse-model chromatin. Physical Review E, 109(4):044502, 2024.

