## Supplementary file for "Cohesin distribution alone predicts chromatin organization in yeast via conserved-current loop extrusion"

August 7, 2024

### Methods

#### CCLE model optimization

The CCLE model has four fitting parameters with which to optimize the agreement between experimental and CCLE-simulated *S. pombe* Hi-C maps, namely the LEF density, $\rho$, the mean LEF processivity, $L$ ($L = 2v \times \tau$, where $v$ is the averaged extrusion rate of an unobstructed LEF anchor and $\tau$ is the mean lifetime of a LEF), the chromatin persistence length, and the cohesive cohesin density (ChIP-seq data background), $\rho_c$. To empirically model polymer volume exclusion in the more compact chromosomes of meiotic and mitotic *S. cerevisiae*, we introduce an additional parameter, which is the standard deviation, $\sigma$, of an *ad hoc* Gaussian function, which causes a decrease of contact probability at large genomic separations.

An initial set of parameter values, which appeared by eye to describe experimental Hi-C maps well, was manually found by trial-and-error. To refine these initial guesses, we considered two objective functions: (1) The MPR, as described in the main text, that compares the simulated and experimental Hi-C maps for genomic separation between 20 kb and 120 kb in *S. pombe* (4 and 64 kb in meiotic *S. cerevisiae*, and 1 and 45 kb in mitotic *S. cerevisiae*); and (2) a version of the MPR that compares experimental and simulated $P(s)$ curves for genomic separations from 20 kb to 500 kb in *S. pombe* (4 to 120 kb in meiotic *S. cerevisiae*, and 1.5 to 100 kb in mitotic *S. cerevisiae*), namely

$$\mathrm{MPR}_{P(s)} = \frac{1}{N}\sum_{n}^{N} e^{|\log(E_{P(n)})-\log(S_{P(n)})|}, \tag{1}$$

where $E_{P(n)}$ and $S_{P(n)}$ are the experimental and simulated mean contact probabilities, respectively, for genomic separation $n$, and $N$ is the total number of equally-spaced genomic separations considered. For *S. pombe*, optimization based on MPR score is performed on the 1 Mb-sized regions given in Table 1 of the main text. For meiotic and mitotic *S. cerevisiae*, optimization based on MPR score is performed on the 500 kb and 100 kb-sized regions, respectively, given in Table 1 of the main text.

We found the minimum value of the MPR for each parameter in sequence (in the order of cohesive cohesin density, LEF density, processivity, persistence length, and $\sigma$), by scanning through each parameter space in steps of equal size around the initial guess until a local minimum was detected. For both *S. pombe* and *S. cerevisiae*, the step sizes were $\sim$0.006 kb$^{-1}$ ($\sim$0.018 kb$^{-1}$ for *S. cerevisiae*) for the cohesive cohesin density, 0.00167 kb$^{-1}$ for the LEF density, $\sim$ 1.2 kb for the processivity, and 5 nm for the persistence length. For *S. cerevisiae*, the step size in $\sigma$ was 1 kb.

Our method for determining best-fit parameter values and their corresponding errors is informed by the observation that the deviation from unity of the MPRs for two separate sets of simulation data (each set consists of 200 independent loop extrusion simulations), compared to the same experimental Hi-C data, differ from each other by about 0.7%. Accordingly, We judge that in order for two MPRs to distinguish a better fit from a worse fit, their deviation from unity should should differ by 2% or more.

Depending on different sensitivities of the two objective functions to perturbations in different model parameters, one of the two objective functions was chosen to be used to optimize each model parameter for different organisms and genomic regions. For the LEF density and the cohesive condensin density, and for the LEF processivity, except for Chr 13 of meiotic *S. cerevisiae*, the range of parameter values was sufficiently broad that, at the ends of the parameter value range, the deviation of the MPR from 1 is larger than 1.02 times the deviation of the minimum MPR from unity, *i.e.* $\mathrm{MPR}_{\mathrm{ends}} - 1 > 1.02 \times (\mathrm{MPR}_{\min} - 1)$. In these cases, we selected the best-fit parameters to correspond to the minimum of the MPR. However, for the persistence length, and for the LEF processivity and $\sigma$ of meiotic *S. cerevisiae*, even at the extremes of the ranges of these parameters, the deviation of the MPR from 1 remains less than or barely exceeds 1.02 times the deviation of the minimum MPR from unity. However, $P(s)$ curves are more sensitive to these parameters than Hi-C maps, so that the deviations of $\text{MPR}_{P(s)}$ from 1 at the extremes of the parameter range do exceed 1.02 times the deviation of the minimum $\text{MPR}_{P(s)}$ from 1. Accordingly, in this circumstance, we select the best-fit parameter values that correspond to the minimum of $\text{MPR}_{P(s)}$. For all parameters of Chr 10 of mitotic *S. cerevisiae*, since the Hi-C data is noisier, the MPR is not an effective metric in this case. Instead, we select best-fit parameter values that correspond to the minimum of $\text{MPR}_{P(s)}$.

To estimate the error of each best-fit parameter for *S. pombe* and meiotic *S. cerevisiae*, we carry out least-mean-squares fits of a parabola centered on the MPRs of the minimum-MPR point and its two neighbors versus parameter value. We then assign the parameter value error to be one-half of the range of parameter values where the parabola is less than $1.02 \times (\text{MPR}_{\text{min}} - 1) + 1$. For parameters of mitotic *S. cerevisiae*, we carry out the same error estimation process as described above, but based on $\text{MPR}_{P(s)}$ instead.

#### Solution to position-dependent loop extrusion velocity in CCLE

By envisioning loop-extruding LEFs to constitute probability currents that flow between chromatin lattice sites, we can write down master equations for the probabilities, $R_n$ and $L_n$, that chromatin site $n$ is occupied by right-moving or left-moving LEF anchors:

$$\frac{dR_n}{dt} = V_{n-1}R_{n-1}(1 - P_n) - V_nR_n(1 - P_{n+1}) + A_n - D_n, \tag{2}$$

$$\frac{dL_n}{dt} = U_{n+1}L_{n+1}(1 - P_n) - U_nL_n(1 - P_{n-1}) + a_n - d_n, \tag{3}$$

where $V_n$ is the rate at which right-moving LEF anchors step from site $n$ to site $n+1$, $U_n$ is the rate at which left-moving LEF anchors step from site $n$ to site $n-1$, $P_n = R_n + L_n$ is the probability that lattice site $n$ is occupied by either a left- or right-moving LEF anchor, $A_n$ and $D_n$ ($a_n$ and $d_n$) are the association and dissociation currents of right-moving (left-moving) LEF anchors at site $n$, respectively.

At steady-state, the left-hand sides of EQ. 2 and EQ. 3 both equal zero, so that

$$0 = V_{n-1}R_{n-1}(1 - P_n) - V_nR_n(1 - P_{n+1}) + A_n - D_n, \tag{4}$$

$$0 = U_{n+1}L_{n+1}(1 - P_n) - U_nL_n(1 - P_{n-1}) + a_n - d_n, \tag{5}$$

EQ. 4 and EQ. 5 are exact, but as-is they are intractable. To make further progress, we first assume that the difference between the association and dissociation terms at each lattice site is small, compared to the flow terms, and therefore can be neglected, *i.e.* we assume that $a_n - d_n \simeq 0$ and $A_n - D_n \simeq 0$. Next, we make a mean-field-type approximation, that right- and left-moving LEF anchors are uncoupled. We also assume that $R_n = L_n = \frac{1}{2}P_n$. It follows that

$$0 = V_{n-1}P_{n-1}(1 - P_n) - V_n P_n(1 - P_{n+1}), \tag{6}$$

and

$$0 = U_{n+1}P_{n+1}(1 - P_n) - U_n P_n(1 - P_{n-1}). \tag{7}$$

In our calculation, we impose periodic boundary conditions, so that

$$0 = V_N P_N(1 - P_1) - V_1 P_1(1 - P_2), \tag{8}$$

and

$$0 = U_1 P_1(1 - P_N) - U_N P_N(1 - P_{N-1}), \tag{9}$$

given that there are a total of $N$ lattice sites.

The systems of equations represented by Eq. 6 and 8, on the one hand, and Eq. 7 and 9, on the other have solutions

$$V_n = \frac{r}{P_n(1 - P_{n+1})}, \tag{10}$$

and

$$U_n = \frac{r}{P_n(1 - P_{n-1})}, \tag{11}$$

where $r$ is a constant rate.

In the absence of obstacles, the actual rate, $V_n$ or $U_n$, is the rate, $r$, scaled by the occupancy level, $P_n$, which depends on the density of LEF anchors, $2\rho$. We relate the actual mean loop extrusion rate of an isolated LEF, $v$, and the rate, $r$, by $v = r/(2\rho)$. So that Eq. 10 becomes

$$V_n = \frac{2\rho v}{P_n(1 - P_{n+1})}, \tag{12}$$

and similarly for Eq. 11. We also have $v = L/(2\tau)$, where $L$ is the mean processivity of isolated LEFs and $\tau$ is the mean lifetime of LEFs. Therefore,

$$V_n = \frac{\rho L/\tau}{P_n(1 - P_{n+1})}, \tag{13}$$

which are the final expressions of $V_n$ and $U_n$ given in the main text.

#### Self-contact probability of a Gaussian polymer in spherical confinement

The three-dimensional conformation of a Gaussian polymer can be represented as the trajectory of a three-dimensional random walk. Accordingly, to model chromatin inside the cell nucleus, we consider a three-dimensional random walk inside a sphere of radius $a$, subject to reflecting boundary conditions at $r = a$. This choice corresponds to an attractive polymer-surface potential, crudely mimicking chromatin-nuclear envelope attachments, *i.e.* Rabl configurations, that are known to be a feature of yeast chromosomes. To determine the probability that two distal genomic loci contact each other, first, we seek the conditional probability , $P(r, \theta, \phi, t|R, \alpha, \beta, 0)$, that the walk arrives at ($r$, $\theta$, $\phi$) at time $t$, given that it starts at ($R$, $\alpha$, $\beta$) at time $t = 0$. The equation governing $P = P(r, \theta, \phi, t|R, \alpha, \beta, 0)$ is the diffusion equation:

$$\frac{\partial P}{\partial t} = D\nabla^2 P, \tag{14}$$

where

$$\nabla^2 = \frac{1}{r^2}\frac{\partial}{\partial r}(r^2\frac{\partial}{\partial r}) + \frac{1}{r^2\sin\theta}\frac{\partial}{\partial\theta}(\sin\theta\frac{\partial}{\partial\theta}) + \frac{1}{r^2\sin^2\theta}\frac{\partial^2}{\partial\phi^2}. \tag{15}$$

Assuming separation of variables, *i.e.* that $P = \rho(r)\Theta(\theta)\Phi(\phi)T(t)$, we find that

$$\frac{d^2\Phi}{d\phi^2} = -m^2\Phi, \tag{16}$$

$$\frac{1}{\sin\theta}\frac{d}{d\theta}(\sin\theta\frac{d\Theta}{d\theta}) + (l(l+1) - \frac{m^2}{\sin^2\theta})\Theta = 0, \tag{17}$$

$$\frac{d}{dr}(r^2\frac{d\rho}{dr}) + [\lambda^2 r^2 - l(l+1)]\rho = 0, \tag{18}$$

$$\frac{dT}{dt} = -D\lambda^2 T. \tag{19}$$

The eigenfunctions of Eq. 16 are $\Phi(\phi) = Ae^{im\phi}$. The requirement that $\Phi(0) = \Phi(2\pi)$ then requires that $m$ is an integer. The solutions to Eq. 17 are the associated Legendre polynomials, $\Theta(\theta) = P_l^m(\cos\theta)$ with integer values of $l$, where $-l < m < l$. The products of these angular eigenfunctions, namely $\Phi(\phi)\Theta(\theta)$, are the spherical harmonics, $Y_l^m(\theta, \phi)$. Solutions to Eq. 18 are spherical Bessel functions of the first kind, $\rho(r) = j_l(\lambda r)$. Since the reflecting boundary condition requires that $\rho'(a) = 0$, it follows that $j_l'(\lambda a) = 0$, *i.e.* $\lambda a = x_{l,p}$, where $x_{l,p}$ is the p-th zero of $j_l'$. Thus, the eigenfunctions of Eq. 18, subject to the reflecting boundary condition at $r = a$, may be written as

$$\rho(r) = j_l(x_{l,p}\frac{r}{a}). \tag{20}$$

For different values of $p$, these functions are orthogonal:

$$\int_0^a j_l(x_{l,p}\frac{r}{a})j_l(x_{l,q}\frac{r}{a})r^2\,dr = \frac{\pi a^3}{4x_{l,p}}(J_{l+\frac{1}{2}}(x_{l,p})^2 - J_{l-\frac{1}{2}}(x_{l,p})J_{l+\frac{3}{2}}(x_{l,p}))\delta_{p,q}$$
$$= \frac{a^3}{2}(j_l(x_{l,p})^2 - j_{l-1}(x_{l,p})j_{l+1}(x_{l,p}))\delta_{p,q}. \qquad (21)$$

The solutions to Eq. 19 are

$$T(t) = e^{\frac{-Dx_{l,p}^2}{a^2}t}. \qquad (22)$$

Thus, we can write down the general solution as a superposition:

$$P(r,\theta,\phi,t|R,\alpha,\beta,0) = \sum_{p=1}^{\infty}\sum_{l=0}^{\infty}\sum_{m=-l}^{l} C_{l,m,p}j_l(x_{l,p}\frac{r}{a})Y_l^m(\theta,\phi)e^{\frac{-Dx_{l,p}^2}{a^2}t}. \qquad (23)$$

The initial condition is

$$P(r,\theta,\phi,0|R,\alpha,\beta,0) = \frac{1}{r^2\sin\theta}\delta(r-R)\delta(\theta-\alpha)\delta(\phi-\beta), \qquad (24)$$

so

$$\frac{1}{r^2\sin\theta}\delta(r-R)\delta(\theta-\alpha)\delta(\phi-\beta) = \sum_{p=1}^{\infty}\sum_{l=0}^{\infty}\sum_{m=-l}^{l} C_{l,m,p}j_l(x_{l,p}\frac{r}{a})Y_l^m(\theta,\phi). \qquad (25)$$

Multiplying both sides by $Y_{l_1}^{m_1 *}(\theta,\phi)$ and integrating over angular variables, we find

$$\int_0^{2\pi} d\phi \int_0^{\pi} \sin\theta\, d\theta \frac{1}{r^2\sin\theta}\delta(r-R)\delta(\theta-\alpha)\delta(\phi-\beta)Y_{l_1}^{m_1 *}(\theta,\phi) = \frac{1}{r^2}\delta(r-R)Y_{l_1}^{m_1 *}(\alpha,\beta)$$
$$= \sum_{p=1}^{\infty}\sum_{l=0}^{\infty}\sum_{m=-l}^{l} C_{l,m,p}j_l(x_{l,p}\frac{r}{a})\int_0^{2\pi} d\phi \int_0^{\pi} d\theta \sin\theta Y_l^m(\theta,\phi)Y_{l_1}^{m_1 *}(\theta,\phi)$$
$$= \sum_{p=1}^{\infty} C_{l_1,m_1,p}j_{l_1}(x_{l_1,p}\frac{r}{a}), \qquad (26)$$

where we used the orthonormality of the spherical harmonics. Next, re-writing $l_1$ and $m_1$ as $l$ and $m$, respectively, and carrying out the radial integral utilizing Eq. 21, we find

$$Y_l^{m*}(\alpha,\beta)\int_0^a r^2\frac{1}{r^2}\delta(r-R)j_l(x_{l,q}\frac{r}{a}) = Y_l^{m*}(\alpha,\beta)j_l(x_{l,q}\frac{R}{a})$$
$$= \sum_{p=1}^{\infty} C_{l,m,p}\int_0^a r^2 j_l(x_{l,p}\frac{r}{a})j_l(x_{l,q}\frac{r}{a})\,dr = \frac{a^3}{2}C_{l,m,q}(j_l(x_{l,q})^2 - j_{l-1}(x_{l,q})j_{l+1}(x_{l,q})), \qquad (27)$$

*i.e.*

$$C_{l,m,q} = \frac{2Y_l^{m*}(\alpha,\beta)j_l(x_{l,q}\frac{R}{a})}{a^3(j_l(x_{l,q})^2 - j_{l-1}(x_{l,q})j_{l+1}(x_{l,q}))} \qquad (28)$$

Inserting EQ. 28 into eq. 23, we arrive at

$$P(r,\theta,\phi,t|R,\alpha,\beta,0) = \frac{2}{a^3}\sum_{p=1}^{\infty}\sum_{l=0}^{\infty}\sum_{m=-l}^{l}\frac{j_l(x_{l,p}\frac{R}{a})j_l(x_{l,p}\frac{r}{a})}{(j_l(x_{l,p})^2 - j_{l-1}(x_{l,p})j_{l+1}(x_{l,p}))}Y_l^{m*}(\alpha,\beta)Y_l^m(\theta,\phi)e^{\frac{-Dx_{l,p}^2}{a^2}t}, \tag{29}$$

which is the desired solution of the diffusion equation inside a sphere of radius $a$ subject to reflecting boundary conditions and the initial condition that $(r,\theta,\phi) = (R,\alpha,\beta)$.

We can find the return probability at time $t$ as

$$P(R,\alpha,\beta,t|R,\alpha,\beta,0) = \frac{2}{a^3}\sum_{p=1}^{\infty}\sum_{l=0}^{\infty}\sum_{m=-l}^{l}\frac{j_l(x_{l,p}\frac{R}{a})j_l(x_{l,p}\frac{R}{a})}{(j_l(x_{l,p})^2 - j_{l-1}(x_{l,p})j_{l+1}(x_{l,p}))}Y_l^{m*}(\alpha,\beta)Y_l^m(\alpha,\beta)e^{\frac{-Dx_{l,p}^2}{a^2}t}. \tag{30}$$

The spherical harmonic addition theorem is

$$\sum_{m=-l}^{l}Y_l^{m*}(\alpha,\beta)Y_l^m(\alpha,\beta) = \frac{2l+1}{4\pi}. \tag{31}$$

Thus, EQ. 30 simplifies to

$$P(R,\alpha,\beta,t|R,\alpha,\beta,0) = \frac{1}{2\pi a^3}\sum_{p=1}^{\infty}\sum_{l=0}^{\infty}\frac{j_l(x_{l,p}\frac{R}{a})j_l(x_{l,p}\frac{R}{a})}{(j_l(x_{l,p})^2 - j_{l-1}(x_{l,p})j_{l+1}(x_{l,p}))}(2l+1)e^{\frac{-Dx_{l,p}^2}{a^2}t}, \tag{32}$$

which sensibly does not depend on the initial angular coordinates. Finally, we average over all possible initial positions, using Eq. 21 again, with the result that

$$\begin{aligned}
&\langle P(R,\alpha,\beta,t|R,\alpha,\beta,0)\rangle \\
&= \frac{3}{4\pi a^3}\int_0^a R^2\,dR\int_0^{\pi}\sin\alpha\,d\alpha\int_0^{2\pi}d\beta P(R,\alpha,\beta,t|R,\alpha,\beta,0) \\
&= \frac{3}{4\pi a^3}\frac{1}{2\pi a^3}4\pi\sum_{p=1}^{\infty}\sum_{l=0}^{\infty}\frac{(2l+1)e^{\frac{Dx_{l,p}^2}{a^2}t}}{(j_l(x_{l,p})^2 - j_{l-1}(x_{l,p})j_{l+1}(x_{l,p}))}\int_0^a R^2 j_l(x_{l,p}\frac{R}{a})j_l(x_{l,p}\frac{R}{a})\,dR \\
&= \frac{3}{4\pi a^3}\frac{1}{2\pi a^3}4\pi\frac{a^3}{2}\sum_{p=1}^{\infty}\sum_{l=0}^{\infty}(2l+1)e^{\frac{Dx_{l,p}^2}{a^2}t} \\
&= \frac{3}{4\pi a^3}\sum_{p=1}^{\infty}\sum_{l=0}^{\infty}(2l+1)e^{\frac{Dx_{l,p}^2}{a^2}t}.
\end{aligned} \tag{33}$$

The first zero of $j_0'(z)$, *i.e.* $x_{0,1}$, occurs at $z = 0$, so that Eq. 33 includes a constant term, which corresponds to a non-zero, steady-state contact probability. To translate Eq. 33 from a random walk context to a Gaussian polymer context, we relate the time variable, $t$, to the Gaussian polymer segment number, $n$, by recognizing that the mean-square particle displacement maps to polymer end-to-end distance. Thus, we have:

$$\langle(\mathbf{r}(t)-\mathbf{r}(0))^2\rangle = 6Dt \tag{34}$$

$$\langle(\mathbf{r}_n-\mathbf{r}_0)^2\rangle = nl_k^2, \tag{35}$$

where $l_k$ is the Kuhn length of the Gaussian polymer. Thus, $t = nl_k^2/6D$, and Eq. 33 becomes

$$\langle P(R,\alpha,\beta,n|R,\alpha,\beta,0)\rangle = \frac{3}{4\pi a^3}\sum_{l=0}^{\infty}\sum_{p=1}^{\infty}(2l+1)e^{-\frac{x_{l,p}^2 l_k^2}{6a^2}n}, \tag{36}$$

which is the self contact probability for two loci separated by $nl_k$ along a Gaussian polymer with Kuhn length $l_k$, inside a reflecting sphere of radius $a$. Eq. 36 is plotted in Fig. 10 for three values of $\frac{l_k}{a}$.

#### Contact probability for looped chromatin and effective genomic separation

The probability versus genomic separation that two loci on a looped polymer come into contact is modified from that in the absence of loops. Here, we show that for a Gaussian polymer, the contact probability in the presence of loops is readily obtained by replacing the actual genomic separation between the two loci in question by their "effective genomic separation", which is specified below [1].

Our starting point is the probability density for the separation, $\mathbf{r}$, between two points on a Gaussian polymer, separated by genomic distance, $N$, namely

$$P(\mathbf{r}) = \frac{1}{(\frac{2\pi N}{3})^{\frac{3}{2}}}e^{-\frac{3\mathbf{r}^2}{2N}}, \tag{37}$$

where $\mathbf{r}$ is measured in units of the chromatin Kuhn length, $l_k$, which is twice its persistence length. Using Eq. 37, we can immediately write down the probability density for the separation, $\mathbf{r}$, between two points that are both in a Gaussian polymer loop, namely,

$$P(\mathbf{r}) = \frac{1}{(\frac{2\pi}{3}\frac{1}{\frac{1}{N_1}+\frac{1}{N_2}})^{\frac{3}{2}}}e^{\frac{3\mathbf{r}^2}{2}(\frac{1}{N_1}+\frac{1}{N_2})}, \tag{38}$$

where $N_1$ and $N_2$ are the genomic distances around each side of the loop, connecting two points in question.

We compute the probability density for the separation between two points, $\mathbf{r}_1$ and $\mathbf{r}_2$, which are in different loops of a bottlebrush configuration, as shown in Fig. 25. Without loss of generality, we pick $\mathbf{r}_1 = \mathbf{0}$. Then, the probability density of $\mathbf{r}_2$ is

$$P(\mathbf{r}_2) = \int d\mathbf{r}_3\, d\mathbf{r}_4 \frac{1}{(\frac{2\pi}{3})^{\frac{9}{2}}(\frac{1}{\frac{1}{N_1}+\frac{1}{N_2}}N_5\frac{1}{\frac{1}{N_3}+\frac{1}{N_4}})^{\frac{3}{2}}}e^{-\frac{3\mathbf{r}_3^2}{2}(\frac{1}{N_1}+\frac{1}{N_2})}e^{-\frac{3(\mathbf{r}_4-\mathbf{r}_3)^2}{2N_5}}e^{-\frac{3(\mathbf{r}_2-\mathbf{r}_4)^2}{2}(\frac{1}{N_3}+\frac{1}{N_4})}, \tag{39}$$

where $\mathbf{r}_3$ and $\mathbf{r}_4$ are coordinates of the loop bases, and $N_1$, $N_2$, $N_3$, $N_4$, and $N_5$ are the genomic separations of interest, shown in Supplementary Fig. 25. Eq. 39 can be expressed in terms of a covariance matrix, $\Sigma$, as

$$P(\mathbf{r}_2) = \int d\mathbf{r}_3\, d\mathbf{r}_4 \frac{1}{(\frac{2\pi}{3})^{\frac{9}{2}}|\Sigma|^{\frac{3}{2}}} e^{-\frac{1}{2}(x_2,x_3,x_4)\cdot\Sigma^{-1}\cdot(x_2,x_3,x_4)^T}$$
$$e^{-\frac{1}{2}(y_2,y_3,y_4)\cdot\Sigma^{-1}\cdot(y_2,y_3,y_4)^T} e^{-\frac{1}{2}(z_2,z_3,z_4)\cdot\Sigma^{-1}\cdot(z_2,z_3,z_4)^T}, \tag{40}$$

where

$$\Sigma^{-1} = 3\begin{bmatrix} \frac{1}{N_3}+\frac{1}{N_4} & 0 & -\frac{1}{N_3}-\frac{1}{N_4} \\ 0 & \frac{1}{N_1}+\frac{1}{N_2}+\frac{1}{N_5} & -\frac{1}{N_5} \\ -\frac{1}{N_3}-\frac{1}{N_4} & -\frac{1}{N_5} & \frac{1}{N_3}+\frac{1}{N_4}+\frac{1}{N_5} \end{bmatrix}, \tag{41}$$

and

$$|\Sigma| = \frac{1}{3^3}\frac{N_1N_2N_3N_4N_5}{(N_1+N_2)(N_3+N_4)} = \frac{1}{3^3}\frac{1}{\frac{1}{N_1}+\frac{1}{N_2}}N_5\frac{1}{\frac{1}{N_3}+\frac{1}{N_4}} \tag{42}$$

is the determinant of $\Sigma$.

To marginalize $\mathbf{r}_3$ and $\mathbf{r}_4$, it is convenient to express $\Sigma^{-1}$ as

$$\Sigma^{-1} = 3\begin{bmatrix} U & W \\ W^T & V \end{bmatrix} \tag{43}$$

where

$$U = \frac{1}{N_3}+\frac{1}{N_4}, \tag{44}$$

$$V = \begin{bmatrix} \frac{1}{N_1}+\frac{1}{N_2}+\frac{1}{N_5} & -\frac{1}{N_5} \\ -\frac{1}{N_5} & \frac{1}{N_3}+\frac{1}{N_4}+\frac{1}{N_5} \end{bmatrix}, \tag{45}$$

and

$$W = \begin{bmatrix} 0 & -\frac{1}{N_3}-\frac{1}{N_4} \end{bmatrix}. \tag{46}$$

Then, carrying out the integral over $\mathbf{r}_3$ and $\mathbf{r}_4$ in Eq. 40 yields

$$P(\mathbf{r}_2) = \frac{1}{(2\pi)^{\frac{3}{2}}(3^2|\Sigma||V|)^{\frac{3}{2}}} e^{-\frac{1}{2}X\mathbf{r}_2^2} \tag{47}$$

where

$$X = 3(U - W\cdot V^{-1}\cdot W^T) = 3\left(\frac{1}{\frac{1}{\frac{1}{N_1}+\frac{1}{N_2}} + N_5 + \frac{1}{\frac{1}{N_3}+\frac{1}{N_4}}}\right) \tag{48}$$

is a renormalized covariance matrix for $\mathbf{r}_2$ alone, and

$$|V| = \frac{1}{N_1N_3} + \frac{1}{N_1N_4} + \frac{1}{N_1N_5} + \frac{1}{N_2N_3} + \frac{1}{N_2N_4} + \frac{1}{N_2N_5} + \frac{1}{N_3N_5} + \frac{1}{N_4N_5} \tag{49}$$

is the determinant of $V$. It follows that

$$3^2|\Sigma||V| = \frac{1}{3}\frac{(N_1+N_2)N_3N_4 + (N_3+N_4)N_1N_2 + (N_1+N_2)(N_3+N_4)N_5}{(N_1+N_2)(N_3+N_4)} \tag{50}$$

$$= \frac{1}{3}\left(\frac{1}{\frac{1}{N_1}+\frac{1}{N_2}} + N_5 + \frac{1}{\frac{1}{N_3}+\frac{1}{N_4}}\right) \tag{51}$$

$$= \frac{1}{X}. \tag{52}$$

Thus, we have

$$P(\mathbf{r}_2) = \frac{1}{(2\pi)^{\frac{3}{2}}X^{-\frac{3}{2}}}e^{-\frac{1}{2}X\mathbf{r}_2^2}. \tag{53}$$

In analogy with Eq. 37, we define $N_{eff} = \frac{3}{X}$ to be the effective genomic distance between $\mathbf{0}$ and $\mathbf{r}_2$, so that

$$P(\mathbf{r}_2) = \frac{1}{(\frac{2\pi N_{eff}}{3})^{\frac{3}{2}}}e^{-\frac{3\mathbf{r}_2^2}{2N_{eff}}}, \tag{54}$$

which is identical to Eq. 37 in form with $N_{eff}$ replacing $N$.

A useful mnemonics for correctly calculating the effective genomic distance, $N_{eff}$, between any two points on a Gaussian polymer in the bottlebrush configuration shown in Fig. 25 is to combine all polymer segments, $N_1$, $N_2$, $N_3$, $N_4$, $N_5$, *etc.*, in the same way that we combine individual electrical resistances to calculate total resistance. According to this prescription, we immediately have

$$N_{eff} = \frac{1}{\frac{1}{N_1}+\frac{1}{N_2}} + N_5 + \frac{1}{\frac{1}{N_3}+\frac{1}{N_4}}, \tag{55}$$

which is the correct result according to Eq. 48 and $N_{eff} = \frac{3}{X}$. In the case that there are loops between $\mathbf{r}_3$ and $\mathbf{r}_4$, $N_5$ becomes the effective genomic distance between $\mathbf{r}_3$ and $\mathbf{r}_4$.

Our derivation of the effective genomic distance, $N_{eff}$, presumes that the polymer of interest is unconfined, even though we model the chromatin as a polymer inside a spherical shell. However, because the mean genomic separation between chromatin loops is much smaller than the mean genomic separation between chromatin-nuclear envelope contacts, we expect that neglecting chromatin-nuclear envelope contacts will introduce negligible errors into the calculation of effective genomic separation.

### References

[1] Mary Lou P. Bailey, Ivan Surovtsev, Jessica F. Williams, Hao Yan, Tianyu Yuan, Kevin Li, Katherine Duseau, Simon G. J. Mochrie, and Megan C. King. Loops and the activity of loop extrusion factors constrain chromatin dynamics. *Molecular Biology of the Cell*, 34(8):ar78, 2023. PMID: 37126401.

[2] Takeshi Mizuguchi, Geoffrey Fudenberg, Sameet Mehta, Jon-Matthew Belton, Nitika Taneja, Hernan Diego Folco, Peter FitzGerald, Job Dekker, Leonid Mirny, Jemima Barrowman, and Shiv I. S. Grewal. Cohesin-dependent globules and heterochromatin shape 3D genome architecture in S. pombe. *Nature*, 516(7531):432–435, 2014.

[3] Tsung-Han S Hsieh, Geoffrey Fudenberg, Anton Goloborodko, and Oliver J Rando. Micro-c xl: assaying chromosome conformation from the nucleosome to the entire genome. *Nature methods*, 13(12):1009–1011, 2016.

[4] Norihiko Nakazawa, Kenichi Sajiki, Xingya Xu, Alejandro Villar-Briones, Orie Arakawa, and Mitsuhiro Yanagida. Rna pol ii transcript abundance controls condensin accumulation at mitotically up-regulated and heat-shock-inducible genes in fission yeast. *Genes to Cells*, 20(6):481–499, 2015.

[5] Ralf Bundschuh and Terence Hwa. Statistical mechanics of secondary structures formed by random RNA sequences. *Physical Review E*, 65(3):031903, 2002.

[6] Stephanie A Schalbetter, Geoffrey Fudenberg, Jonathan Baxter, Katherine S Pollard, and Matthew J Neale. Principles of meiotic chromosome assembly revealed in S. cerevisiae. *Nature Communications*, 10(1):4795, 2019.

[7] Christine K Schmidt, Neil Brookes, and Frank Uhlmann. Conserved features of cohesin binding along fission yeast chromosomes. *Genome Biology*, 10(5):1–16, 2009.

[8] Patricia Garcia, Rita Fernandez-Hernandez, Ana Cuadrado, Ignacio Coca, Antonio Gomez, Maria Maqueda, Ana Latorre-Pellicer, Beatriz Puisac, Feliciano J Ramos, Juan Sandoval, Manel Esteller, Jose Luis Mosquera, Jairo Rodriguez, J Pié, Ana Losada, and Ethel Queralt. Disruption of nipbl/scc2 in cornelia de lange syndrome provokes cohesin genome-wide redistribution with an impact in the transcriptome. *Nature Communications*, 12(1):4551, 2021.

[9] Lorenzo Costantino, Tsung-Han S Hsieh, Rebecca Lamothe, Xavier Darzacq, and Douglas Koshland. Cohesin residency determines chromatin loop patterns. *Elife*, 9:e59889, 2020.

[10] Christophe Chapard, Robert Jones, Till van Oepen, Johanna C Scheinost, and Kim Nasmyth. Sister dna entrapment between juxtaposed smc heads and kleisin of the cohesin complex. *Molecular cell*, 75(2):224–237, 2019.

[11] Mark Mattingly, Chris Seidel, Sofía Muñoz, Yan Hao, Ying Zhang, Zhihui Wen, Laurence Florens, Frank Uhlmann, and Jennifer L Gerton. Mediator recruits the cohesin loader scc2 to rna pol ii-transcribed genes and promotes sister chromatid cohesion. *Current Biology*, 32(13):2884–2896, 2022.

[12] Tianyu Yuan, Hao Yan, Mary Lou P Bailey, Jessica F Williams, Ivan Surovtsev, Megan C King, and Simon GJ Mochrie. Effect of loops on the mean-square displacement of rouse-model chromatin. *Physical Review E*, 109(4):044502, 2024.

| | Chr1: 0.6–1.6 Mb | Chr1: 4.3–5.3 Mb | Chr2: 0.3–1.3 Mb | Chr2: 1.9–2.9 Mb | Chr3: 1.3–2.3 Mb |
|---|---|---|---|---|---|
| Chr1: 0.6–1.6 Mb | 1 (1) | - | - | - | - |
| Chr1: 4.3–5.3 Mb | 1.1753 (0.1591) | 1 (1) | - | - | - |
| Chr2: 0.3–1.3 Mb | 1.1830 (0.0237) | 1.1850 (0.0456) | 1 (1) | - | - |
| Chr2: 1.9–2.9 Mb | 1.1934 (-0.0293) | 1.1868 (0.0362) | 1.1965 (-0.0443) | 1 (1) | - |
| Chr3: 1.3–2.3 Mb | 1.2002 (-0.1818) | 1.1511 (0.3174) | 1.1901 (-0.0784) | 1.1766 (0.0964) | 1 (1) |

Table 1: Mean pairwise ratio (MPR) score and $P(s)$-scaled Pearson correlation coefficient (PCC) between experimental Hi-C maps from different genomic regions in *S. pombe*. $P(s)$-scaled Pearson correlation coefficients (PCCs) are listed in parentheses. To calculate the $P(s)$-scaled PCC between two Hi-C maps, each diagonal of the Hi-C maps is scaled relative to the mean of all entries of that diagonal before the PCC calculation is performed.

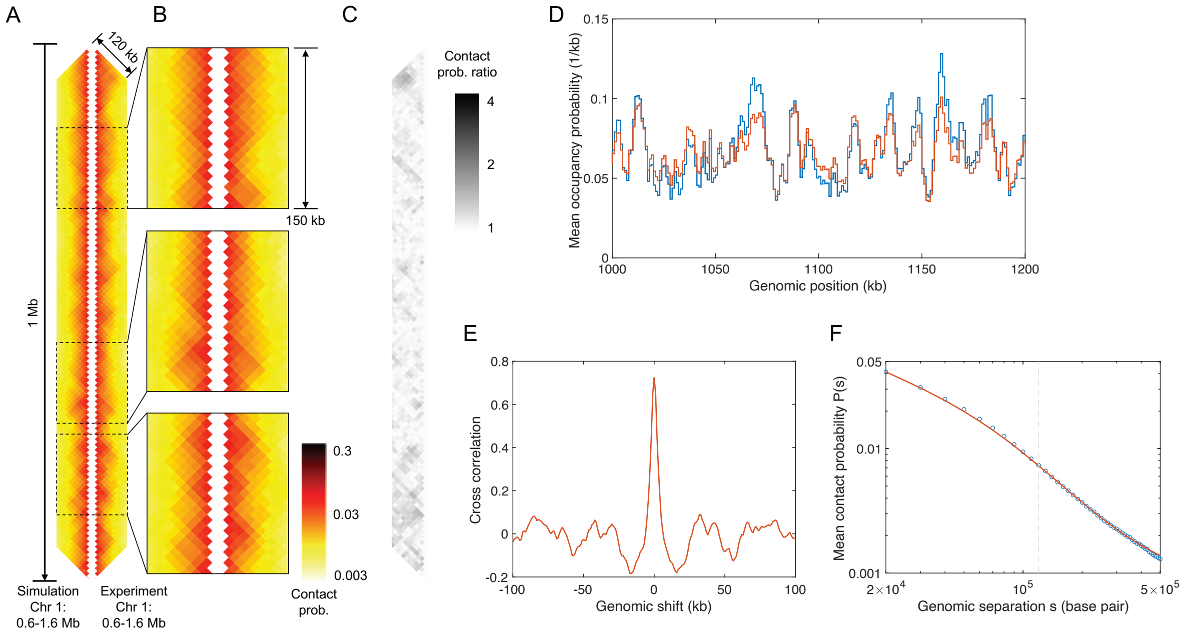


Supplementary Figure 1: Conserved-current loop extrusion (CCLE) model recapitulates TAD-scale chromatin organization in the 0.6–1.6 Mb region of Chr 1 of interphase *S. pombe.* (A) Comparison between Hi-C map of 1 Mb region generated by the CCLE model (using interphase Psc3 ChIP-seq data [2]) and the experimental Hi-C map [2] of the same region. Both Hi-C maps show interactions up to genomic separation of 120 kb. (B) Magnified comparison of experimental and simulated Hi-C for three representative sub-regions, each 150 kb in size, located at 0.76–0.91 Mb, 1.16–1.31 Mb, and 1.33–1.48 Mb, from top to bottom. (C) Contact probability ratio map between the Hi-C maps in panel (A). (D) Normalized experimental cohesin (Psc3) occupancy (blue) and simulated LEF occupancy probability (red) of the 1.0–1.2 Mb region of Chr 1. (E) Cross-correlation between the experimental interphase cohesin (Psc3) ChIP-seq and simulated LEF occupancy probability. (F) Chromatin contact probability, $P(s)$, as a function of genomic distance, $s$, for the experimental (blue) and simulated (red) Hi-C. Results shown use the best-fit parameters given in the main text. The optimization process is discussed in Supplementary Methods.

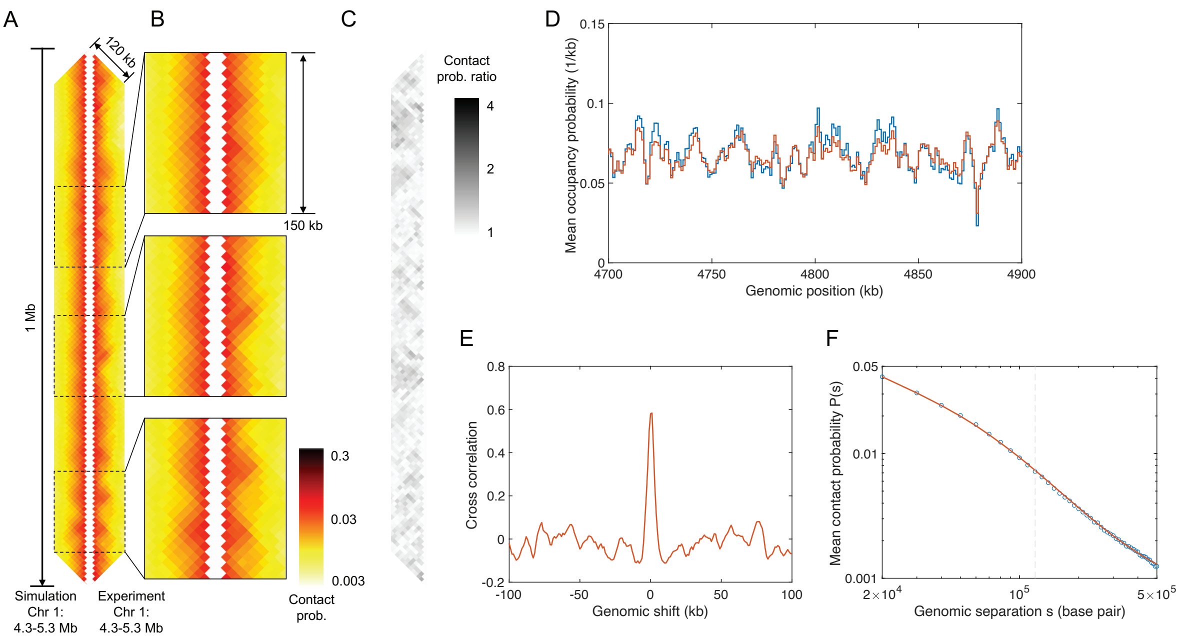


Supplementary Figure 2: Conserved-current loop extrusion (CCLE) model recapitulates TAD-scale chromatin organization in the 4.3–5.3 Mb region of Chr 1 of interphase *S. pombe*. (A) Comparison between Hi-C map of 1 Mb region generated by the CCLE model (using interphase Psc3 ChIP-seq data [2]) and the experimental Hi-C map [2] of the same region. Both Hi-C maps show interactions up to genomic separation of 120 kb. (B) Magnified comparison of experimental and simulated Hi-C for three representative sub-regions, each 150 kb in size, located at 4.56–4.71 Mb, 4.80–4.95 Mb, and 5.09–5.24 Mb, from top to bottom. (C) Contact probability ratio map between the Hi-C maps in panel (A). (D) Normalized experimental cohesin (Psc3) occupancy (blue) and simulated LEF occupancy probability (red) of the 4.7–4.9 Mb region of Chr 1. (E) Cross-correlation between the experimental interphase cohesin (Psc3) ChIP-seq and simulated LEF occupancy probability. (F) Chromatin contact probability, $P(s)$, as a function of genomic distance, $s$, for the experimental (blue) and simulated (red) Hi-C. Results shown use the best-fit parameters given in the main text. The optimization process is discussed in Supplementary Methods.

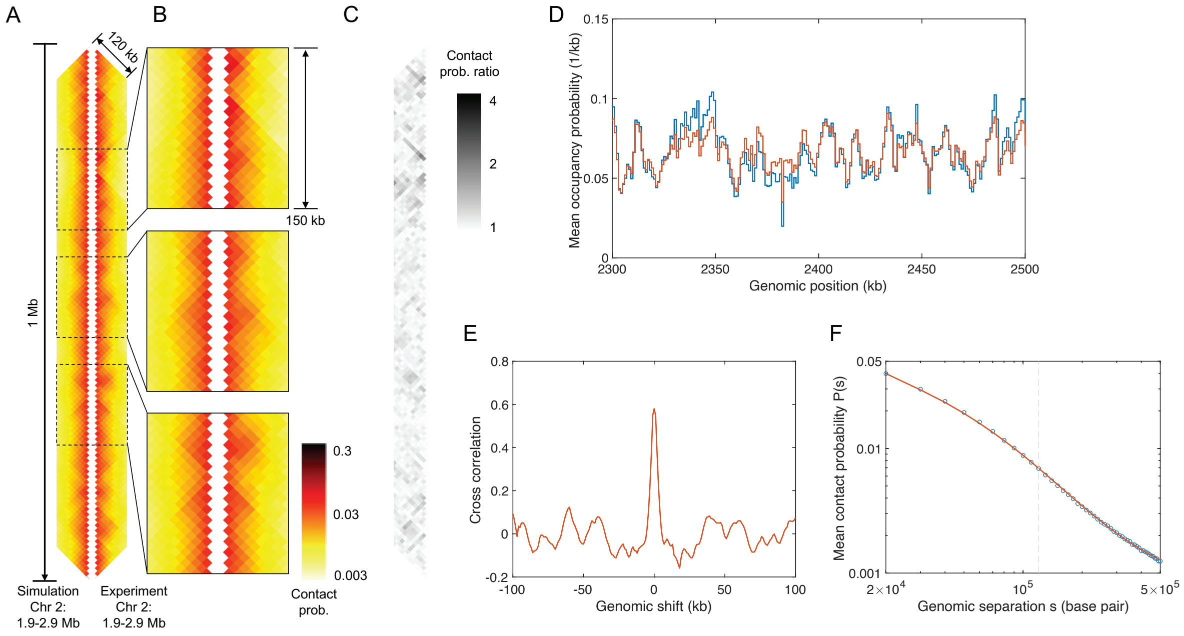


Supplementary Figure 3: Conserved-current loop extrusion (CCLE) model recapitulates TAD-scale chromatin organization in the 1.9–2.9 Mb region of Chr 2 of interphase *S. pombe*. (A) Comparison between Hi-C map of 1 Mb region generated by the CCLE model (using interphase Psc3 ChIP-seq data [2]) and the experimental Hi-C map [2] of the same region. Both Hi-C maps show interactions up to genomic separation of 120 kb. (B) Magnified comparison of experimental and simulated Hi-C for three representative sub-regions, each 150 kb in size, located at 2.10–2.25 Mb, 2.30–2.45 Mb, and 2.50–2.65 Mb, from top to bottom. (C) Contact probability ratio map between the Hi-C maps in panel (A). (D) Normalized experimental cohesin (Psc3) occupancy (blue) and simulated LEF occupancy probability (red) of the 2.3–2.5 Mb region of Chr 2. (E) Cross-correlation between the experimental interphase cohesin (Psc3) ChIP-seq and simulated LEF occupancy probability. (F) Chromatin contact probability, $P(s)$, as a function of genomic distance, $s$, for the experimental (blue) and simulated (red) Hi-C. Results shown use the best-fit parameters given in the main text. The optimization process is discussed in Supplementary Methods.

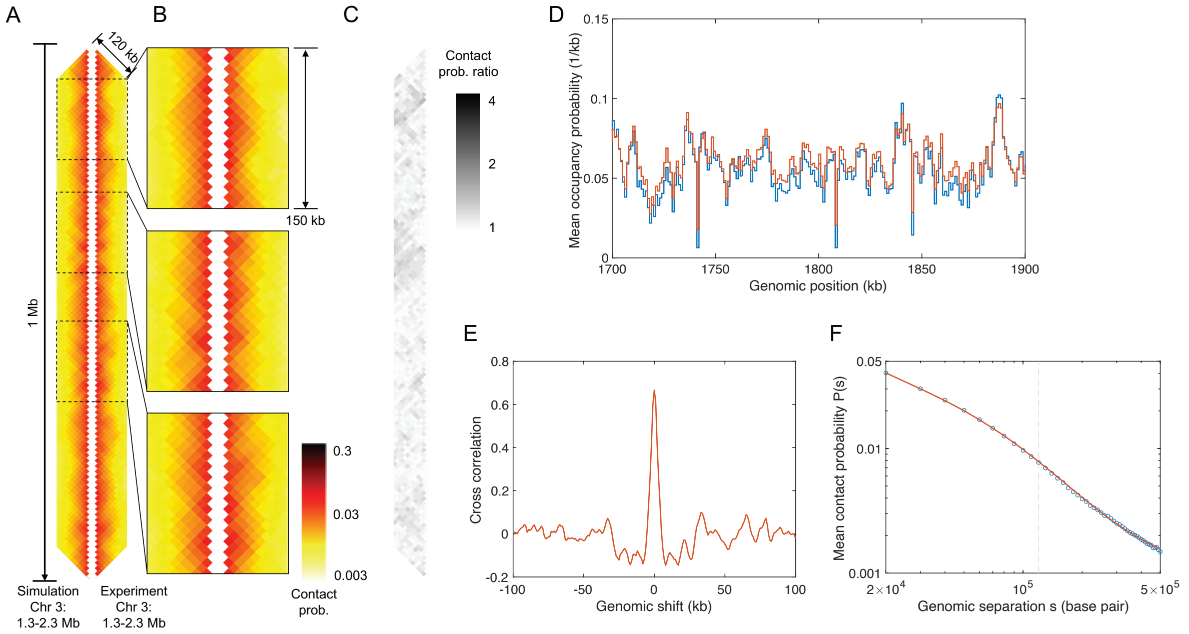


Supplementary Figure 4: Conserved-current loop extrusion (CCLE) model recapitulates TAD-scale chromatin organization in the 1.3–2.3 Mb region of Chr 3 of interphase *S. pombe*. (A) Comparison between Hi-C map of 1 Mb region generated by the CCLE model (using interphase Psc3 ChIP-seq data [2]) and the experimental Hi-C map [2] of the same region. Both Hi-C maps show interactions up to genomic separation of 120 kb. (B) Magnified comparison of experimental and simulated Hi-C for three representative sub-regions, each 150 kb in size, located at 1.37–1.52 Mb, 1.58–1.73 Mb, and 1.82–1.97 Mb, from top to bottom. (C) Contact probability ratio map between the Hi-C maps in panel (A). (D) Normalized experimental cohesin (Psc3) occupancy (blue) and simulated LEF occupancy probability (red) of the 1.7–1.9 Mb region of Chr 3. (E) Cross-correlation between the experimental interphase cohesin (Psc3) ChIP-seq and simulated LEF occupancy probability. (F) Chromatin contact probability, $P(s)$, as a function of genomic distance, $s$, for the experimental (blue) and simulated (red) Hi-C. Results shown use the best-fit parameters given in the main text. The optimization process is discussed in Supplementary Methods.

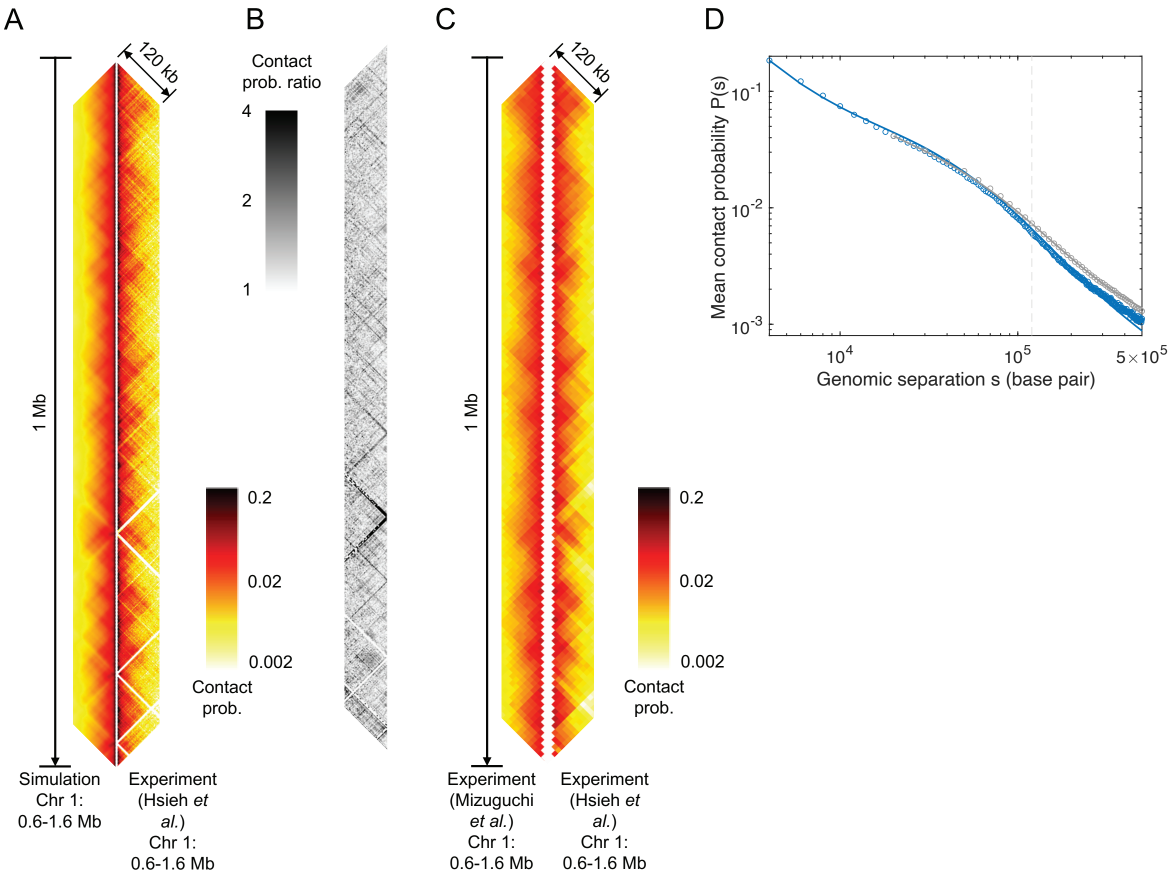


Supplementary Figure 5: Conserved-current loop extrusion (CCLE) model recapitulates TAD-scale chromatin organization in the region Chr 1: 0.6–1.6 Mb of interphase *S. pombe.* (A) Comparison between Hi-C map of 1 Mb region generated by the CCLE model (using interphase Psc3 ChIP-seq data [2]) and the experimental Hi-C map [3] of the same region, binned to 2 kb resolution. Both Hi-C maps show interactions up to genomic separation of 120 kb. (B) Contact probability ratio map between the Hi-C maps in panel (A). (C) Comparison between two Hi-C maps of the same regions from two different sources [2, 3], binned to 10 kb resolution. (D) Chromatin contact probability, $P(s)$, as a function of genomic separation, $s$, for the experimental (circle) and simulated (solid line) Hi-C. P(s) curve of Hi-C data from Hsieh *et al.* and that of its corresponding simulation are shown in blue; P(s) curve of Hi-C data from Mizuguchi *et al.* and that of its corresponding simulation are shown in gray. Parameters for the simulation results corresponding to the Hi-C data of Hsieh *et al.* are the same as those presented in Table 2 in the main text except for the LEF density, which is 0.037 $kb^{-1}$, and the persistence length, which is 30 nm. The MPR and PCC scores are 1.6005 and 0.4277, respectively, for the comparison in panel (A), and 1.2248 and 0.7665, respectively, for the comparison in panel (C).

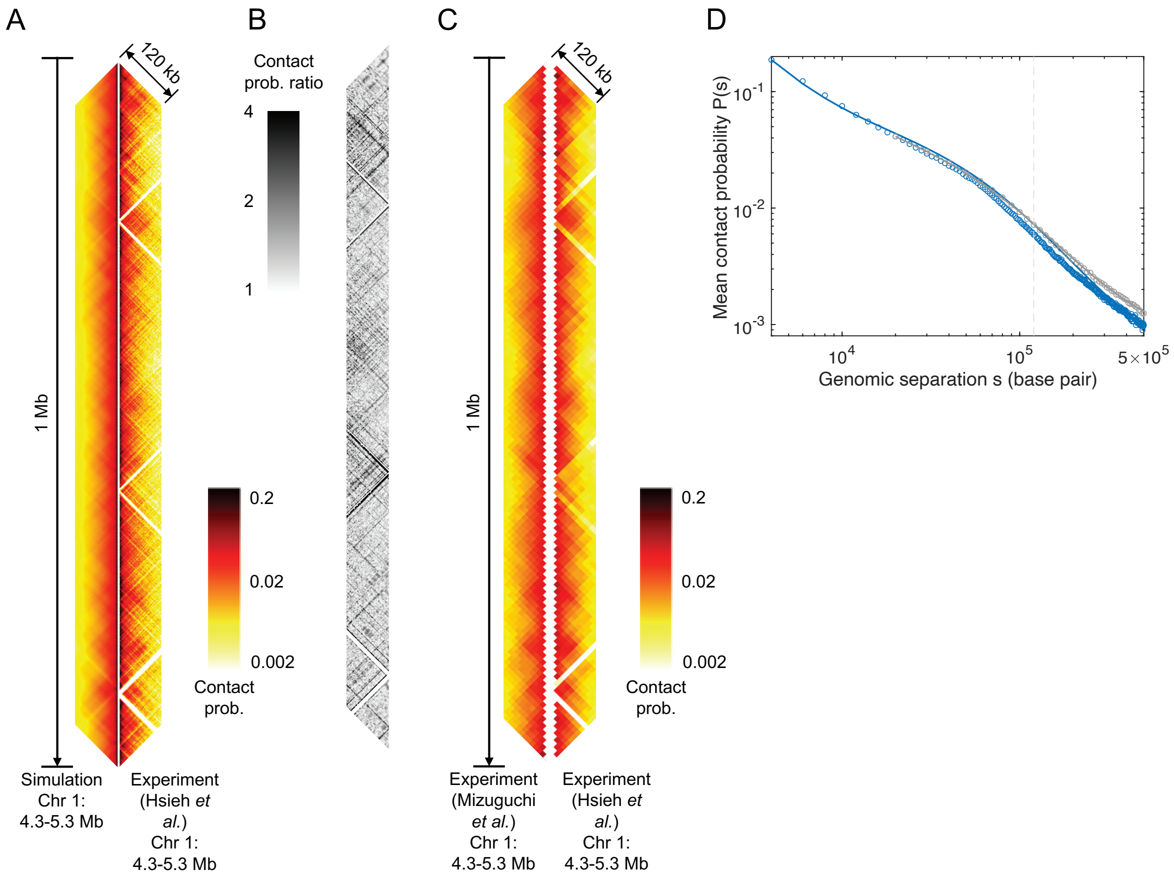


Supplementary Figure 6: Conserved-current loop extrusion (CCLE) model recapitulates TAD-scale chromatin organization in the region Chr 1: 4.3–5.3 Mb of interphase *S. pombe.* (A) Comparison between Hi-C map of 1 Mb region generated by the CCLE model (using interphase Psc3 ChIP-seq data [2]) and the experimental Hi-C map [3] of the same region, binned to 2 kb resolution. Both Hi-C maps show interactions up to genomic separation of 120 kb. (B) Contact probability ratio map between the Hi-C maps in panel (A). (C) Comparison between two Hi-C maps of the same regions from two different sources [2, 3], binned to 10 kb resolution. (D) Chromatin contact probability, $P(s)$, as a function of genomic distance, $s$, for the experimental (circle) and simulated (solid line) Hi-C. P(s) curve of Hi-C data from Hsieh *et al.* and that of its corresponding simulation are shown in blue; P(s) curve of Hi-C data from Mizuguchi *et al.* and that of its corresponding simulation are shown in gray. Parameters for the simulation results corresponding to the Hi-C data of Hsieh *et al.* are the same as those presented in Table 2 in the main text except for the persistence length, which is 30 nm. The MPR and PCC scores are 1.5620 and 0.2629, respectively, for the comparison in panel (A), and 1.6327 and 0.6187, respectively, for the comparison in panel (C).

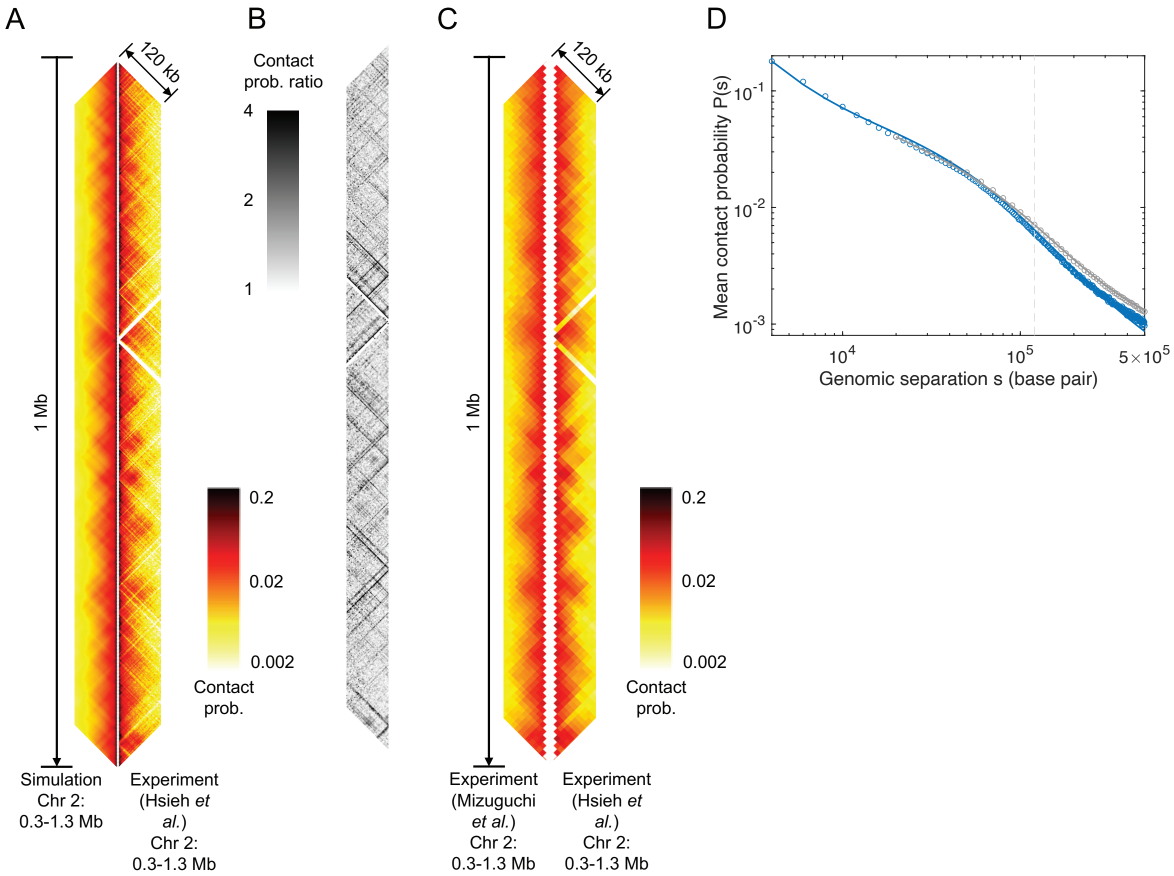


Supplementary Figure 7: Conserved-current loop extrusion (CCLE) model recapitulates TAD-scale chromatin organization in the region Chr 2: 0.3–1.3 Mb of interphase *S. pombe.* (A) Comparison between Hi-C map of 1 Mb region generated by the CCLE model (using interphase Psc3 ChIP-seq data [2]) and the experimental Hi-C map [3] of the same region, binned to 2 kb resolution. Both Hi-C maps show interactions up to genomic separation of 120 kb. (B) Contact probability ratio map between the Hi-C maps in panel (A). (C) Comparison between two Hi-C maps of the same regions from two different sources [2, 3], binned to 10 kb resolution. (D) Chromatin contact probability, $P(s)$, as a function of genomic distance, $s$, for the experimental (circle) and simulated (solid line) Hi-C. P(s) curve of Hi-C data from Hsieh *et al.* and that of its corresponding simulation are shown in blue; P(s) curve of Hi-C data from Mizuguchi *et al.* and that of its corresponding simulation are shown in gray. Parameters for the simulation results corresponding to the Hi-C data of Hsieh *et al.* are the same as those presented in Table 2 in the main text except for the LEF density, which is 0.037 $kb^{-1}$, and the persistence length, which is 30 nm. The MPR and PCC scores are 1.4943 and 0.4648, respectively, for the comparison in panel (A), and 1.2579 and 0.7668, respectively, for the comparison in panel (C).

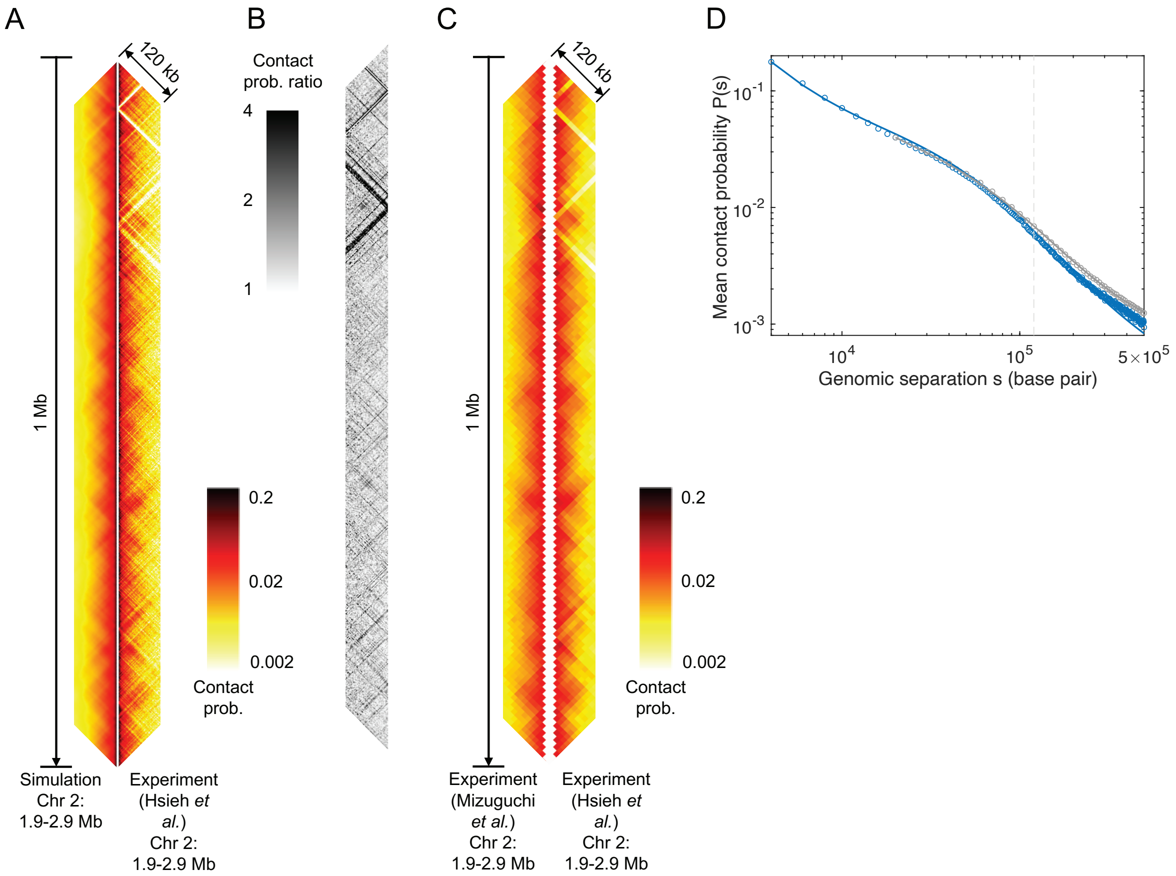


Supplementary Figure 8: Conserved-current loop extrusion (CCLE) model recapitulates TAD-scale chromatin organization in the region Chr 2: 1.9–2.9 Mb of interphase *S. pombe*. (A) Comparison between Hi-C map of 1 Mb region generated by the CCLE model (using interphase Psc3 ChIP-seq data [2]) and the experimental Hi-C map [3] of the same region, binned to 2 kb resolution. Both Hi-C maps show interactions up to genomic separation of 120 kb. (B) Contact probability ratio map between the Hi-C maps in panel (A). (C) Comparison between two Hi-C maps of the same regions from two different sources [2, 3], binned to 10 kb resolution. (D) Chromatin contact probability, $P(s)$, as a function of genomic distance, $s$, for the experimental (circle) and simulated (solid line) Hi-C. P(s) curve of Hi-C data from Hsieh *et al.* and that of its corresponding simulation are shown in blue; P(s) curve of Hi-C data from Mizuguchi *et al.* and that of its corresponding simulation are shown in gray. Parameters for the simulation results corresponding to the Hi-C data of Hsieh *et al.* are the same as those presented in Table 2 in the main text except for the LEF density, which is 0.037 $kb^{-1}$, and the persistence length, which is 30 nm. The MPR and PCC scores are 1.5107 and 0.3640, respectively, for the comparison in panel (A), and 1.2367 and 0.5398, respectively, for the comparison in panel (C).

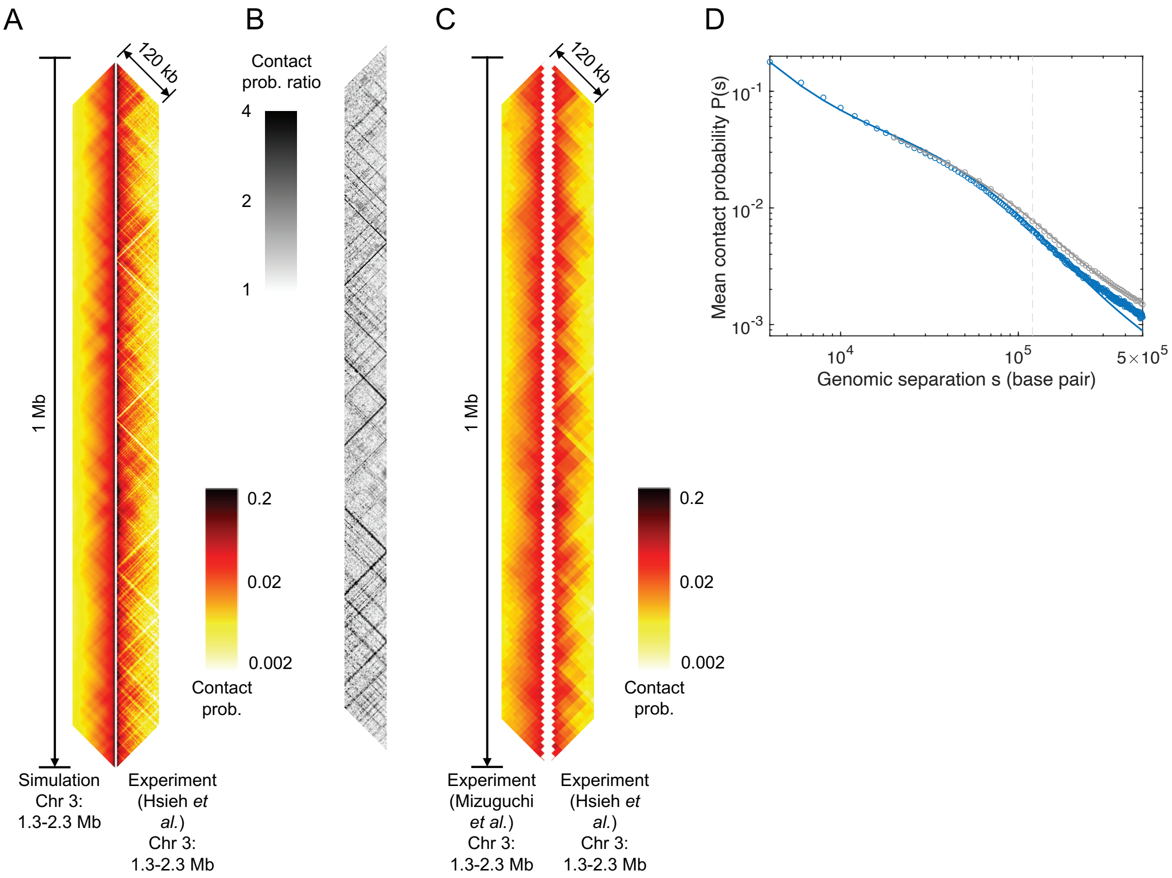


Supplementary Figure 9: Conserved-current loop extrusion (CCLE) model recapitulates TAD-scale chromatin organization in the region Chr 3: 1.3–2.3 Mb of interphase *S. pombe*. (A) Comparison between Hi-C map of 1 Mb region generated by the CCLE model (using interphase Psc3 ChIP-seq data [2]) and the experimental Hi-C map [3] of the same region, binned to 2 kb resolution. Both Hi-C maps show interactions up to genomic separation of 120 kb. (B) Contact probability ratio map between the Hi-C maps in panel (A). (C) Comparison between two Hi-C maps of the same regions from two different sources [2, 3], binned to 10 kb resolution. (D) Chromatin contact probability, $P(s)$, as a function of genomic distance, $s$, for the experimental (circle) and simulated (solid line) Hi-C. P(s) curve of Hi-C data from Hsieh *et al.* and that of its corresponding simulation are shown in blue; P(s) curve of Hi-C data from Mizuguchi *et al.* and that of its corresponding simulation are shown in gray. Parameters for the simulation results corresponding to the Hi-C data of Hsieh *et al.* are the same as those presented in Table 2 in the main text except for the LEF density, which is 0.033 $kb^{-1}$, and the persistence length, which is 30 nm. The MPR and PCC scores are 1.4922 and 0.4232, respectively, for the comparison in panel (A), and 1.2196 and 0.6277, respectively, for the comparison in panel (C).

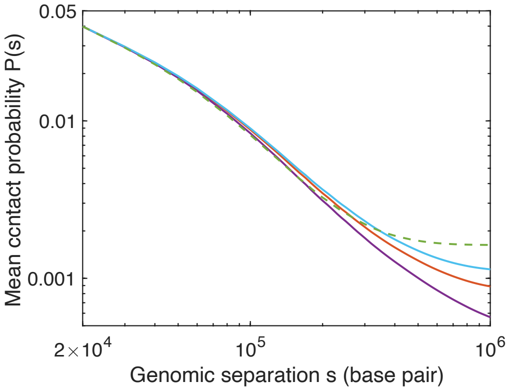


Supplementary Figure 10: Chromatin contact probability is sensitive to persistence lengths and boundary conditions only for large genomic separations. Chromatin mean contact probability, $P(s)$, as a function of genomic separation, $s$, of the CCLE-simulated Hi-C map of Chr 2: 1.8 Mb–3.0 Mb in interphase *S. pombe*, for different persistence lengths of 50 nm (purple), 80 nm (red), and 100 nm (cyan), using reflecting boundary condition. The green dashed line shows the $P(s)$ curve of the simulated Hi-C map of the same region with persistence length of 80 nm and absorbing boundary condition (calculation not included in the text).

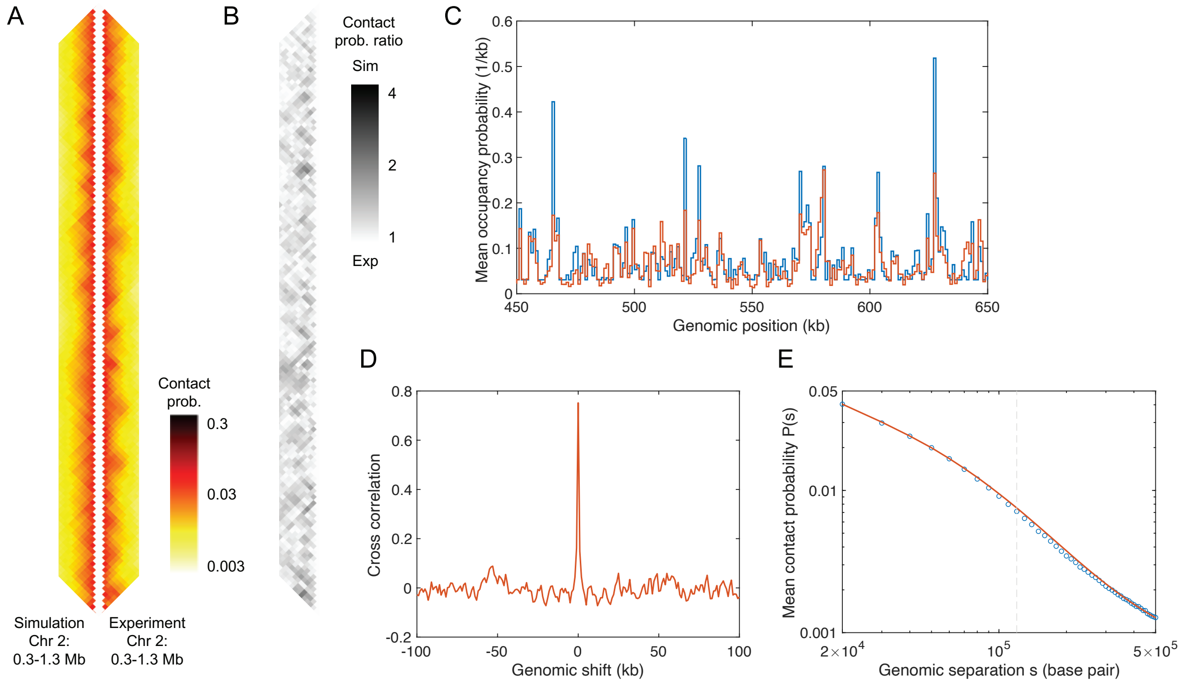


Supplementary Figure 11: Condensin (Cut14) ChIP-seq data generates simulated Hi-C map that poorly agrees with experimental Hi-C map in interphase *S. pombe*. (A) Comparison between Hi-C map of 1 Mb region generated by the CCLE model (using Cut14 ChIP-seq data of interphase *S. pombe* [4]) and the experimental Hi-C map [2] of the same region. Both Hi-C maps show interactions up to genomic separation of 120 kb. The MPR and PCC scores, using best-fit parameters, are 1.1793 and -0.0837, respectively. (B) Contact probability ratio map between the Hi-C shown in panel (A). (C) Normalized experimental condensin (Cut14) occupancy (blue) and simulated LEF occupancy probability (red) of 0.45–0.65 Mb region of Chr 2. (D) Cross-correlation between the experimental interphase condensin (Cut14) ChIP-seq and simulated LEF occupancy probability. (E) Chromatin contact probability, $P(s)$, as a function of genomic distance, $s$, for the experimental (blue) and simulated (red) Hi-C.

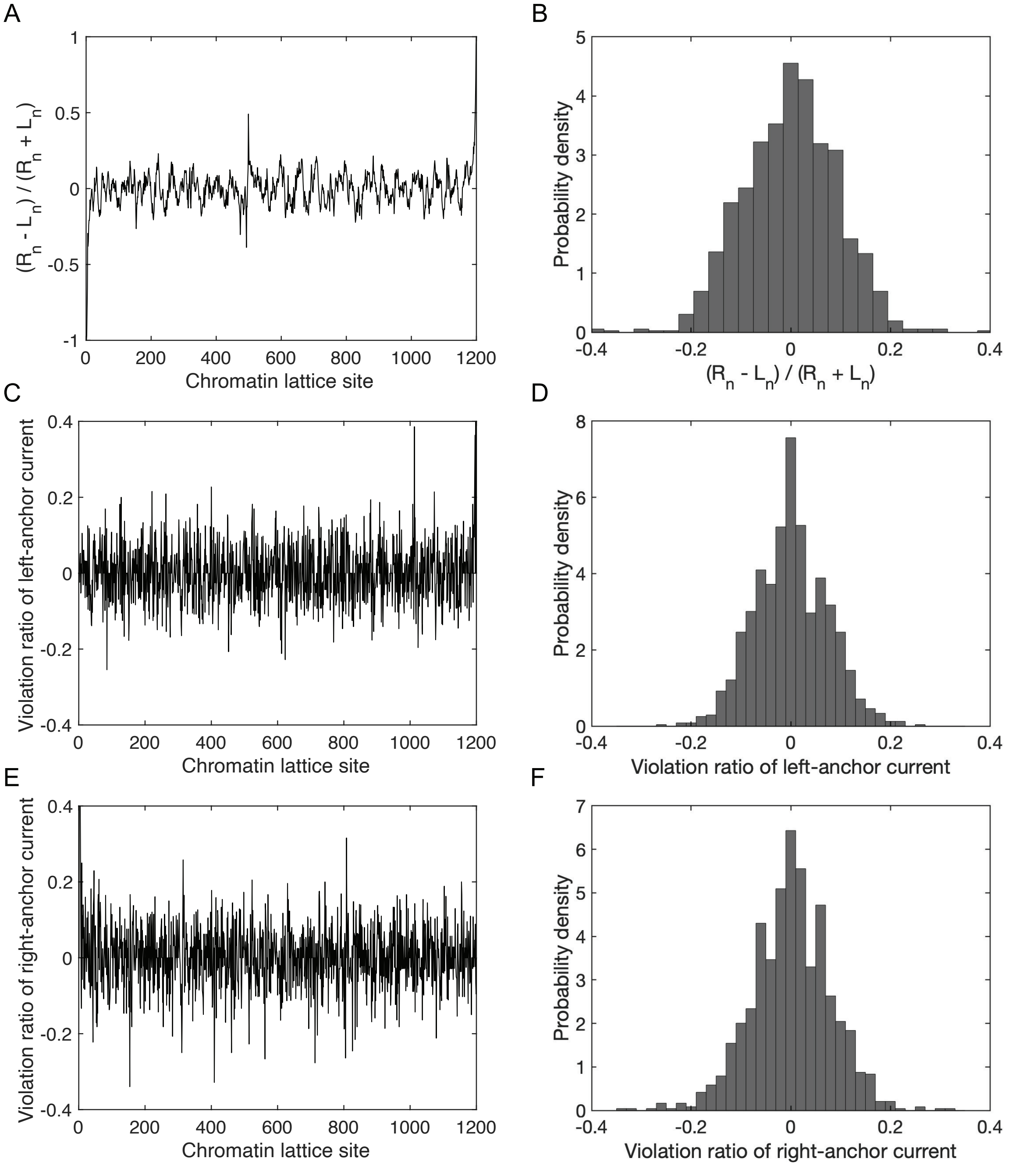


Supplementary Figure 12: Assumptions of CCLE are satisfied in simulations for interphase *S. pombe.* (A) Fractional imbalance of right- and left-moving LEF anchors, *i.e.* $(R_n - L_n)/(R_n + L_n)$, at each lattice site $n$. (C) Ratio of net current of left-moving LEF anchors at each site, that violates current conservation, as a result of binding and unbinding, to the mean current of left-moving LEF anchors, that satisfies current conservation. (E) Ratio of net current of right-moving LEF anchors at each site, that violates current conservation, as a result of binding and unbinding, to the mean current of right-moving LEF anchors, that satisfies current conservation. (B), (D), and (F) Corresponding distributions of values in panels (A), (C), and (E), respectively. Means of distributions in panels (B), (D), and (F) are 0.000005, 0.0035, and 0.0038, respectively; standard deviations are 0.1180, 0.0967, and 0.1023, respectively.

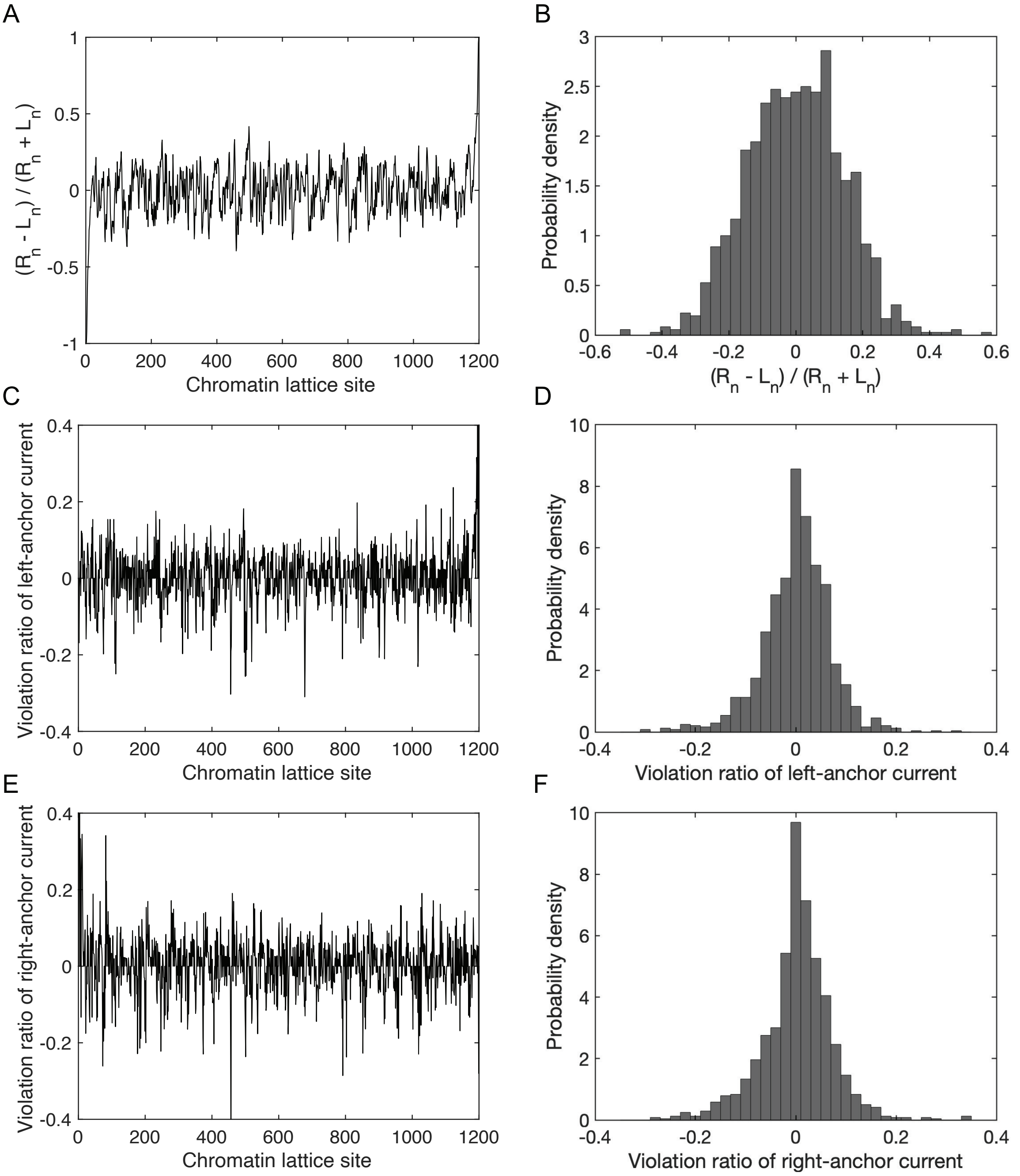


Supplementary Figure 13: Assumptions of CCLE are satisfied in simulations for meiotic *S. cerevisiae*. (A) Fractional imbalance of right- and left-moving LEF anchors, *i.e.* $(R_n - L_n)/(R_n + L_n)$, at each lattice site $n$. (C) Ratio of net current of left-moving LEF anchors at each site, that violates current conservation, as a result of binding and unbinding, to the mean current of left-moving LEF anchors, that satisfies current conservation. (E) Ratio of net current of right-moving LEF anchors at each site, that violates current conservation, as a result of binding and unbinding, to the mean current of right-moving LEF anchors, that satisfies current conservation. (B), (D), and (F) Corresponding distributions of values in panels (A), (C), and (E), respectively. Means of distributions in panels (B), (D), and (F) are -0.0014, 0.0046, and 0.0047, respectively; standard deviations are 0.1645, 0.0945, and 0.0991, respectively.

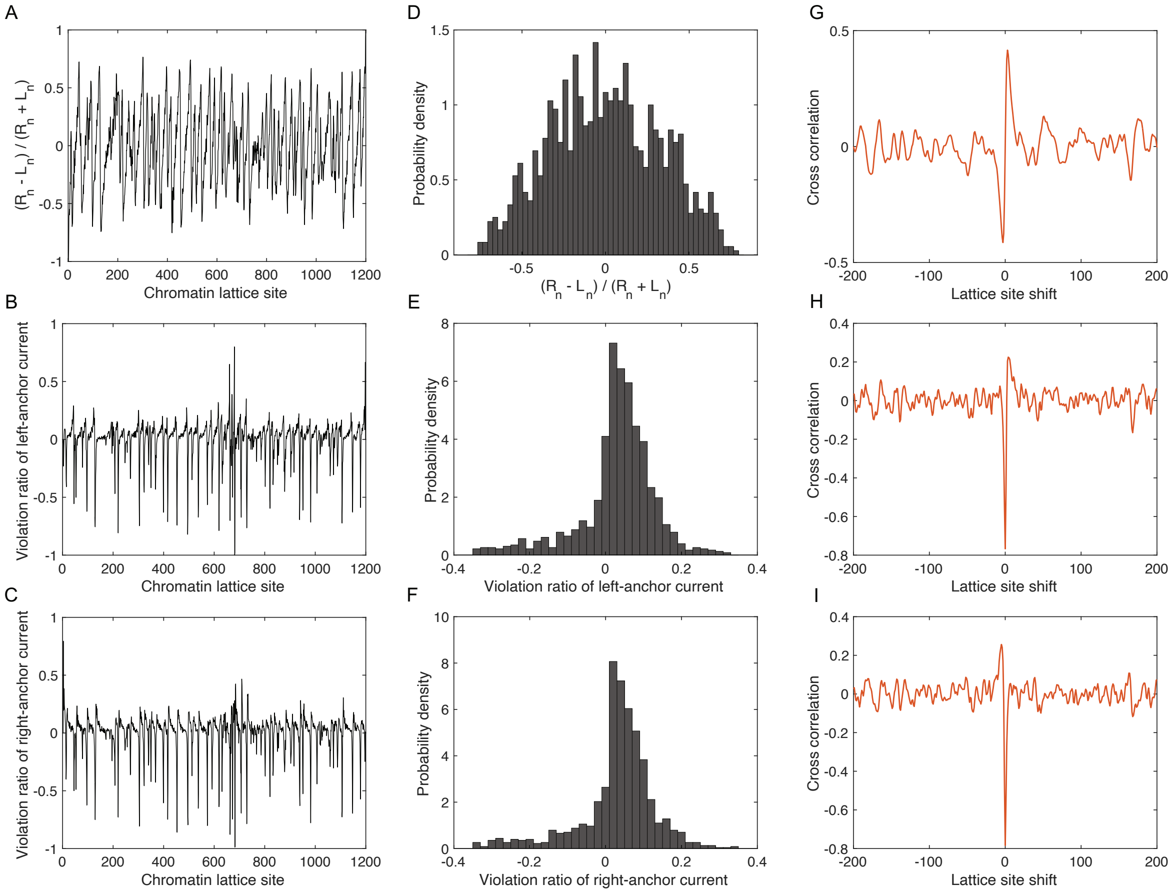


Supplementary Figure 14: Assumptions of CCLE are satisfied in simulations of mitotic *S. cerevisiae* away from cohesin peaks. (A) Fractional imbalance of right- and left-moving LEF anchors, *i.e.* $(R_n - L_n)/(R_n + L_n)$, at each lattice site $n$. (B) Ratio of net current of left-moving LEF anchors at each site, that violates current conservation, as a result of binding and unbinding, to the mean current of left-moving LEF anchors, that satisfies current conservation. (C) Ratio of net current of right-moving LEF anchors at each site, that violates current conservation, as a result of binding and unbinding, to the mean current of right-moving LEF anchors, that satisfies current conservation. (D), (E), and (F) Corresponding distributions of values in panels (A), (B), and (C), respectively. Means of distributions in panels (D), (E), and (F) are -0.0023, 0.0030, and 0.0033, respectively; standard deviations are 0.3332, 0.1710, and 0.1687, respectively. (G), (H), and (I) Cross-correlations between the curve shown in each of panel (A), (B), and (C), respectively, and the mitotic cohesin ChIP-seq data.

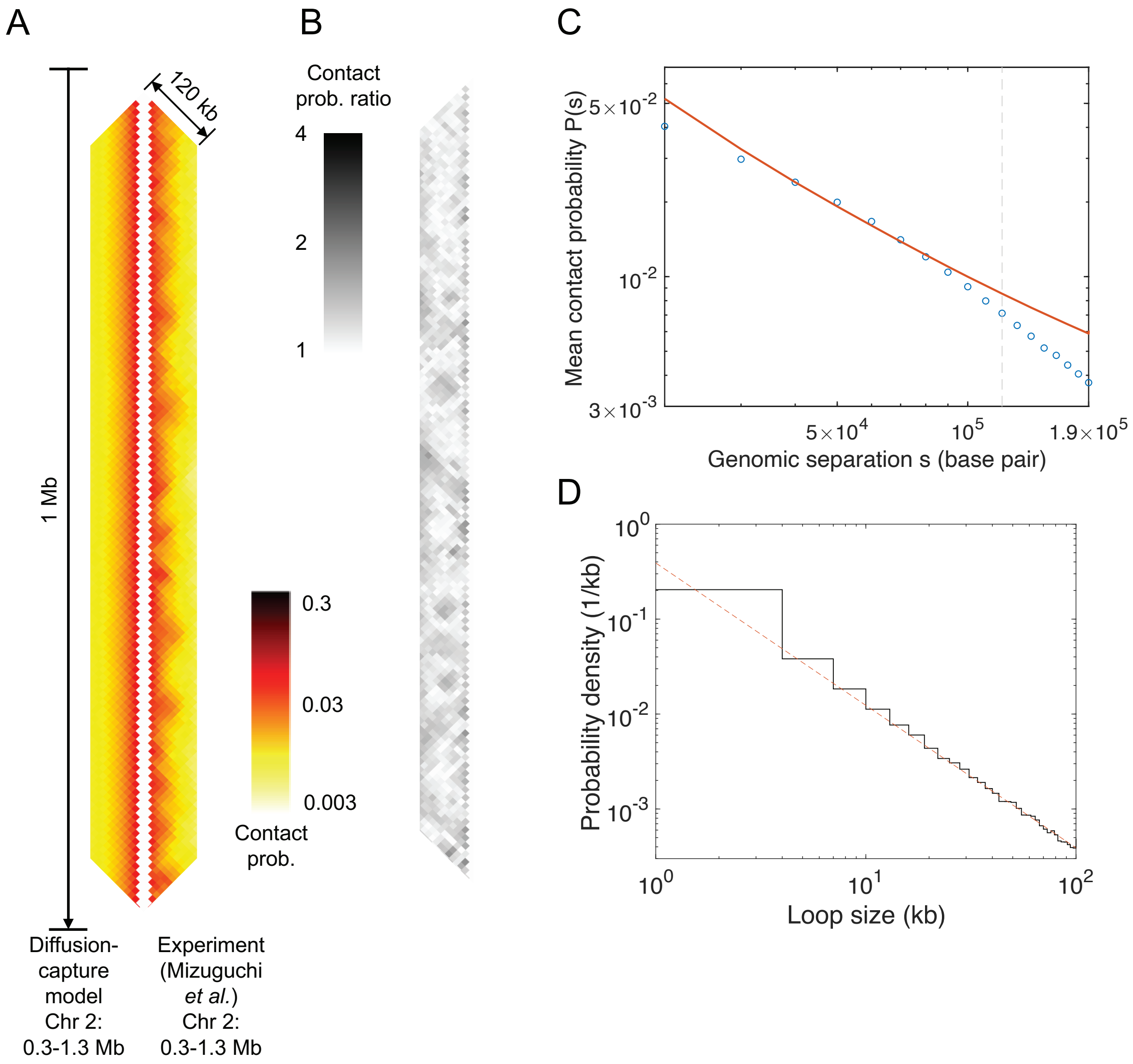


Supplementary Figure 15: Diffusion capture model poorly describes TAD-scale chromatin organization in the region Chr 2: 0.3-1.3 Mb of interphase *S. pombe*. (A) Comparison between Hi-C map of 1 Mb region generated by the diffusion capture model (using interphase Psc3 ChIP-seq data [2]) and the experimental Hi-C map [2] of the same region, binned to 10 kb resolution. Both Hi-C maps show interactions up to genomic separation of 120 kb. Diagonals of the simulation contact map are overall scaled such that the mean of the fourth diagonal (corresponding to genomic separation of 40 kb) equals to that of the experimental Hi-C map. (B) Contact probability ratio map between the Hi-C maps in panel (A). (C) Chromatin contact probability, $P(s)$, as a function of genomic separation, $s$, for the experimental (blue circle) and simulated (red line) Hi-C. Vertical dashed line indicates the maximum genomic separation illustrated in the Hi-C and ratio maps in panels (A) and (B). (D) Loop size distribution in the diffusion capture model (solid line). The red, dashed line indicates the $-\frac{3}{2}$-power scaling. The only two parameters in the diffusion capture model, namely the LEF density and persistence length, are set to be 0.033 $kb^{-1}$ and 80 nm, respectively, for the simulation results shown. The MPR and PCC scores are 1.1754 and 0.2675, respectively, for the comparison in panel (A).

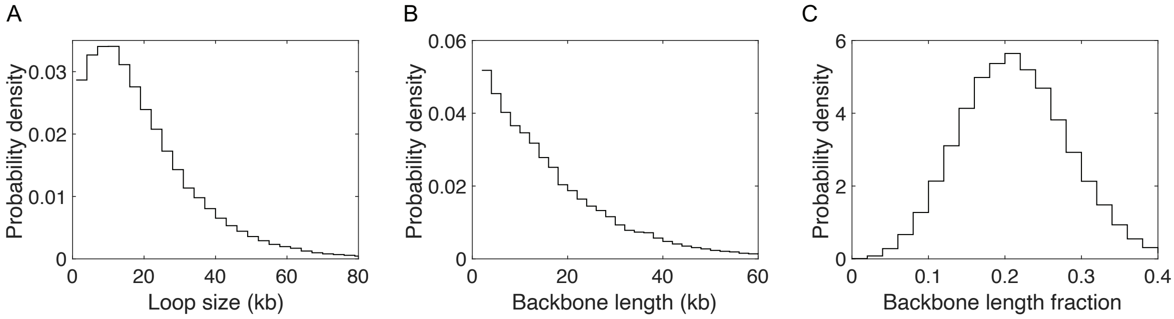


Supplementary Figure 16: Loop properties in meiotic *S. cerevisiae.* (A), (B), and (C) Distributions of loop size, backbone segment length, and chromatin compaction ratio (as measured by the fraction of the chromatin contour length within the backbone), respectively, for the region Chr 13: 240–840 kb of meiotic *S. cerevisiae.* Means of loop size, backbone segment length, and chromatin compaction ratio are 19.82 kb, 16.06 kb, and 0.2129, respectively; standard deviations are 15.97 kb, 14.67 kb, and 0.0702, respectively.

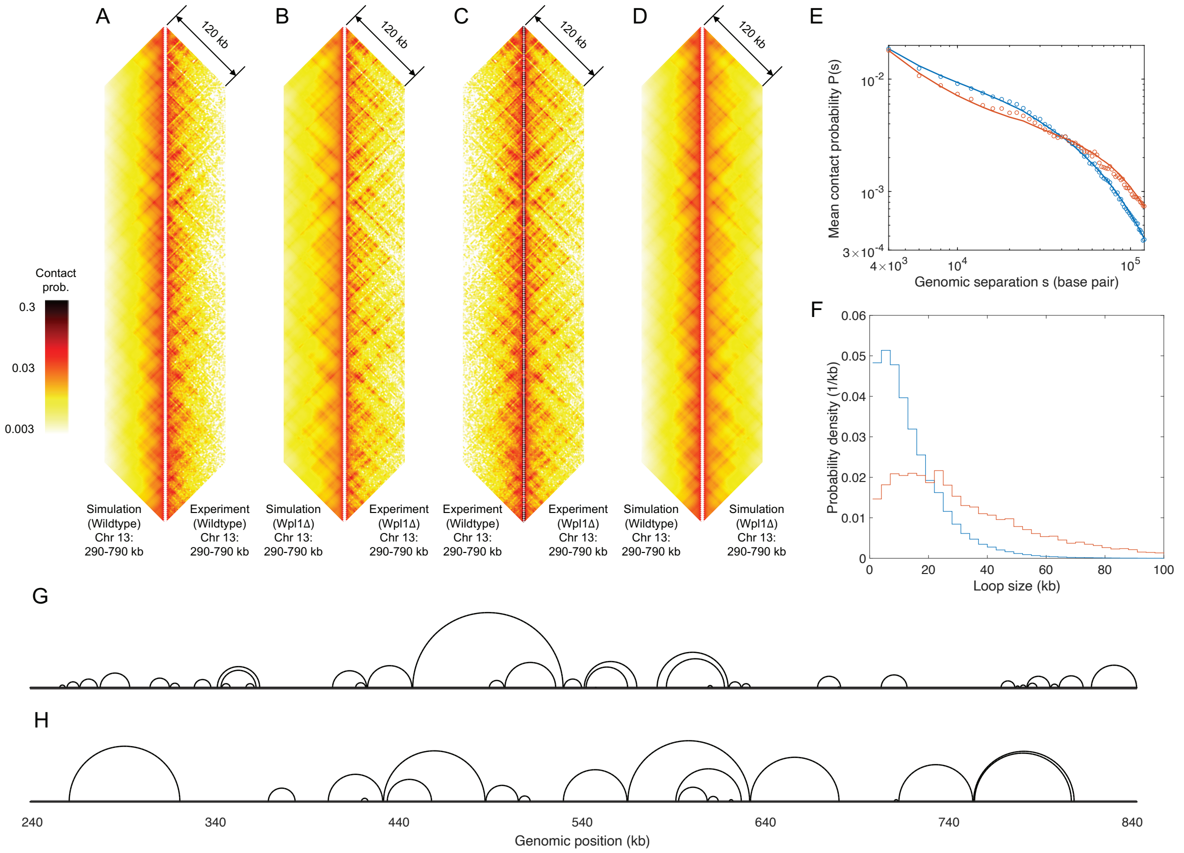


Supplementary Figure 17: Conserved-current loop extrusion (CCLE) model recapitulates TAD-scale chromatin organization in meiotic *S. cerevisiae* lacking Wpl1. Comparison between (A) CCLE-simulated and experimental Hi-C maps of wild-type cells, (B) CCLE-simulated and experimental Hi-C maps of Wpl1-depleted cells, (C) experimental Hi-C maps of wild-type and Wpl1-depleted cells, and (D) simulated Hi-C maps of wild-type and Wpl1-depleted cells, for the genomic region of 290-790 kb of Chr 13. (E) Chromatin mean contact probability, $P(s)$, plotted as a function of genomic separation, $s$, for the experimental Hi-C maps of wild-type (blue circles) and Wpl1-depleted (red circles) cells, and for the corresponding simulated Hi-C maps of wild-type (blue line) and Wpl1-depleted (red line) cells. (F) Distributions of loop size in wild-type (blue) and Wpl1-depleted (red) cells, generated by CCLE for the 240–840 kb region of Chr 13 of meiotic *S. cerevisiae*. Means and standard deviation are 13.48 and 11.33 kb, for wild-type cells, and 33.70 and 27.81 kb, for Wpl1-depleted cells, respectively. Snapshots of representative simulated meiotic loop configurations in (G) wild-type and (H) Wpl1-depleted cells, for the 240–840 kb region of Chr 13. In each case, the chromatin backbone is represented as a straight line, while loops are represented as semicircles connecting loop anchors, following Ref. [5]. Simulation results shown in this figure are generated by CCLE using the best-fit parameters. For wild-type cells, LEF density ($\rho$) is 0.063 kb$^{-1}$; mean processivity ($L$) is 18.95 kb; persistence length is 160 nm; standard deviation of the Gaussian scaling factor is 90 kb. For Wpl1-depleted cells, LEF density is 0.032 kb$^{-1}$; mean processivity ($L$) is 55.58 kb; persistence length is 150 nm; standard deviation of the Gaussian scaling factor is 92 kb. Because the ChIP-seq of both wild-type and Wpl1-depleted meiotic *S. cerevisiae* cells has minimal background signal, the parameters of cohesive cohesin density are set to zero in both cases.

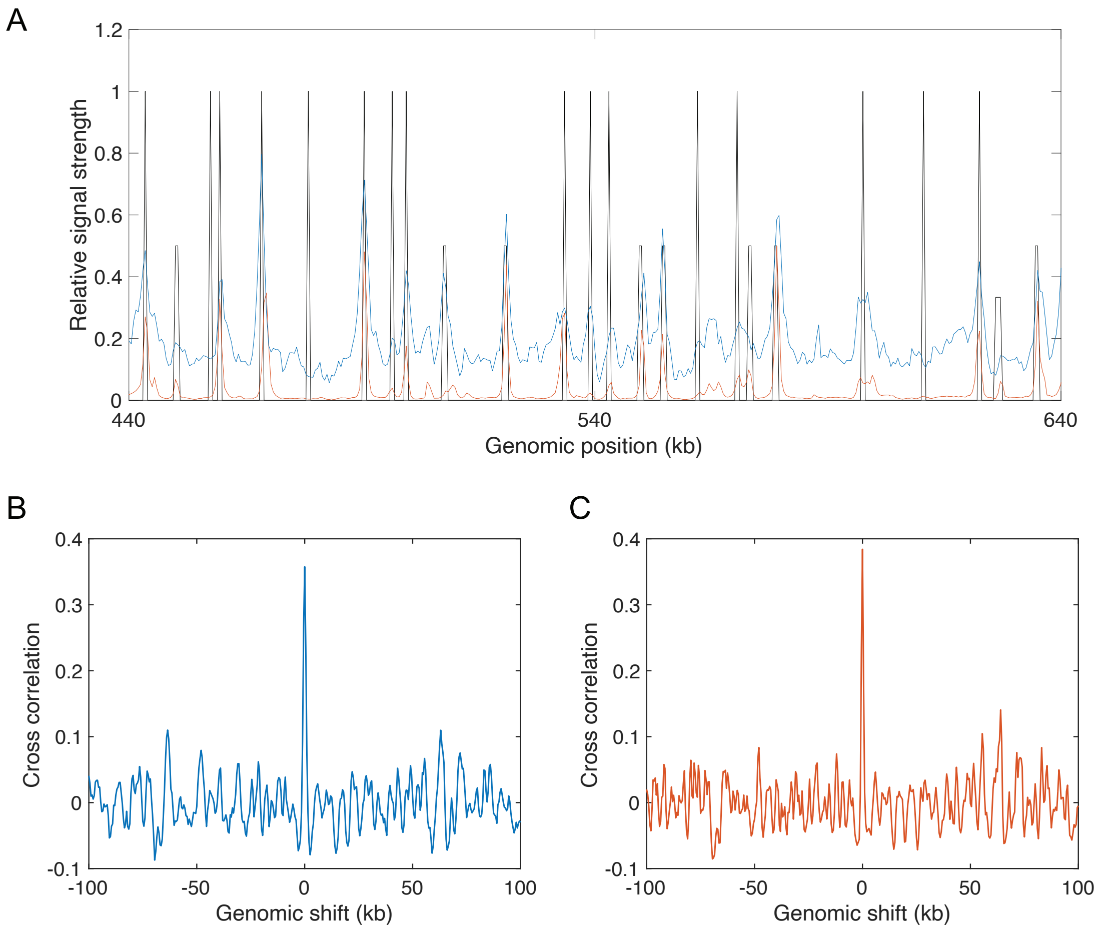


Supplementary Figure 18: Meiotic and mitotic cohesin share the same peak positions and are both correlated to positions of convergent gene pairs in *S. cerevisiae* . (A) Meiotic (blue) and mitotic (red) cohesin ChIP-seq plotted alongside the convergent gene variable (black) for the 440–640 kb region of Chr 13 of *S. cerevisiae*. The convergent gene variable is defined such that it has a non-zero value between convergent genes and an integrated weight of unity for each convergent gene pair. (B) Cross-correlation between meiotic cohesin ChIP-seq and the convergent gene variable. (C) Cross-correlation between mitotic cohesin ChIP-seq and the convergent gene variable.

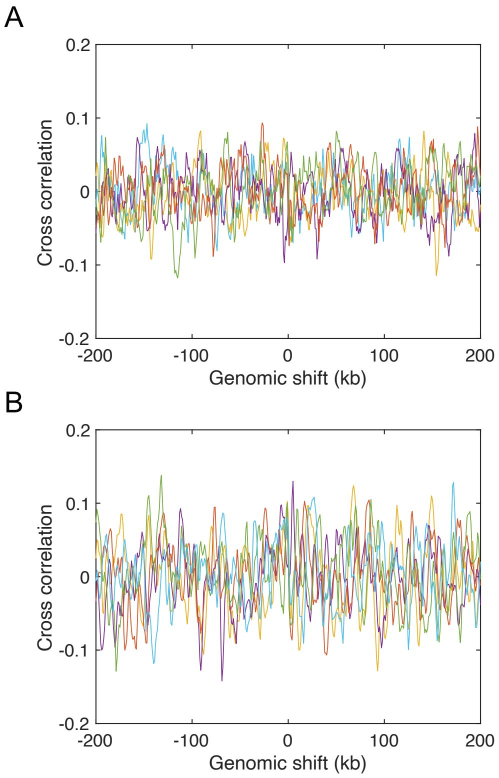


Supplementary Figure 19: (A) Cross-correlations between cohesin ChIP-seq signal (Psc3 [2]) and the convergent gene variable in interphase *S. pombe*, for the five different genomic regions. Yellow, Chr 1: 0.5–1.7 Mb; purple, Chr 1: 4.2–5.4 Mb; red, Chr 2: 0.2–1.4 Mb; green, Chr 2: 1.8–3.0 Mb; cyan, Chr 3: 1.2–2.4 Mb. (B) Cross-correlations between cohesin and Pol II ChIP-seq in interphase *S. pombe*, for the same five genomic regions specified above.

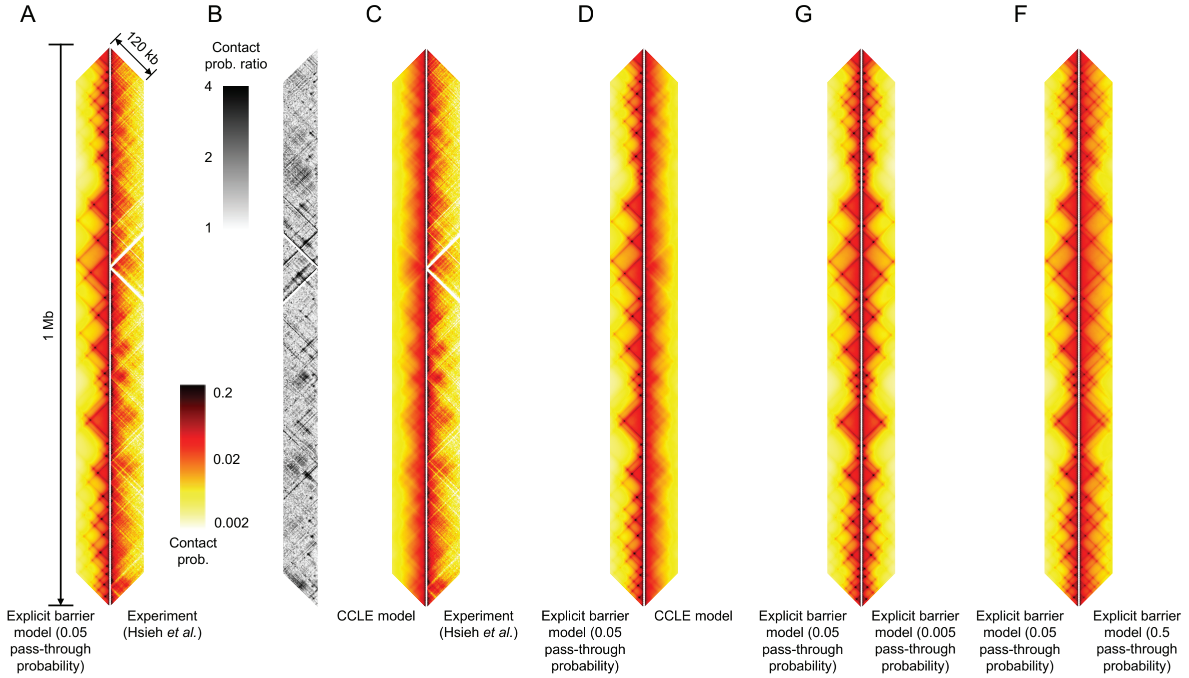


Supplementary Figure 20: Explicit barrier model poorly describes TAD-scale chromatin organization in the 0.3-1.3 Mb region of Chr 2 of interphase *S. pombe*. (A) Comparison between simulated Hi-C map of 1 Mb region generated by the explicit barrier model (using peaks of interphase Psc3 ChIP-seq [2] to define loop extrusion barriers) and the experimental Hi-C map [3] of the same region, binned to 2 kb resolution. The barrier pass-through probability is set to 0.05, according to the best-fit value given by Ref. [6]. Both Hi-C maps show interactions up to genomic separation of 120 kb. The MPR and PCC scores for this comparison are 1.6489 and 0.2267. (B) Contact probability ratio map between the Hi-C maps shown in panel (A). (C) Comparison between the CCLE-simulated and experimental Hi-C maps from Ref. [3], at 2 kb resolution. (D) Comparison between simulated Hi-C maps generated by the CCLE and the explicit barrier model, using barrier pass-through probability of 0.05. (E) Comparison between simulated Hi-C maps generated by the the explicit barrier model using pass-through probability of 0.05 and 0.005. (F) Comparison between simulated Hi-C maps generated by the the explicit barrier model using pass-through probability of 0.05 and 0.5.

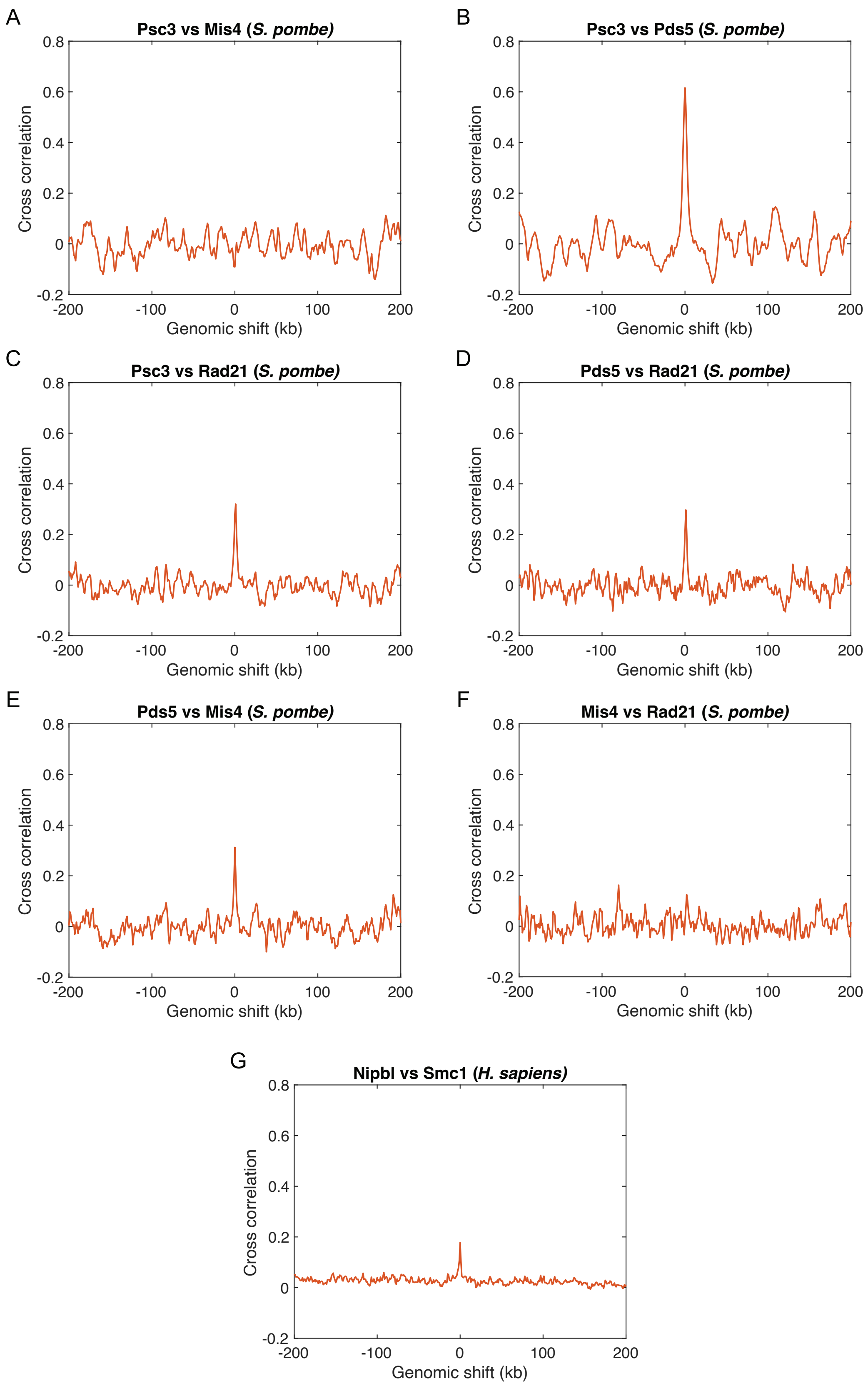


Supplementary Figure 21: (A)–(F) Cross-correlations between different pairs of the ChIP-seq data for Psc3 [2], Rad21 [4], Mis4 [7], and Pds5 [7] of interphase *S. pombe*, as a function of relative genomic shift between each pair of data in comparison. ChIP-seq data of the 0.2–1.4 Mb region of interphase *S. pombe*'s Chr 2 is used. (G) Cross-correlation between the ChIP-seq of Nipbl and Smc1 of *H. sapiens* [8], for the region 66–78 Mb of Chr 5. Both regions considered in *S. pombe* and *H. sapiens* are non-centromeric.

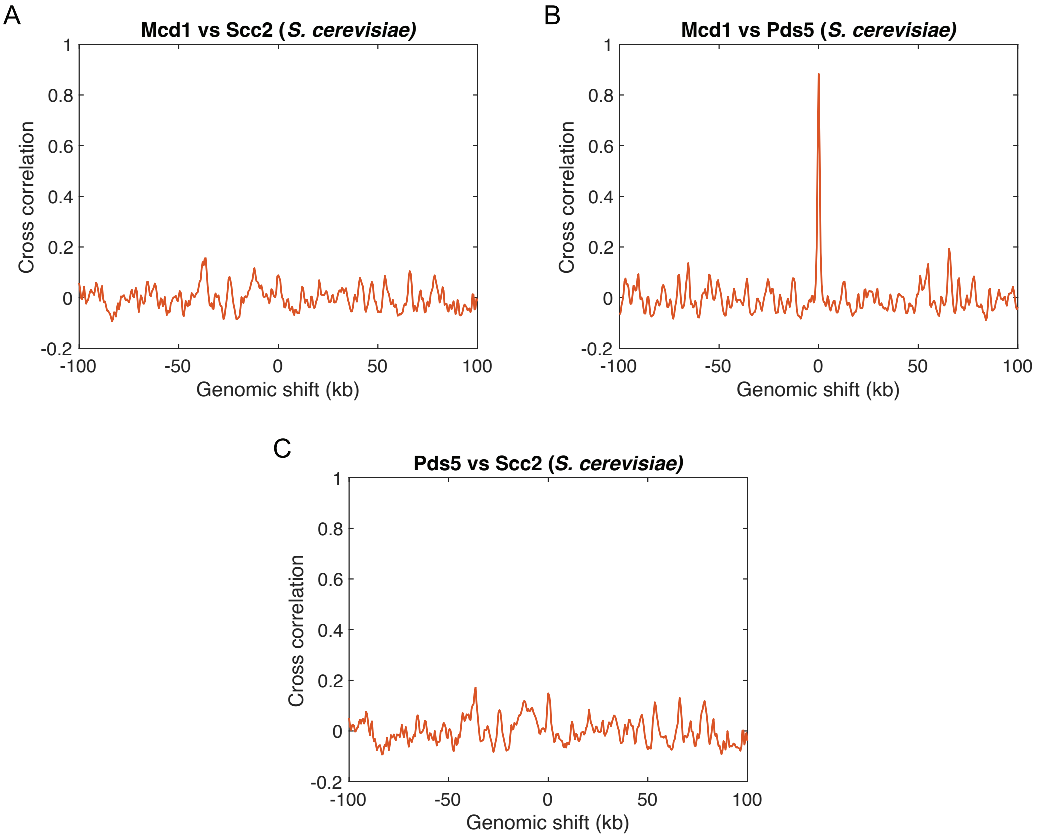


Supplementary Figure 22: Cross-correlations between different pairs of the ChIP-seq data for Mcd1 [9], Pds5 [10], and Scc2 [11] of mitotic *S. cerevisiae*, as a function of relative genomic shift between each pair of data in comparison. ChIP-seq data of 0.55–1.05 Mb (non-centromeric) region of mitotic *S. cerevisiae*'s Chr 7 is used.

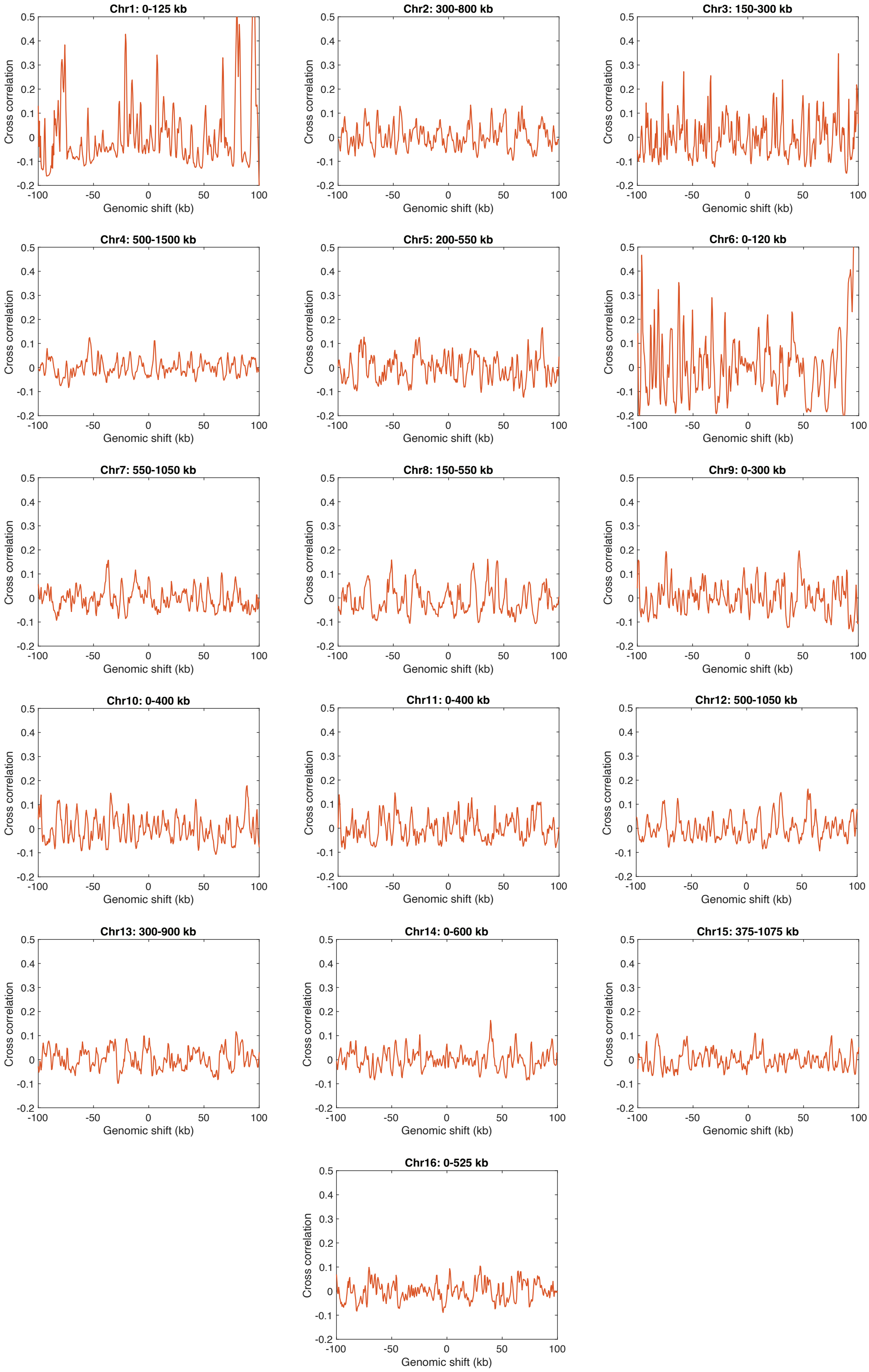


Supplementary Figure 23: Cross-correlations between the ChIP-seq data of Mcd1 [9] and Scc2 [11] in non-centromeric regions of all chromosomes in mitotic *S. cerevisiae*, as a function of relative genomic shift between each pair of data in comparison. The title above each panel indicates the non-centromeric region for which the cross-correlation is calculated.

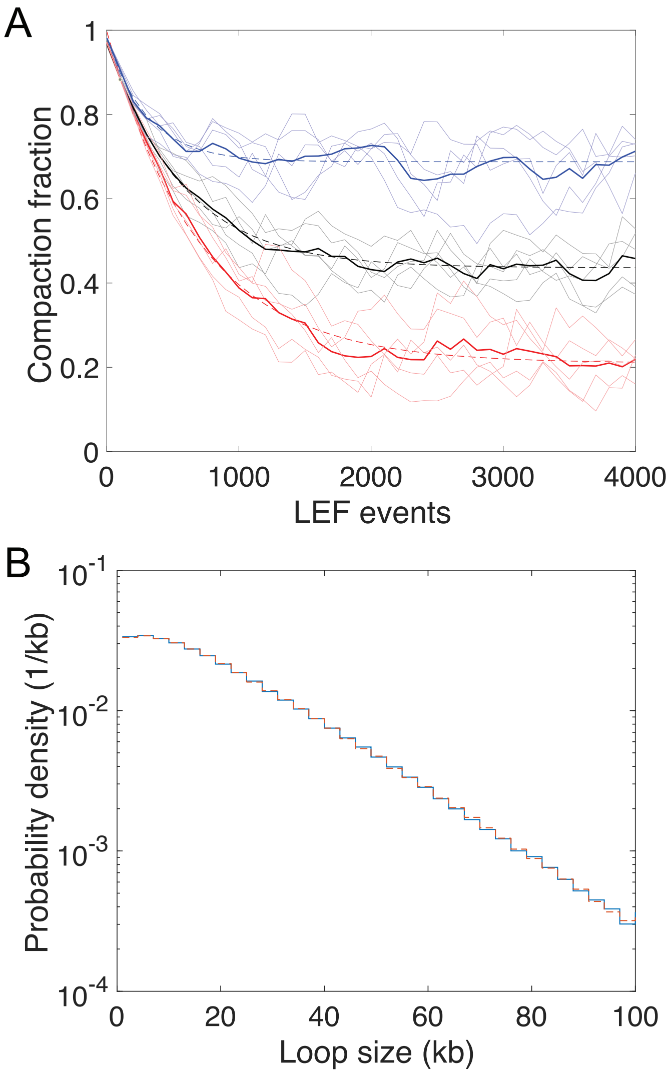


Supplementary Figure 24: LEF simulations reach dynamic steady state well within 15000 LEF events. (A) Time evolution of polymer compaction by loops in interphase *S. pombe* (black), meiotic *S. cerevisiae* (red) and mitotic *S. cerevisiae* (blue). Each solid thin line represents the data from an independent LEF simulation. Each solid thick line is the average value of the corresponding five independent simulations. The dashed lines are the exponential fits of the averages. Compaction is defined as the ratio of mean-squared radius of gyration of a looped polymer to that of a Gaussian polymer without loop, as described in more detail in Ref. [12]. (B) Loop size distributions from two different simulation periods of the same CCLE simulation of interphase *S. pombe*. Each distribution encompasses a set of 1000 data points, equally separated in time, one between LEF event 15000 and 35000 (blue), and the other between LEF event 80000 and 100000 (red).

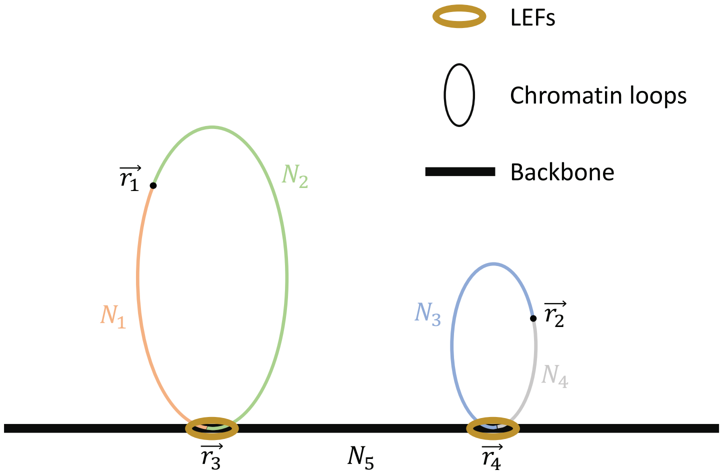


Supplementary Figure 25: A bottlebrush configuration with two loops. $\mathbf{r}_1$ and $\mathbf{r}_2$ are the coordinates of the points of interest in different loops. $\mathbf{r}_3$ and $\mathbf{r}_4$ are the coordinates of the two loop bases. $N_1$, $N_2$, $N_3$, $N_4$, and $N_5$ are the contour distances of the corresponding genomic segments.
